# Circadian dynamics of the teleost skin immune-microbiome interface

**DOI:** 10.1101/2021.01.29.428758

**Authors:** Amy R Ellison, David Wilcockson, Jo Cable

**Author notes:** Joint senior authors, contributed equally (David Wilcockson, Jo Cable).

## Abstract

Circadian rhythms of host immune activity and their microbiomes are likely pivotal to health and disease resistance. The integration of chronotherapeutic approaches to disease mitigation in managed animals, however, is yet to be realised. In aquaculture, light manipulation is commonly used to enhance growth and control reproduction but may have unknown negative consequences for animal health. Infectious diseases are a major barrier to sustainable aquaculture and understanding the circadian dynamics of fish immunity and crosstalk with the microbiome is urgently needed. We demonstrate daily rhythms in fish skin immune expression and microbiomes, that are modulated by photoperiod and parasitic infection. We identify putative associations of host clock and immune gene profiles with microbial composition. Our results suggest circadian perturbation that shifts the magnitude and timing of immune and microbiota activity, is detrimental to fish health. This study represents a valuable foundation for investigating the utility of chronotherapies in aquaculture, and more broadly contributes to our understanding of circadian health in vertebrates.

## Introduction

Circadian rhythms – endogenous daily cycles in physiological and behavioural processes – are a ubiquitous phenomenon to life. Living organisms are adapted to anticipate the daily variations in light, temperature, or food availability driven by the relentless 24 h rotation of Earth. Circadian rhythms are orchestrated by so-called “clock genes” driving transcriptional-translational autoregulatory feedback loops^1^, which are transduced to temporally coordinate biological activities. Immune functions are energetically costly^2^ and often highly rhythmic, enabling organisms to mount their most efficient response at times when risk of infection or injury are highest^3–5^. Conversely, immune factors and infections can affect expression of molecular clocks^6–8^ and subsequent rhythmic phenotypes^9,10^. Disruption of normal circadian cycles can impact immune functioning^11,12^ and may increase disease risks^13^.

A primary function of immune systems is to protect the host from invading pathogenic microbes. However, animals are invariably colonized by a suite of microorganisms – their “microbiome” - which span the spectrum of symbiosis from mutualists to opportunistic pathogens. In vertebrates, it is increasingly apparent that immune systems and microbiomes are intricately linked, together mediating homeostasis and influencing disease outcomes^14,15^. Intriguingly, microbiomes may also be rhythmic, exhibiting diurnal fluctuations in community composition and activity^16^. In studies of the mammalian gut, it has been demonstrated that not only does expression of host clock genes shape microbiome rhythms^17^, but disruption of microbial rhythms in turn impacts host circadian functioning^18^.

Aquaculture is the world’s fastest growing food sector, but infectious disease is the principle barrier to sustainability^19^ and a multi-billion-dollar problem for the global industry^20^. Whilst understanding of fish microbiomes is still in its infancy compared to mammalian systems, there is rapidly growing interest in their role for fish nutrition, health and disease resistance^21–23^. Photoperiod manipulation is commonly used in fish farms, with extended day lengths and, in the extreme, constant light, to promote increased growth rates, or control maturation and reproduction^24–26^. Fish are thought to have a decentralised clock, with cells from multiple tissues expressing circadian genes^27,28^, self-sustained rhythmicity and light responsiveness (see ^29^ for review). In common with higher vertebrates, fish appear to exhibit circadian rhythmicity in certain immune factors^27,28,30–33^. Therefore, extreme lighting regimes may have profound implications for fish health and response to infection. Moreover, there are indications that infection and/or stress may impact expression of fish circadian clocks^34,35^. Currently, the extent to which light manipulation practices contribute to disease in aquaculture is unknown. More fundamentally, the daily dynamics of the fish immune-microbiome interface is yet to be explored. Uncovering the effects of infection and photoperiod on fish immune and microbiome rhythms will be pivotal for both aquaculture disease mitigation strategies, and a broader understanding of the role of holobiont chronobiological interactions for animal health.

Here, using rainbow trout (*Oncorhynchus mykiss*) as a model, we combine 16S rRNA metabarcoding and direct mRNA quantification methods to simultaneously characterise the circadian dynamics of skin clock and immune gene expression, and daily changes of skin microbiota. We compare circadian rhythms of host clock and immune gene expression and microbial community composition in healthy fish under regular light-dark cycles (12:12 LD) with those in fish experimentally infected with the ectoparasite crustacean *Argulus foliaceus* and/or raised under constant light (24:0 LD, hereafter LL). In addition, we assess rhythmicity in the functional potential of trout skin microbiomes and establish host expression-microbiome association networks.

## Results

### Photoperiod impacts host responses to infection

Photoperiod (12:12 LD vs LL) had no significant impact on growth of juvenile rainbow trout over the 16-week trial period (weight: t_956_ = 0.073, P = 0.942, length: t_956_ = 0.222, P = 0.825, Supplementary Figure 1a & 1b). However, a significantly higher number of *Argulus* lice survived 7 days post-inoculation on fish maintained in constant light conditions (t_115_ = −8.418, P = 1.23 × 10^-23^, Supplementary Figure 1c). To examine overall immune responses to *Argulus* infection, we grouped fish from all timepoints, and contrasted expression of 27 genes from innate and adaptive immune pathways between treatment groups (12:12 LD control, 12:12 LD infected, LL control, LL infected). Infected trout had significantly higher expression of 24 immune genes (89%) under 12:12 LD, whereas only 14 (52%) were significantly higher in infected fish compared to healthy controls under constant light (Figure 1). Two genes (*c3* and *tgfb*) were significantly reduced by infection in both light conditions (Figure 1). Expression levels were broadly similar among infected groups, although upregulation of the pro-inflammatory interleukins *il4* and *il6* was lower under constant light (Figure 1). Conversely, comparisons of healthy (unchallenged) fish under LD and LL revealed a substantial difference in immune expression profiles, with unchallenged fish under constant light exhibiting elevated expression levels in 21 genes (78%), more similar to both infected groups in most immune genes (Figure 1).

**Figure 1:**
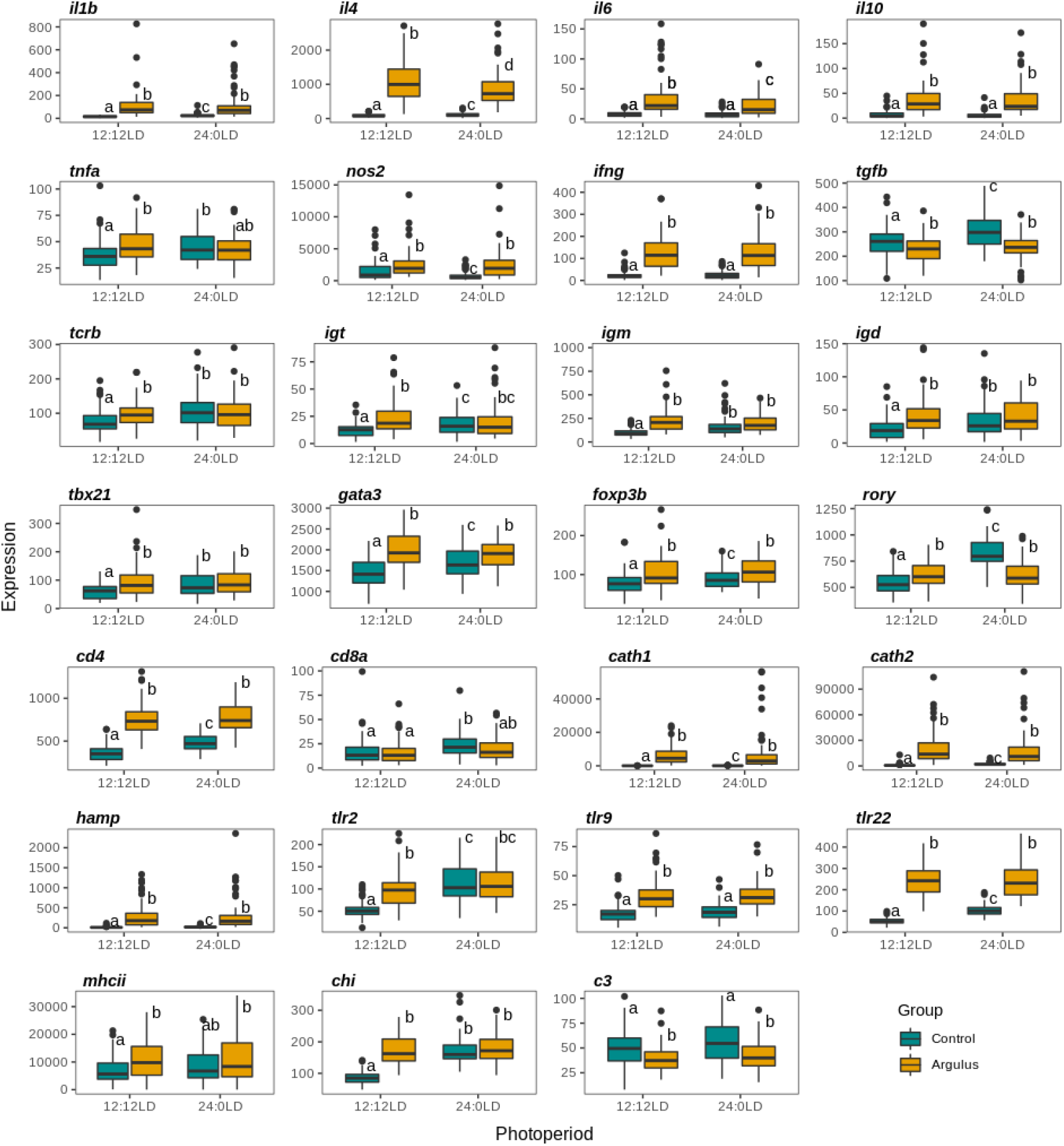
Expression of immune genes in uninfected (control; cyan) and *Argulus*-infected (orange) rainbow trout maintained under 12:12 LD and 24:0 LD conditions. Letters denote significant differences in expression between groups. Expression is normalised counts of mRNA copies detected via Nanostring nCounter.

### Circadian rhythmicity of host expression is altered by infection and photoperiod

Under 12:12 LD, core and accessory vertebrate clock genes exhibited significant circadian rhythmicity in healthy trout skin (Figure 2, Table 1, Supplementary Figure 2). Many of these genes are also found to be expressed rhythmically in fish raised in constant light (Figure 2, Table 1, Supplementary Figure 2) and when fish are placed into “free-running” (DD) conditions (Supplementary Figure 3, Table 1). However, overall expression levels of clock genes are elevated in the absence of light cues (Figure 2, Supplementary Figure 2), except for *timeless* (suppressed expression in LL). In addition, *bmal2, clock1b, per1*, and *rora* exhibited a significantly different phase of expression in constant light (Table 1, Figure 2, Supplementary Figure 2).

**Figure 2:**
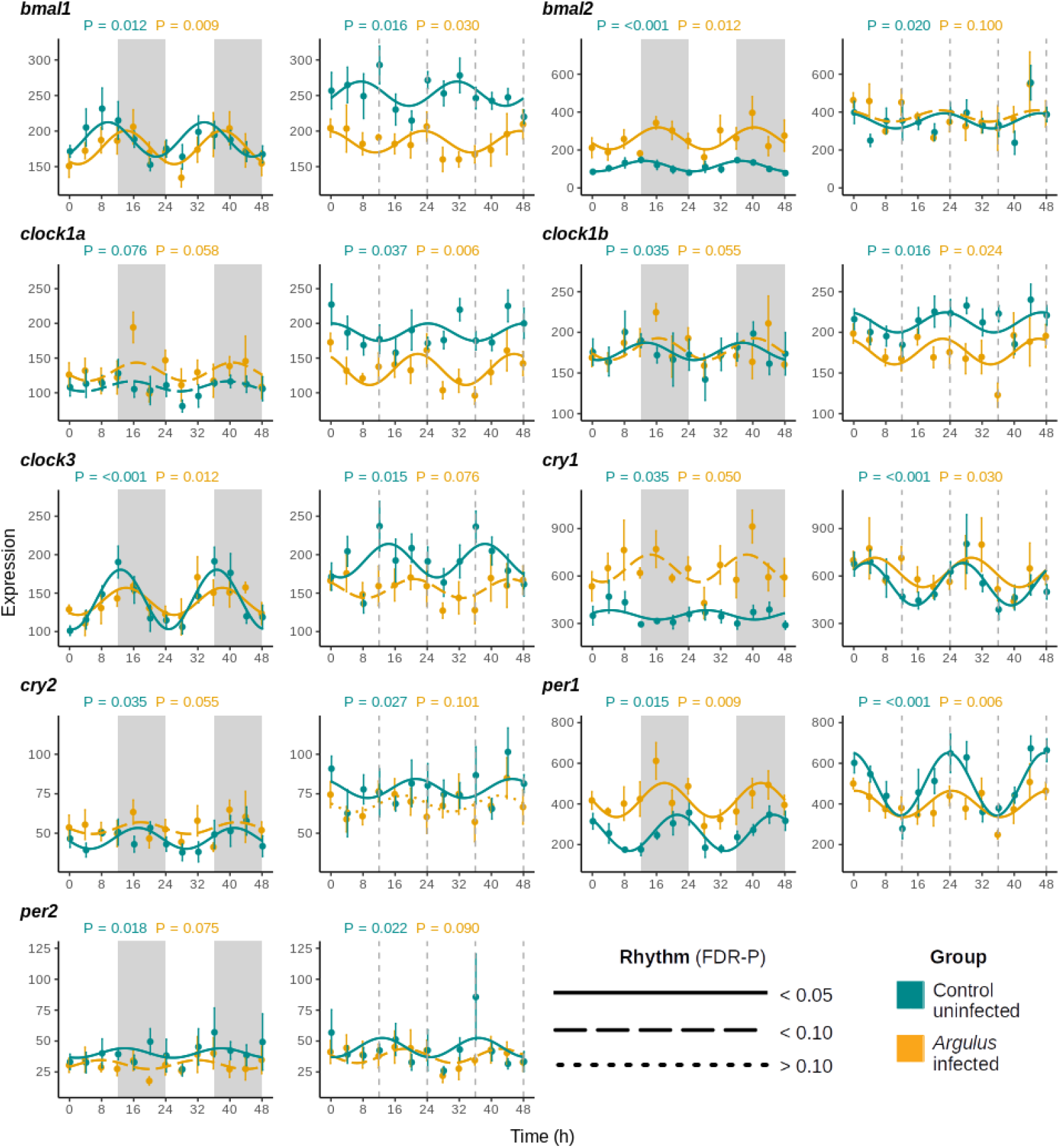
Mean expression (± 1 S.E.) of core clock genes of uninfected (cyan) and *Argulus*-infected (orange) rainbow trout maintained at 12:12 LD (left) and 24:0 LD (LL, right). Expression is normalised counts of mRNA copies detected via Nanostring nCounter. Curves denote cosinor waveform fitted using CircaCompare. Grey shading indicates time periods in darkness (grey dashing indicates equivalent 12:12 LD light transitions on LL plots).

**Table 1:**
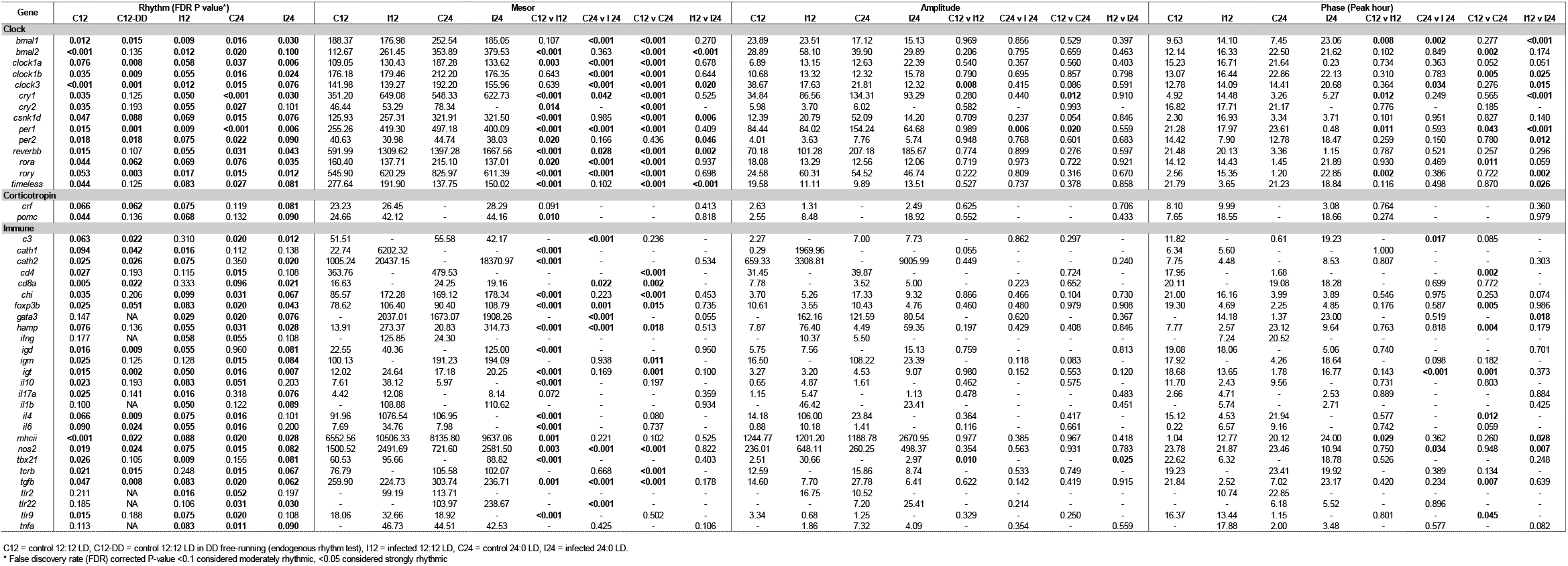
Summary of gene expression rhythmic analyses. Rhythm significance determined via eJTK_cycle. Rhythm parameters (mesor, amplitude, phase) estimated and contrasted in CircaCompare.

*Argulus* lice infections had variable impacts on the expression levels and rhythmicity of the clock genes. When contrasted with healthy control groups, some gene rhythms were dampened in infected fish (i.e. significantly reduced amplitude; 12:12 LD *clock3*, LL per*1*), rendered arrhythmic (*cry2* in LL), and/or phase-shifted (*bmal1* in both light treatments, *cry1* and *per1* in 12:12 LD, *clock3* in LL). Rhythms of clock gene expression in infected fish under the two photoperiod treatments did not differ in amplitude. But, *bmal1, clock1b, clock3, cry1, per1, per2, rory* and *timeless* had significantly different phases of expression between infected fish under 12:12 LD and those raised in constant light. In addition, *bmal2, clock3, csnk1d, per2, reverbb* had increased rhythm mesors in LL, whilst *timeless* was suppressed (Table 1, Figure 2, Supplementary Figure 2).

Significant rhythmicity in expression was found in both innate and adaptive immune markers (Table 1, Supplementary Figures 4 & 5), with a substantial proportion remaining rhythmic under free-running (DD) conditions (Supplementary Figure 6). The cathelicidins (*cath1, cath2), igd, il17a*, and *tbx21*, while rhythmic in healthy fish under 12:12 LD, were arrhythmic in fish maintained in constant light (Table 1, Supplementary Figures 4 & 5). Of the immune genes rhythmic in healthy fish under both light conditions, the innate markers *chi, hamp* and *nos2*, and the adaptive markers *cd4, cd8a, foxp3b, igm, igt, tcrb* and *tgfb* had significantly different mesors; with the exception of *nos2*, all were more highly expressed in LL. However, some of these more highly expressed genes (*cd4, foxp3b, hamp, igt, tgfb*) and others with similar expression levels between photoperiods (*il4, tlr9*), were phase-shifted in constant light (Table 1, Supplementary Figures 4 & 5).

Fewer immune genes were rhythmically expressed in infected fish: 76% and 67% of rhythmic genes found in healthy fish were also rhythmic in the 12:12 LD and LL infected groups respectively. Under 12:12 LD, the vast majority (94%) of the immune genes assayed with rhythmicity in both healthy and infected fish exhibited higher mesors in the infected group. In contrast, only 57% of immune genes with rhythms in healthy and infected fish in LL had different expression levels (Table 1). Only *tbx21* had a significantly altered amplitude in rhythm; with a higher amplitude in infected fish at 12:12 LD compared to both healthy 12:12 LD fish and infected fish in constant light. *Argulus* infection also shifted the phase of expression of *mhcii* under 12:12 LD and *c3, nos2* and *igt* in LL (Table 1).

### Argulus infection impacts skin mucus microbiome communities

After read pre-processing, error correction, chimera removal, and filtering, a total of 1,037 amplified sequence variants (ASVs) were found across all samples. Rarefaction curves confirmed a minimum read depth of 2,000 was sufficient to reach saturation of diversity in trout skin (Supplementary Figure 7a). Background water samples were distinct from fish groups (Supplementary Figure 7b) and had a significantly higher alpha diversity (Supplementary Figure 7c). Contrasts of alpha diversity among fish samples revealed that the microbiomes of healthy fish under constant light were significantly less diverse than all other groups (Faith’s PD, all pairwise Kruskal-Wallis tests P<0.001, Supplementary Table 2). Multivariate permutational analysis of beta diversity indicated significant compositional differences among all groups (Supplementary Figure 7b, Supplementary Table 3).

The skin microbiome communities in all groups were dominated by *Proteobacteria*, with *Pseudomonadaceae* and *Burkholderiaceae* accounting for over 50% of the communities in all groups and timepoints (Figure 3). Wilcox rank-sum testing and DESeq2 both revealed substantial differences in the relative abundances of microbial taxa between healthy and lice-infected fish (Figure 4). At the higher taxonomic levels, healthy fish under both light treatments had a greater proportion of *Actinobacteria* and *Firmicutes* lineages, whilst both infected fish groups had increased *Bacterodia* lineages (Figure 4a). At the genus level, many *Gammaproteobacteria* were more abundant in both infected groups (e.g. *Aeromonas, Perlucidibaca, Undibacterium*, Figure 4b). *Bacteroidia* genera, including several *Chryseobacterium, Flectobacillus* and *Flavobacterium* ASVs were also increased in infected fish, with *Flavobacterium* accounting for some of the highest fold-changes in abundance (Figure 4b). Full lists of differentially abundant taxa are provided in Supplementary Table 4.

**Figure 3:**
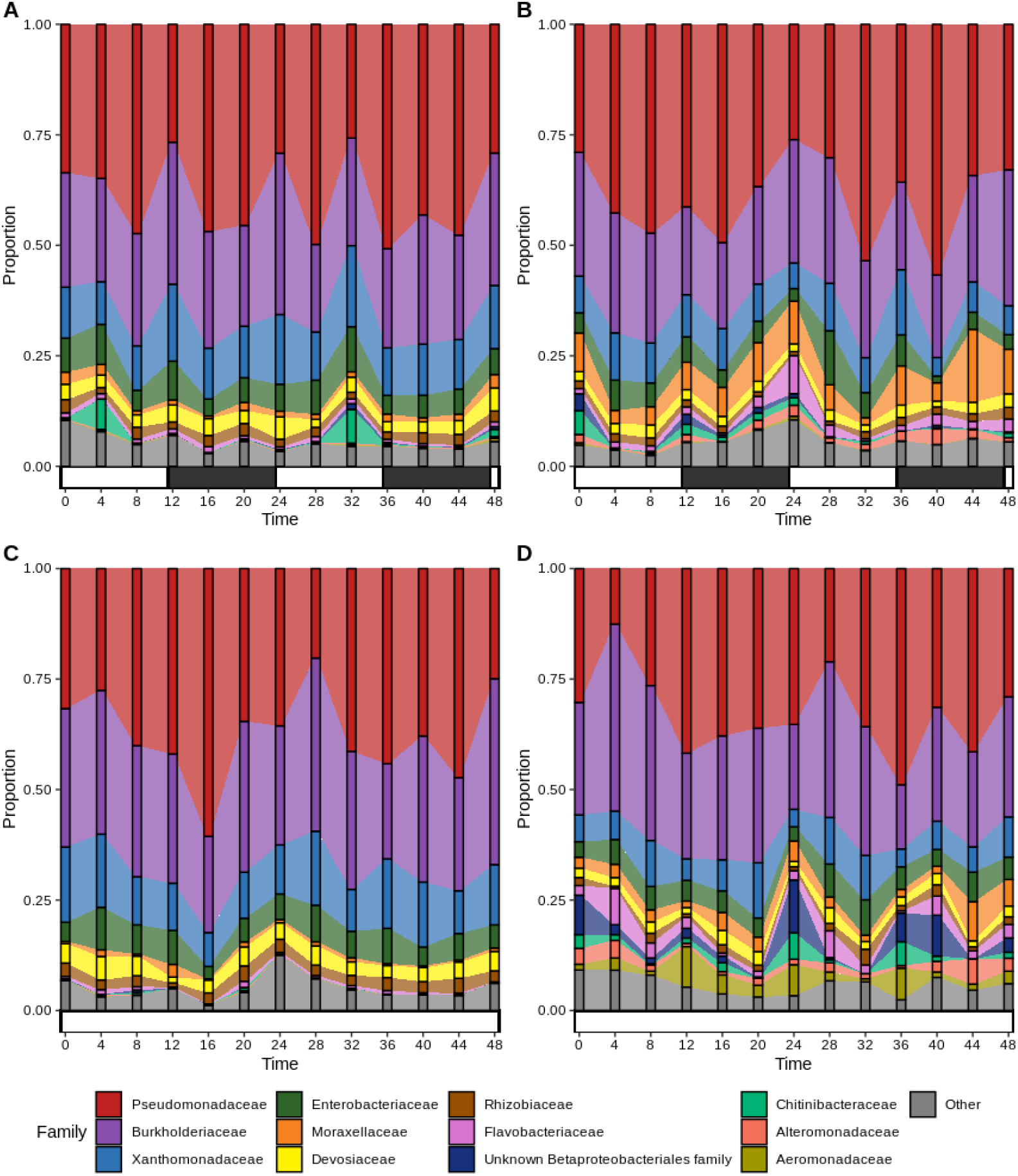
Alluvial plots of most abundant bacteria families (average >1% across all data) in healthy (A, C) and *Argulus foliaceus* infected (B, D) trout under 12:12 LD (A, B) and 24:0 LD (C, D) photoperiods. Horizontal bars indicate periods of light (white) and dark (black).

**Figure 4:**
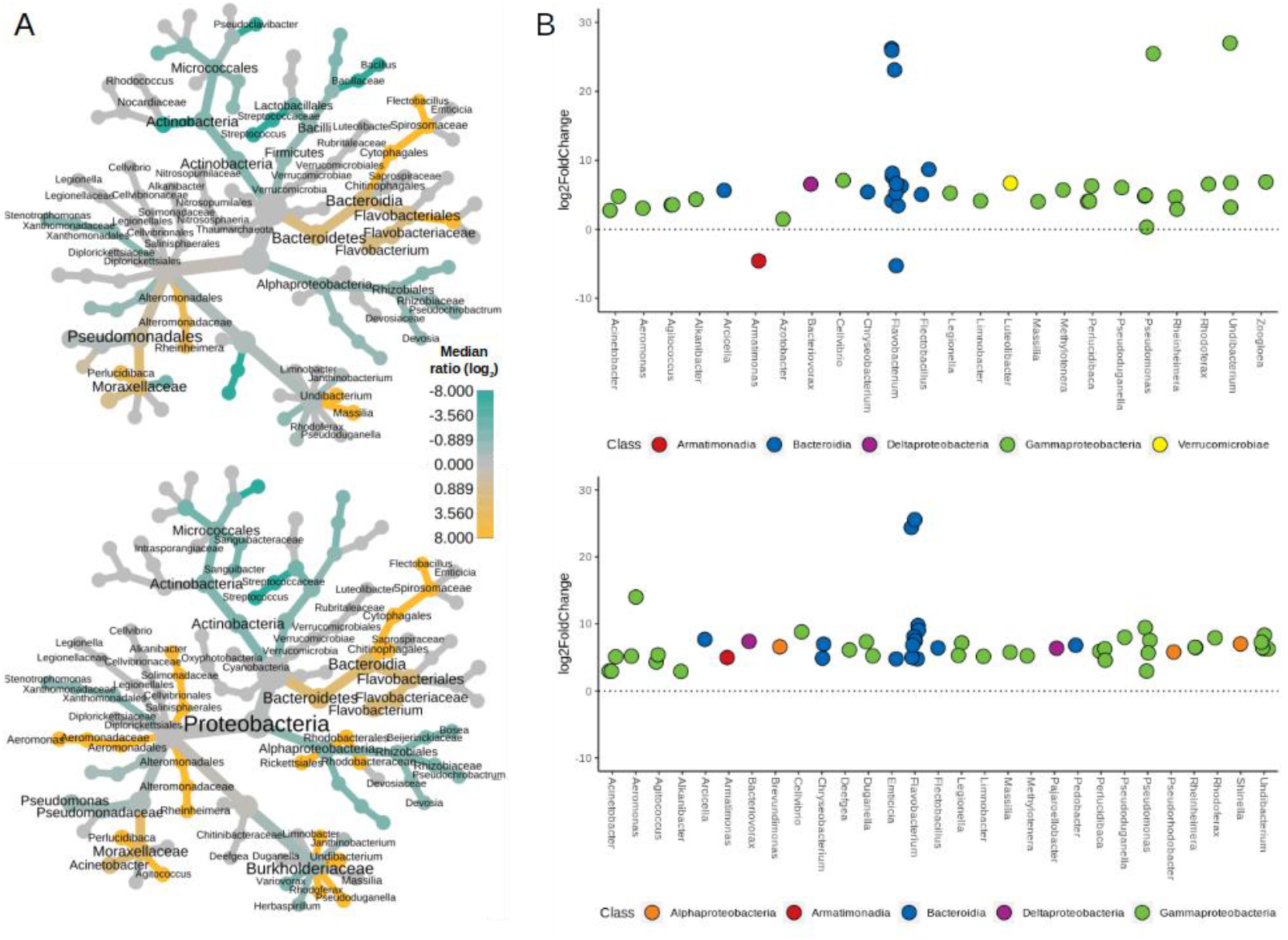
A) Heat trees contrasting bacteria taxa abundance between healthy and *Argulus foliaceus* infected fish under 12:12 LD (top) or 24:0 LD (bottom) photoperiods. The colour of each taxon represents the log-2 ratio of median proportions of reads. Taxa with significant differences are labelled, determined using a Wilcox rank-sum test followed by a Benjamini-Hochberg (FDR) correction for multiple comparisons. Taxa coloured cyan are enriched in healthy fish and those coloured orange are enriched in infected fish. Node size is relative to prevalence in all samples. B) Taxa with significantly different abundances (FDR-corrected p-value <0.05) between healthy and *A. foliaceus* infected fish under 12:12 LD (top) or 24:0 LD (bottom) photoperiods, determined via DESeq2 analyses. Taxa above the dotted line are significantly more abundant in infected fish, below the line are more abundant in healthy fish.

Functional prediction of microbiomes revealed putative differences in the activity of microbial communities among healthy and infected fish. LefSe analyses indicated pathways enriched in healthy fish groups were predominantly degradative classes including amino acid, aromatic compound, and carbohydrate degradation (Table 2). In contrast, functional enrichment of lice-infected fish microbiomes was dominated by biosynthetic pathways in both light conditions, particularly those involved in cofactor, carrier and vitamin biosynthesis (Table 2). Overall, a greater number of pathways were identified as differentially abundant between healthy and infected fish in LL, suggestive of a greater disruption in microbiota functional potential due to parasitic infection in fish maintained under constant light.

**Table 2:**
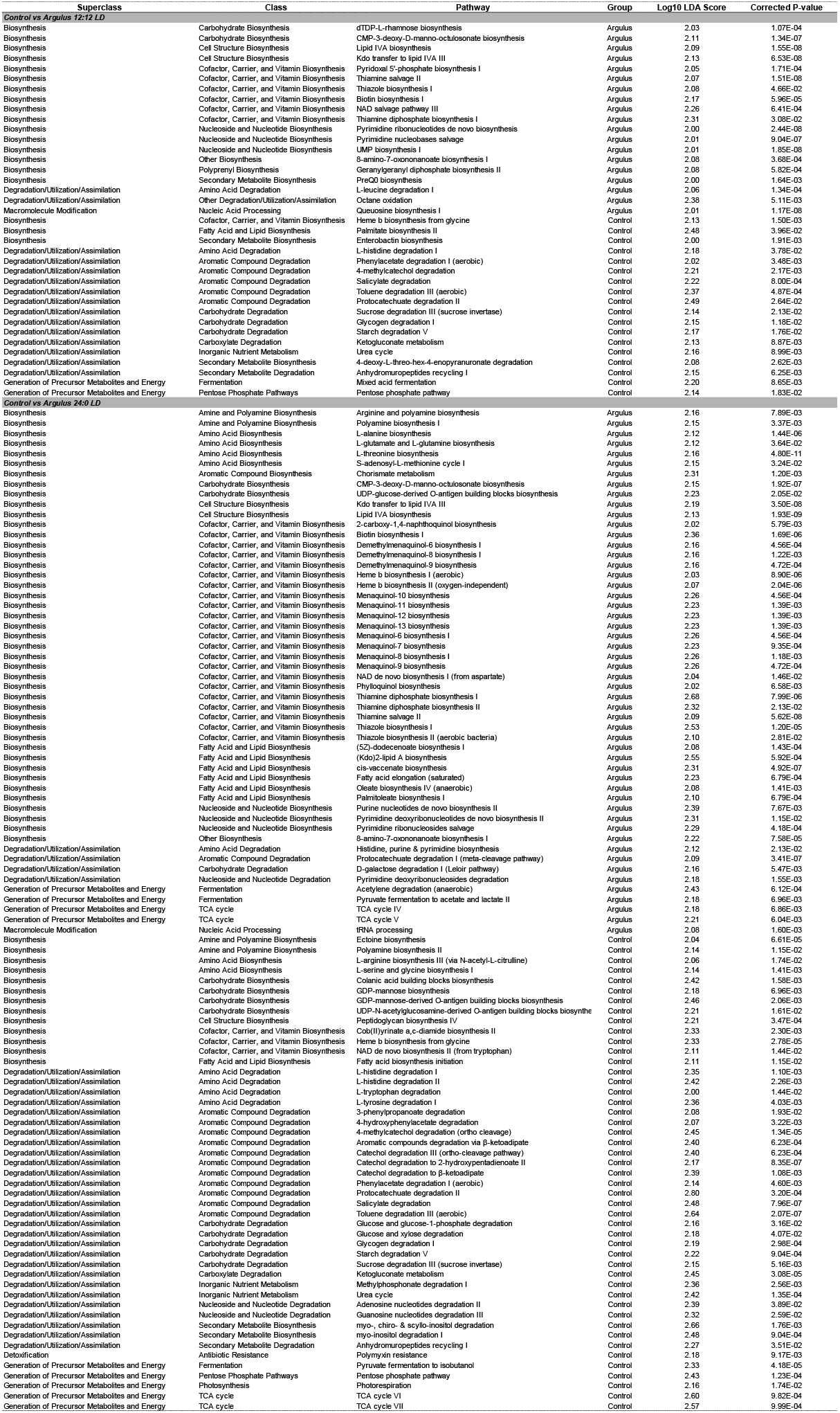
Results of LefSe analyses to identify group differences in the inferred gene abundance of MetaCyc pathways.

### Circadian rhythmicity of skin microbiota and association with host gene expression

Circadian rhythmicity in relative abundance was apparent in 49 skin bacteria genera in one or more of the treatment groups (Table 3, Figure 5). Of the 41 genera rhythmic in both healthy and infected fish at 12:12 LD, 17 (41.5%) had significantly different mesors. In contrast, 60.5% (23/38) had significantly different mesors when comparing healthy and infected fish under constant light. *Perlucidibaca, Undibacterium*, and *Rhodoferax* had significantly greater rhythm amplitudes in infected fish under both light treatments. In addition, *Flectobacillus, Alkanibacter* and an unassigned *Burkholderiaceae* genus had higher rhythm amplitudes in infected 12:12 LD fish, whilst *Duganella* had higher amplitude in LL infected fish only. Under 12:12 LD, lice infection significantly altered rhythm phases of seven bacteria genera (Unknown *Rhizobiaceae*, Unknown *Rickettsiales, Deefgea, Massilia*, Unknown *Neisseriaceae*, Unknown *Chitinophagales* and *Legionella). Pseudoclavibacter* was the only genus found to have altered rhythm phase in LL healthy vs infected comparisons.

**Figure 5:**
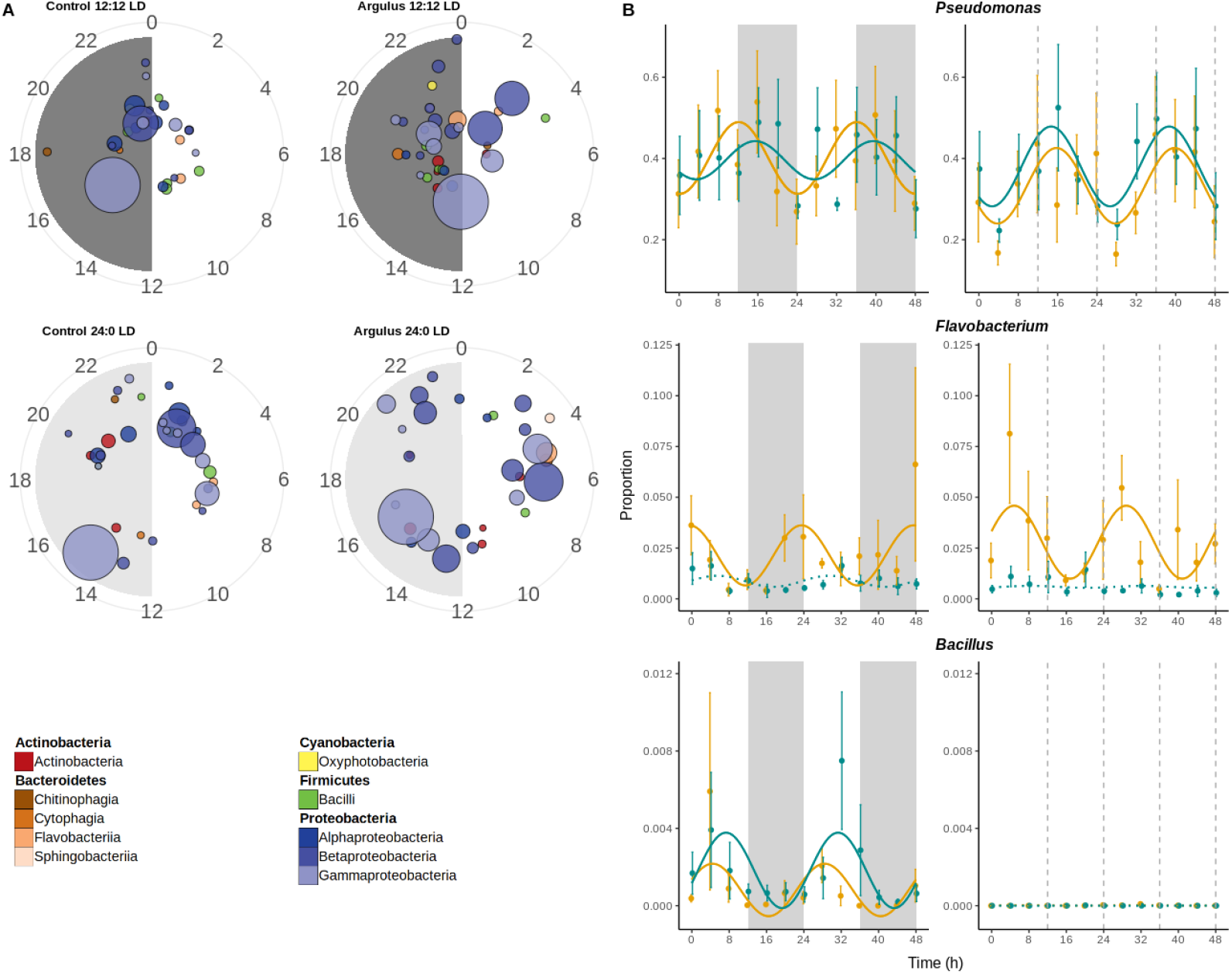
A) Polar plots showing times of peak relative abundance of significantly rhythmic microbiome genera. Each circle represents a genus, coloured by class and scaled by average relative abundance. Radian indicates time of peak and distance from centre indicates significance (more significant/stronger rhythms toward edge of plot). B) Examples of rhythmic bacteria genera (full results presented in Table 3).

**Table 3:**
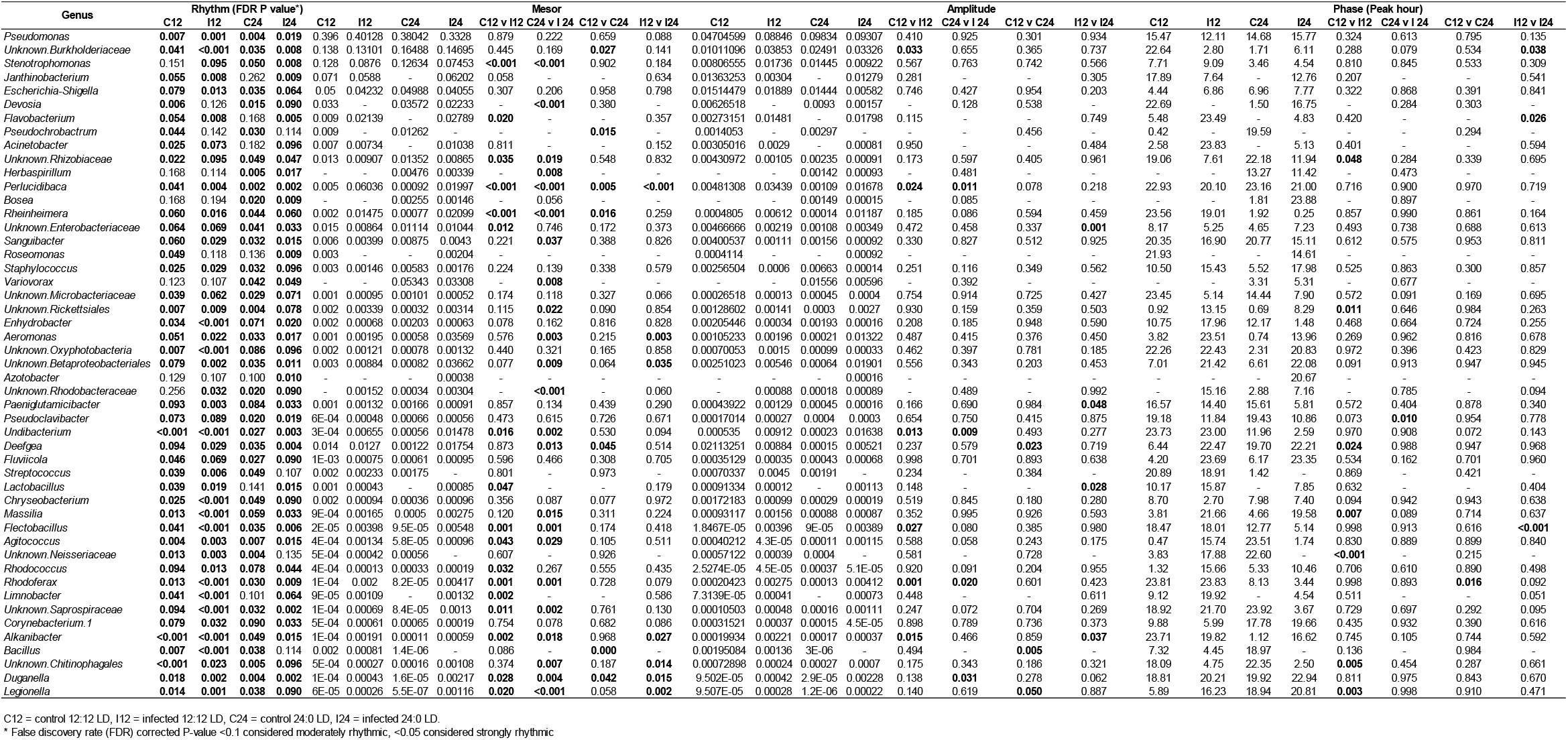
Summary of microbiome rhythmic analyses. Rhythm significance determined via eJTK_cycle. Rhythm parameters (mesor, amplitude, phase of genus relative abundance) estimated and contrasted in CircaCompare.

Visualisation of the timings of peak abundances of rhythmic taxa indicated no clear phylogenetic patterns (e.g. rhythmic *Proteobacteria* genera peak abundances were spread across the circadian cycle, Figure 5a). However, when considering the rhythms of the functional potential of the microbiome communities, we found evidence of temporal patterns (Figure 6). In healthy fish under 12:12 LD, the majority of rhythmic biosynthetic (e.g. heme b, L-lysine and isoprene biosynthesis) and energy generation (e.g. glycolysis, TCA cycle) functions peaked in the first hours of light (ZT0-3), whilst degradation function peaks were found primarily in dark hours (ZT12-21). In contrast, in infected fish under 12:12 LD, rhythmic biosynthetic and energy generation functions predominantly peak in abundance towards the end of the dark period (ZT19-23), whilst degradation pathways peaked just before dark (ZT10-12). Constant light conditions also appeared to shift the broad temporal patterns of function abundances. In healthy fish under LL, many biosynthetic pathways (e.g. L-valine, heme b and enterobactin biosynthesis) peaked at ZT0-3, similar to the 12:12 LD group. However, we also found a large cluster of biosynthetic pathways peaking at ZT14-15 (e.g. fatty acid biosynthesis) and at ZT20-23 (spirillozanthin and coenzyme M biosynthesis). In infected fish under LL, biosynthetic pathway rhythms were more dispersed, with peaks spread around the majority of the 24 h cycle. For degradation and generation of energy pathways in both healthy and infected fish under LL, we found multiple clusters of peak abundances around the 24 h cycle, rather than a single predominant cluster as in 12:12 LD conditions (Figure 6).

**Figure 6:**
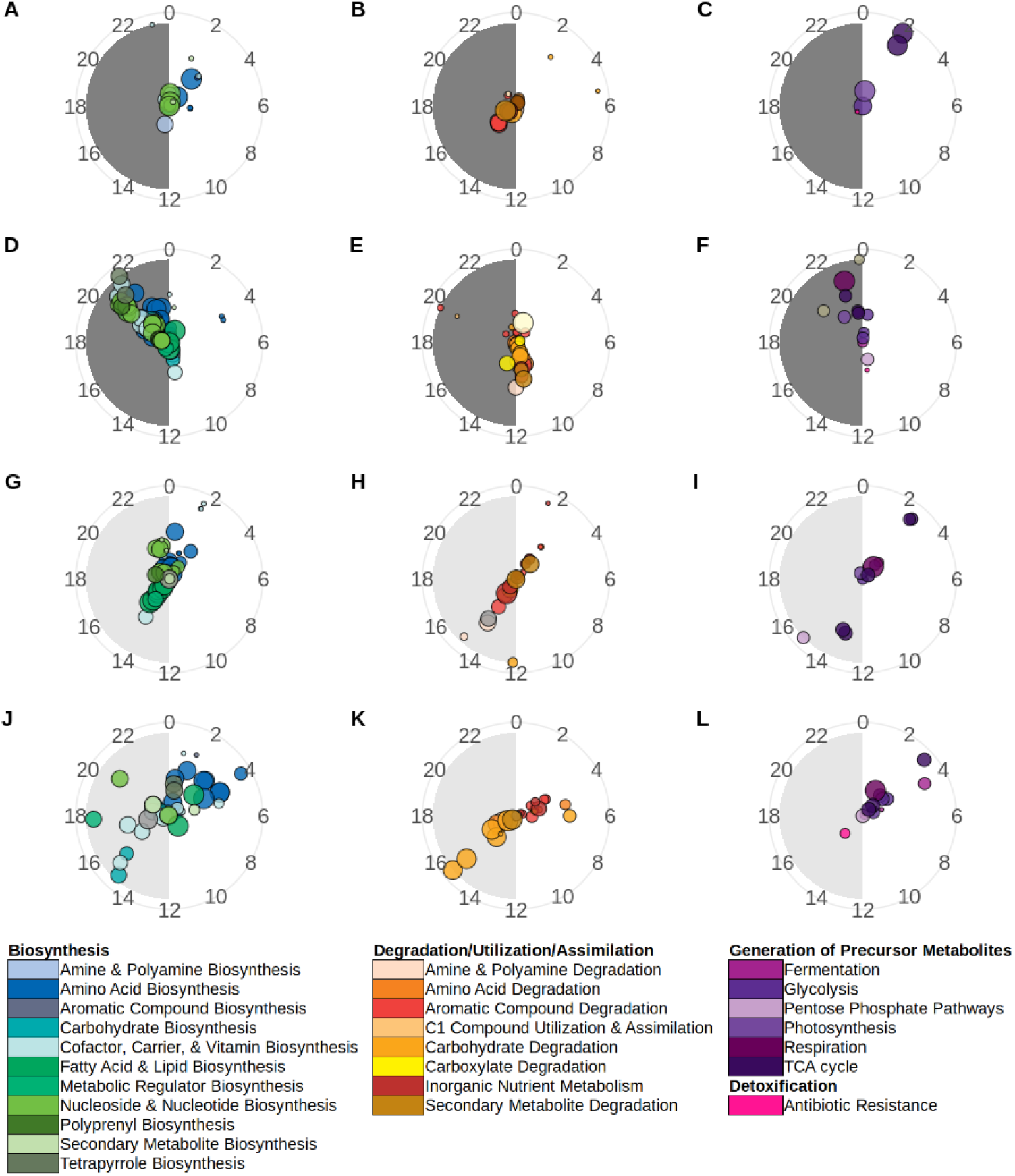
Polar plots showing peak relative abundance of significantly rhythmic microbiome MetaCycle pathways. Each circle represents a pathway, coloured by MetaCycle class and sized by average relative abundance. Pathway radian indicates time of peak and distance from centre indicates significance (more significant/stronger rhythms toward edge of plot). Pathway identity determined via Picrust2 and rhythmicity significance determined via eJTK_cycle (Bonferoni-corrected P-values <0.05). Circacompare was used to fit waveforms and determine estimates of rhythms peaks. A, B, C = healthy trout under 12:12 LD. D, E, F = *Argulus*–infected trout under 12:12 LD. H, I, J = healthy trout under 24:0 LD. K, L, M = *Argulus*-infected trout under 24:0 LD. Full details of pathways are provided in Supplementary Datafile 1.

We used co-occurrence network analyses to assess associations of host gene expression and their microbiomes, using betweeness centrality scores and number of connections (degrees) to identify influential genes and bacteria genera^36,37^. In healthy 12:12 LD fish, there was a high level of connectivity within host immune and clock genes, and within microbial taxa (Figure 7). Links across the gene expression and bacteria subnetworks were primarily via the rhythmically expressed clock genes *clock1b, clock3, bmal1, rora*, and *csnk1d*. However, expression of the toll-like receptors *tlr2* and *tlr9* were significantly associated with abundance of *Bacillus* and *Enhydrobacter* respectively. In contrast, networks of infected fish under 12:12 LD revealed a higher level of connectivity between host expression and bacteria (Figure 7). The immune markers *cd4* and *tcrb*, and the clock gene *reverbb* were found to be most influential in terms of their betweeness centrality scores and number of significantly associated microbial taxa (Figure 7).

**Figure 7:**
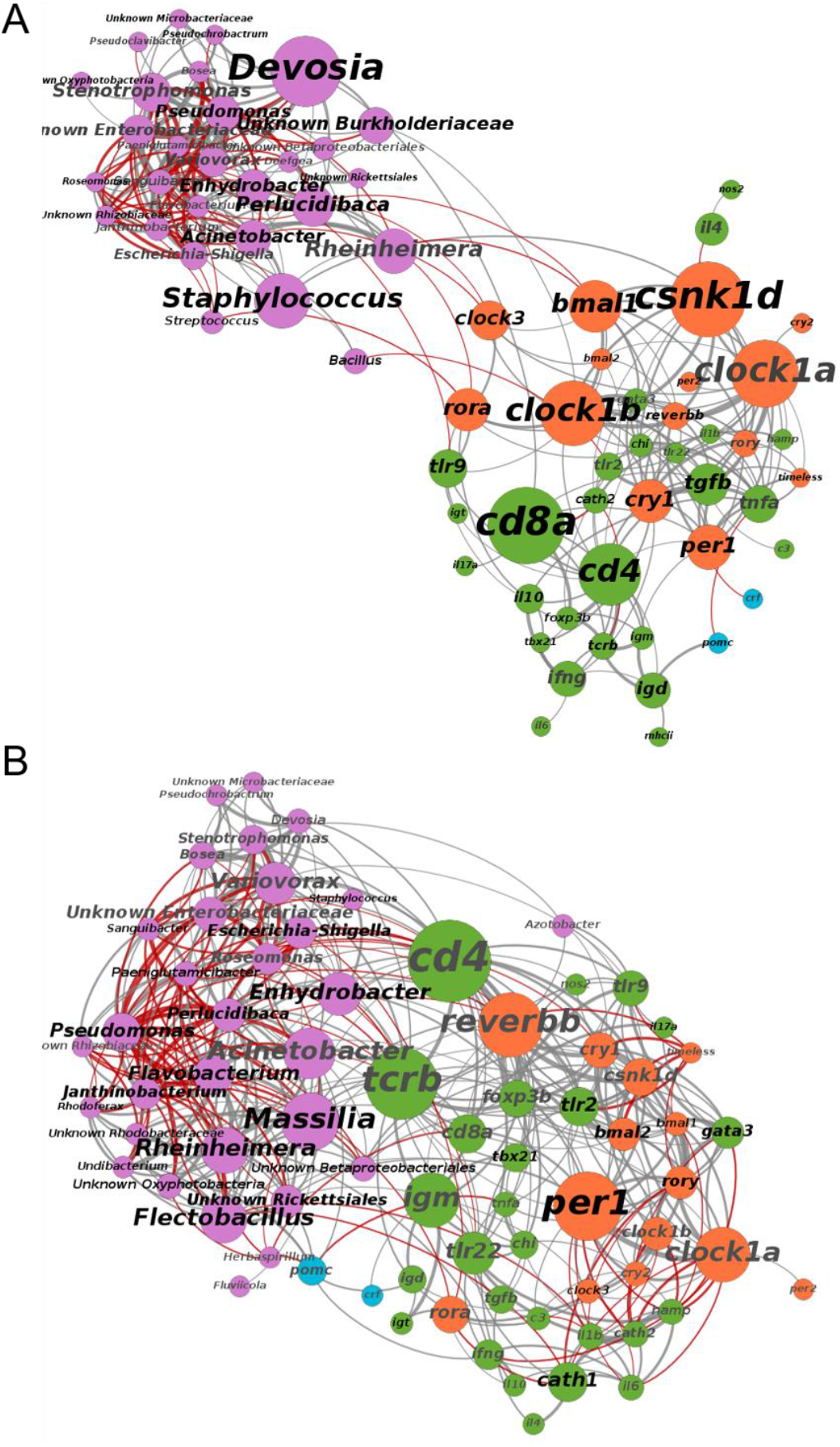
Co-occurrence networks of microbial genera (pink) and host gene expression (orange = clock, green = immune, blue = corticotropin) in healthy (top) and *Argulus*-infected (bottom) trout under 12:12 LD. Node and label size scaled to degree centrality score. Label colour denotes rhythmicity (black = rhythm FDR p-value <0.05, grey = rhythm FDR p-value >0.05). Connection colour indicates association (grey = positive, red = negative, determined by Spearman correlation tests) and connection width scaled to correlation strength (thicker lines denote a higher correlation coefficient).

In contrast to 12:12 LD, clock genes were less influential (in terms of centrality) in gene-microbe networks for uninfected fish under constant light (Supplementary Figure 8). However, several immune genes (*igd, ifng, nos2, hamp, tcrb, foxp3b*) were significantly associated with one or more bacteria genera. *Tcrb* was most influential by betweeness centrality (expression positively correlated with *Janthinobacterium* and negatively with *Flavobacterium*), whilst *ifng* was linked to the highest number of taxa (*Escherichia-Shigella, Pseudomonas, Varioivorax, Stenotrophomonas* and *Pseudoclavibacter*). Similar to 12:12 LD contrasts, the network of infected fish under LL showed a higher level of connectivity between host gene expression and microbiota compared to the healthy network (Supplementary Figure 8), with the immune markers *cd8a* and *tcrb* found to be the most influential genes (in terms of number of associations with taxa and centrality score).

## Discussion

We demonstrate the daily dynamics of immune expression and microbiome composition in fish skin and show ectoparasite infection and constant light – a commonly used environmental condition in aquaculture – can significantly alter circadian rhythms of immunity and microbiota, which may be detrimental for host disease resistance. In addition, we present association networks of host gene expression and their microbiomes, revealing clock expression and T cell populations are likely key in shaping the skin host-microbiome interface of teleosts. Our examination of the skin circadian immune response to infection under extreme photic regimes are directly relevant to fish culture practices; fish peripheral tissues are thought to have entrainable, light-responsive clocks^29^, which may make them particularly susceptible to negative health consequences from constant lighting as used in aquaculture.

Over our trial period, we found no significant difference in the growth of trout fry maintained under 12:12 LD and constant light (LL) when fish were provided equivalent food rations. However, when challenged with *Argulus* lice, their ability to clear infection was significantly altered by photoperiod. Under constant light, trout had a significantly higher lice burden 1 week after inoculation, indicating a reduced ability to mount an effective immune response. These findings are consistent with previous studies showing extended day length increases ectoparasite susceptibility and altered expression in specific immune genes in sticklebacks^38^. Immune profiles in uninfected fish showed elevated levels of expression in both innate and adaptive pathways under constant light. When infected with lice, trout under both photoperiods showed similar patterns of immune gene responses, except for the interleukins *il4* (mediator of Th2 differentiation) and *il6* (key to initiate inflammation) which were expressed at lower levels in constant light. Early inflammatory responses and subsequent initiation of Th2 processes are thought to be critical to resistance of crustacean ectoparasites in salmonids^39^. Taken together, chronic elevation of the immune gene expression – which may result in immune exhaustion^40^ or other immunopathologies^41^ – and reduced ability to mount effective responses key to lice resistance suggest rearing of fish in the absence of light cues are likely to be detrimental for health.

The impact of photoperiod on overall magnitude of immune gene activation is not be the only factor important to parasite resistance; the rhythmicity and the appropriate timing of immune activity (i.e. when fish are maximally vulnerable to pathogen attack) may also be key to pathogen defences. Under regular light-dark cycles, we show trout skin is highly rhythmic in expression of the core vertebrate clock genes and many immune genes in both innate and adaptive pathways. In essence, we find the highest expression of pro-inflammatory markers (e.g. *il6, il17a*) at the onset of the light period and peaks in anti-microbial peptide genes (e.g. cathelecidins) mid-light phase, whilst immunoglobulin and T cell markers were highest during dark hours. The timing of different facets of immune systems, typically peaks of inflammatory mechanisms during active phases and pathways of repair and infection resolution during resting phases, are considered to have evolved to offer hosts greatest protection from invading pathogens when most likely to encounter them, whilst avoiding energetically inefficient and potentially immunopathological risk of continual immune activation^42^. We found that constant light resulted in arrhythmic expression of genes involved in mucosa anti-microbial (e.g. cathelecidins, *igd, il17a*) and Th1 (*tbx21*) responses. Furthermore, genes with phase-shifted expression rhythms in constant light were dominated by those involved in T cell differentiation and regulation (e.g. *cd4, foxp3b, il4, tgfb*). Loss of synchrony between host immunity and parasite activity and/or immune evasion rhythms are very likely to be detrimental for host fitness and survival^43^. Our results indicate that this is a factor in the reduced clearance of lice in fish reared in constant light. Clearly, the impacts of light cycle perturbation, be it intentional such as in aquaculture or unintentionally due to light pollution^44^, must be more carefully considered for animal health.

The primary function of fish skin mucus is as a protective barrier and hosts diverse communities of microbes^45^ which are thought to contribute to protection from microbial pathogens via competitive and/or antagonistic activities^46,47^. While pathogenic taxa occur mostly at low levels in healthy teleost microbiomes, their proliferation is a common signal of microbiome perturbation and dysbiosis^48^. *Argulus* lice infestations are commonly observed alongside bacterial, fungal or viral infections^49^. Here, we demonstrate significant reorganisation of bacterial communities and their potential functional activities in trout skin when infected with *A. foliaceus*, including notable increases in abundance of genera associated with infectious disease^50,51^. Fish lice may elicit host immune profiles and/or destabilize skin microbiota communities resulting in reduced “colonization resistance”^48^, or be direct vectors^52,53^. Further research into the microbiota of *Argulus* and other fish ectoparasites, and their pathogen vectoring capabilities, will be valuable for understanding their role in coinfection dynamics. Intriguingly, trout raised in constant light had a significantly lower microbiome diversity and, when challenged with *Argulus*, exhibited greater shifts in both taxonomic composition and functional potential compared to fish under regular light-dark regimes. Given the growing body of evidence for the importance of “healthy” microbial communities^54^ for effective host homeostasis and disease resistance^55,56^, characterising circadian disruption to microbiomes is important for understanding animal disease risks.

We demonstrate significant daily dynamicity in the skin microbiome of trout; a substantial proportion of bacteria genera exhibit rhythmic changes in relative abundance, suggesting a temporal structure to microbiome functional activity. Parasitic infection appears to perturb microbiome composition, and shift the timings of peak biosynthetic, degradative and energy generation pathway activity in the microbial community. Understanding of the functional importance to the host of commensal microbiota in teleost skin is still in its infancy^48^, and predictive metagenomic analyses are only indicative of actual microbial activity^57^. Temporal metatranscriptomic profiling will be an important means to build upon our results to decipher the functional significance of teleost mucosal microbiota and their daily coordination of activity. Nevertheless, as interest builds towards the utility of microbiome engineering strategies to promote health and productivity in aquaculture^23,48,58^, we propose that a chronobiological understanding of fish microbiomes may be crucial for their effectiveness. The daily rhythms of both fish host immunity and their microbiome communities, for example, could be critical to uptake and establishment of probiotics treatments. Chronotherapeutics – the timed application of treatments and vaccines^59^ – in human medicine holds great promise for improving efficacies but is yet to be given full consideration for managed animal health.

In the mammalian gut – by far the most studied host-microbiome interface – there is a complex interplay between immune factors that shape microbial communities and, conversely, microbiota profoundly affecting immune system development and maintenance^14,15^. Mammal gut microbiome daily rhythms may themselves play a role in host circadian health^60,61^. However, in other tissues, and particularly for non-mammalian vertebrates, host immune-microbiome connectivity and circadian dynamics remains poorly understood. For teleosts, there is evidence that macrophages^62^ and adaptive immune components (e.g. T cells^63^ and immunoglobulins^64^) may be key to mucosal microbiome composition. Our study is the first to present an integrated analysis of skin microbiomes with a broad set of immune and circadian clock gene expression profiles in fish. We found genes of the core secondary feedback loops (e.g. *bmal, clock, rora, csnk1d*) that define the vertebrate molecular clock to be strongly associated with microbial taxa relative abundances in uninfected trout under 12:12 LD, yet these direct clock-microbe associations were largely absent in constant light. Similarly, mice faecal microbiota composition appears closely linked to *bmal1*, with knock-outs resulting in arrhythmicity and altered abundance of microbial taxa^17^. Our results suggest this arm of the biological clock may be pivotal to orchestrating changes in mucosal microbiomes across vertebrates. However, we also find perturbation of microbial communities via ectoparasite infection reconfigures the connectivity of host expression and microbiota. In LL and LD conditions, lice infected fish immune-microbe networks show a greater level of connectivity between host immune gene expression and microbial taxa compared to uninfected individuals. In particular, our results indicate T cell markers to be central to this host-microbiome interface during ectoparasite infection. Under 12:12 LD, we find the T helper cell gene *cd4* to be strongly linked to microbiome composition, whilst in constant light the cytotoxic T cell marker *cd8a* appears to be more influential to microbiome-immune associations. For teleost fish, the ratios and distributions of T cell populations are not well defined^65,66^, although CD4+ and CD8+ subsets appear to have different roles in pathogen defence^67^. Our results suggest their relative importance to shaping fish mucosal microbiomes, or vice versa, warrant further investigation. Disentangling the directionality of the associations we find via controlled manipulations of host immune cell populations, clock gene expression, and microbiota will undoubtedly be key to advancing the concept of circadian holobiont health.

Our study demonstrates the complex daily interaction of fish immune expression and microbiomes, which are impacted by photoperiod and infection status. There is rapidly growing recognition for the detrimental impacts of circadian rhythm perturbation in human medicine^13^, though little attention has been paid to the implications for animal health. In an industry that heavily utilises light manipulation, contemporary aquaculture practices may be significantly exacerbating current disease issues. We provide here an important resource for furthering efforts to integrate chronobiology into animal disease mitigation strategies. In addition, as artificial light at night (i.e. light pollution) encroaches on ever greater proportions of the world’s ecosystems^68^, it is vital studies such as ours are considered for the implications on health and disease dynamics in wild populations.

## Methods

### Experimental design and sample collection

Juvenile female triploid rainbow trout fry (*O. mykiss*, 10 days post-yolk sac absorption, n = 500) were obtained from a commercial hatchery (Bibury Trout Farm, UK). Fry were visually and microscopically determined free of parasitic infections upon arrival and maintained in a re-circulating aquaculture system (RAS) in Cardiff University (water temperature 12 ± 0.5 °C, pH 7.5 ± 0.2). The trout were randomly assigned to duplicate tanks (45 × 60 × 60 cm, 150 L) under one of two photoperiod conditions; 12:12 LD (lights on at zeitgeber time 0; ZT0, off at ZT12) or 24:0 LD (constant light, LL). Each tank was individually illuminated with a full-spectrum white LED bar (80 lux at surface) and surrounded with blackout material to ensure no disturbance from ambient light. Fish were fed with a commercial trout feed (Nutraparr, Skretting, UK) *ad libitum* at ZT2-3 and ZT9-10 daily. Water oxygen saturation (>90%), ammonia (<0.02 mg/L), nitrite (<0.01 mg L–1) and nitrate (<15 mg L–1) were maintained within an appropriate range.

After one month acclimation to light conditions, 130 fish from each light treatment were individually isolated in 1 L plastic containers. Half of the fish from each light treatment (n = 65 per treatment) were individually inoculated with ten *Argulus foliaceus* metanauplii (24 hrs post-hatching). *Argulus* metanauplii were obtained from eggs of wild-caught adult pairs (sourced from Risca Canal, Newport), maintained at Cardiff University. Egg strings were collected and hatched under laboratory conditions according to Stewart et al. (2018). Inoculations were performed at ZT4-5. Fish were individually held in a glass container with 50 ml of tank water and 10 metanauplii added. Fish were observed until all lice had attached (within 2 minutes) and then returned to their 1 L container. Control fish (those not inoculated with *Argulus* lice) were also held for 2 min in 50 ml of water to control for handling stress. Water in all individual containers were changed daily, feeding continued on schedule outlined above, and light conditions maintained at same intensity, spectrum and duration as during acclimation period. The remaining fish were maintained in the RAS system. Once a week, 30 random fish per light treatment were weighed (g) and measured (standard length, SL in cm) for 16 weeks to monitor growth rates. General linear models of standard length and weight, including photoperiod and sampling day, were used to assess differences in growth between light treatments. All procedures were performed under Home Office project license PPL 303424 with full approval of Cardiff University Animal Ethics committee.

One week after inoculation, sampling of fish was performed over a 48 h period to encompass two full circadian cycles. Starting at ZT0 (lights on in 12:12 LD treatment), every 4 h, five fish from each condition (12:12 LD control, 12:12 LD *Argulus*-infected, LL control, LL *Argulus*–infected) were euthanised using an overdose of tricaine methanesulfonate (MS222, 500 mg L-1) according to Home Office Schedule 1. At timepoints during dark periods in 12:12 LD treatment, fish were handled and euthanised in dim red light. Immediately after euthanasia, infected fish were visually inspected to quantify number of lice surviving and the lice removed to ensure they were not included in tissue samples. Welch’s two sample T test was used to determine difference in infection load (number of *Argulus*) between light treatments. All sampled fish were weighed (g) and measured (standard length, SL in cm). Skin swabs (MWE MW-100) were rubbed along the entire lateral body surface five times each side and immediately frozen at −80 C to preserve skin mucus microbiota for DNA extraction. All skin from immediately posterior to opercula to the caudal peduncle was dissected using sterile forceps, preserved in RNAlater (Invitrogen), and stored at −80 C until RNA extraction. All dissections for each timepoint were performed within an 1 hour window. At each timepoint-treatment combination, 10 ml of water from all containers was pooled and frozen at −80°C to provide background controls for skin microbiome analyses. To test for endogenous expression rhythms, an additional 65 uninfected fish maintained at 12:12 LD were individually isolated and held in constant darkness (DD). After 24 h, starting at ZT0, five fish every 4 h were sampled as above.

### RNA extraction, gene expression quantification and analyses

Total RNA was individually extracted from each skin sample using RNeasy Mini kits (Qiagen). RNA was quantified using Qubit Broad Range RNA assays (ThermoFisher Scientific). mRNA expression patterns in the skin were measured by Nanostring analysis, following manufacturer’s guidelines, at Liverpool Centre for Genomic Research. The nCounter PlexSet oligonucleotide and probe design was performed at NanoString Technologies (NanoString Technologies) for 48 genes, including four housekeeping genes (Supplementary Table S1). The oligonucleotide probes were synthesized at Integrated DNA Technologies. Titration reactions were performed according to supplier’s instructions with RNA inputs between 250 ng and 700 ng to determine the required RNA amount for hybridization reaction. 600 ng total RNA per sample was used for PlexSet hybridization reaction for 20 h according to manufacturer’s instructions.

Samples were processed on a nCounter MAX prep station (NanoString Technologies) and cartridges were scanned in a generation II nCounter Digital Analyzer (NanoString Technologies). RCC files (nCounter data files) were used for data analysis. RCC files were imported into the NanoString nSolver 4.0 analysis software and raw data pre-processing and normalization was performed according to manufacturer’s instructions for standard procedures (positive normalization to geomean of top 3 positive controls, codeset content normalization using housekeeping genes *hprt1, polr1b, polr2i* and codeset calibration with the reference sample). The housekeeping gene *rplp0* and *aanat2* expression were not detected and excluded from analyses.

To assess overall differences in immune responses to infection under the different light treatments, pairwise t-tests comparing normalised expression of immune genes were performed in R (version 4.0.3). To detect rhythmicity in expression of clock and immune genes, empirical JTK Cycle (eJTK_cycle^69^) analyses were applied with a set period of 24 h, a phase search every 4 h from ZT0 to ZT20, and an asymmetry search every 4 h from ZT4 to ZT20. FDR-corrected empirical p-values less than 0.1 were considered moderately rhythmic^70–72^, and less than 0.05 strongly rhythmic. CircaCompare^31^ was used to estimate rhythmic genes’ peak expression time, mesor and amplitude, and to statistically contrast rhythms.

### DNA extraction, 16S rRNA gene amplification, Illumina sequencing and analyses

DNA was extracted from skin swabs using Qiagen DNeasy Blood and Tissue kits according to ^73^ to maximise lysis of microbiome community and DNA recovery. PCR amplification of the 16S rRNA V4 region, using 515F and 806R primers, was performed in triplicate for each DNA extract, pooled and prepared for Illumina MiSeq sequencing according to ^74^. Gel electrophoresis was used to estimate concentrations for pooling individual amplicon libraries. Negative controls for extractions and PCR, and mock community positive controls were included for sequencing. Libraries were sequenced using a 2 x 250 bp Illumina MiSeq run at the Cardiff Biosciences Genomics Hub.

Paired-end demultiplexed Illumina sequencing reads were imported into the Quantitative Insights Into Microbial Ecology 2 (QIIME2^75^). Sequences were then quality filtered, dereplicated, chimeras identified and paired-end reads merged in QIIME2 using DADA2 with default settings. Classification of Amplicon Sequence Variants (ASVs) was performed using a Naïve Bayes algorithm trained using sequences representing the bacterial V4 rRNA region available from the SILVA database (https://www.arb-silva.de/download/archive/qiime;Silva_132), and the corresponding taxonomic classifications were obtained using the q2-feature-classifier plugin in QIIME2. The classifier was then used to assign taxonomic information to representative sequences of each ASV. Following rarefaction analysis, samples with less than 2000 sequences were excluded from further analyses. QIIME2 was used to analyse alpha (Kruskal-Wallis pairwise tests of Faith’s phylogenetic distance) and beta (pairwise PERMANOVA) diversity measures. ASVs were filtered to exclude those assigned to eukaryotes or eukaryotic organelles and include ones with at least 100 copies in at least two samples. The QIIME2 output data were imported in RStudio (Version 1.3.959) with the Bioconductor package phyloseq^76^, for subsetting, normalizing, and plotting of the data.

Differential abundance of ASVs between healthy and infected fish in both light treatments were determined using DESeq2^77^, with FDR-corrected p-values less than 0.05 considered significant. Differential abundances of all taxonomic levels were also determined and visualised using MicrobiomeAnalyst^78^ heat trees using default settings. We inferred the microbial gene content from the taxa abundance using PICRUSt2^79^. We used LefSe analyses to identify group differences in the inferred gene abundance of MetaCyc pathways, using the online galaxy server (https://huttenhower.sph.harvard.edu/galaxy/). LDA scores >2.0 were considered significant. Rhythmicity of microbial genera and MetaCyc pathway abundances were determined following the same methods as gene expression (see above). To determine potential associations of host gene expression and the microbiome, Spearman correlation tests were performed including only genera found in at least 50% of samples in each treatment group. Corrected p-values (using qvalue R package) of less than 0.05 were considered significantly correlated. Correlation networks were visualised using gephi^80^ and influential nodes determined using degree centrality scores and number of connections (degrees).

## Supporting information

Supplementary Datafile 1

## Acknowledgments

This study was funded by a BBSRC Discovery Fellowship awarded to AE (BB/R010609/1). We thank Liverpool Centre for Genomic Research and Cardiff Biosciences Genome Hub for their assistance in data generation. We also thank members of the Cable research group for their assistance in animal husbandry.

---

**Supplementary Figure 1:**
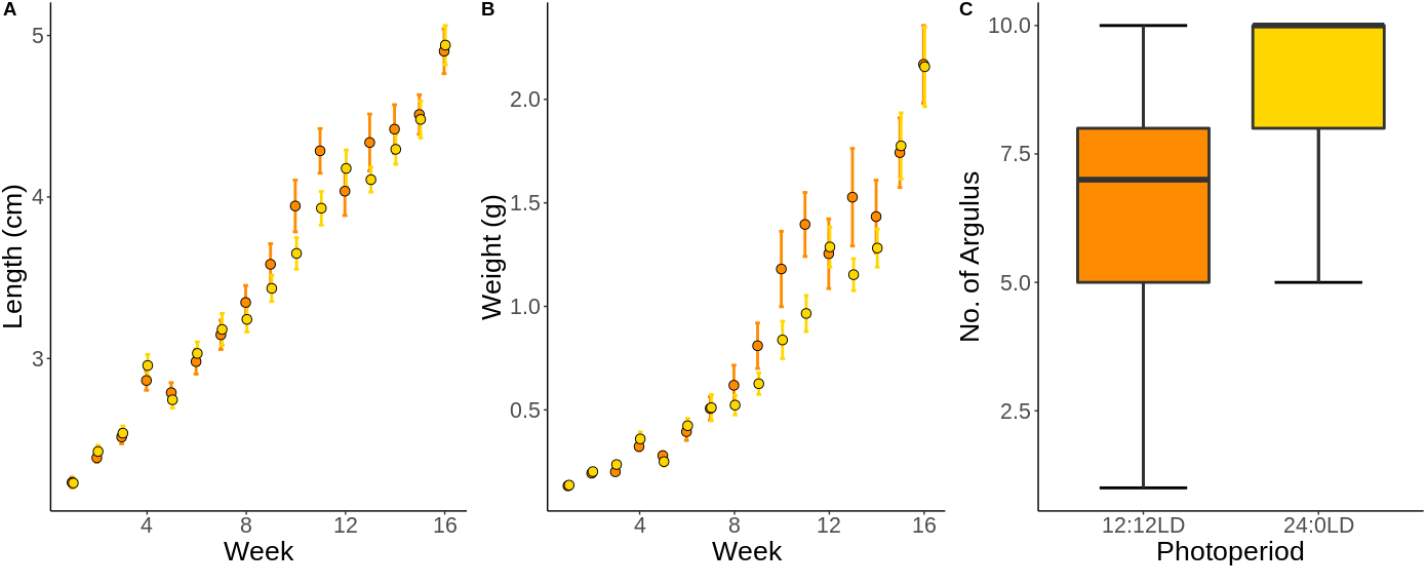
Average A) standard length and B) weight of trout (±1 S.E.) over 16-week growth trial under 12:12 LD (orange) and 24:0 LD (yellow). C) Boxplots of number of *Argulus foliaceus* lice infecting fish 7 days post-inoculation.

**Supplementary Figure 2:**
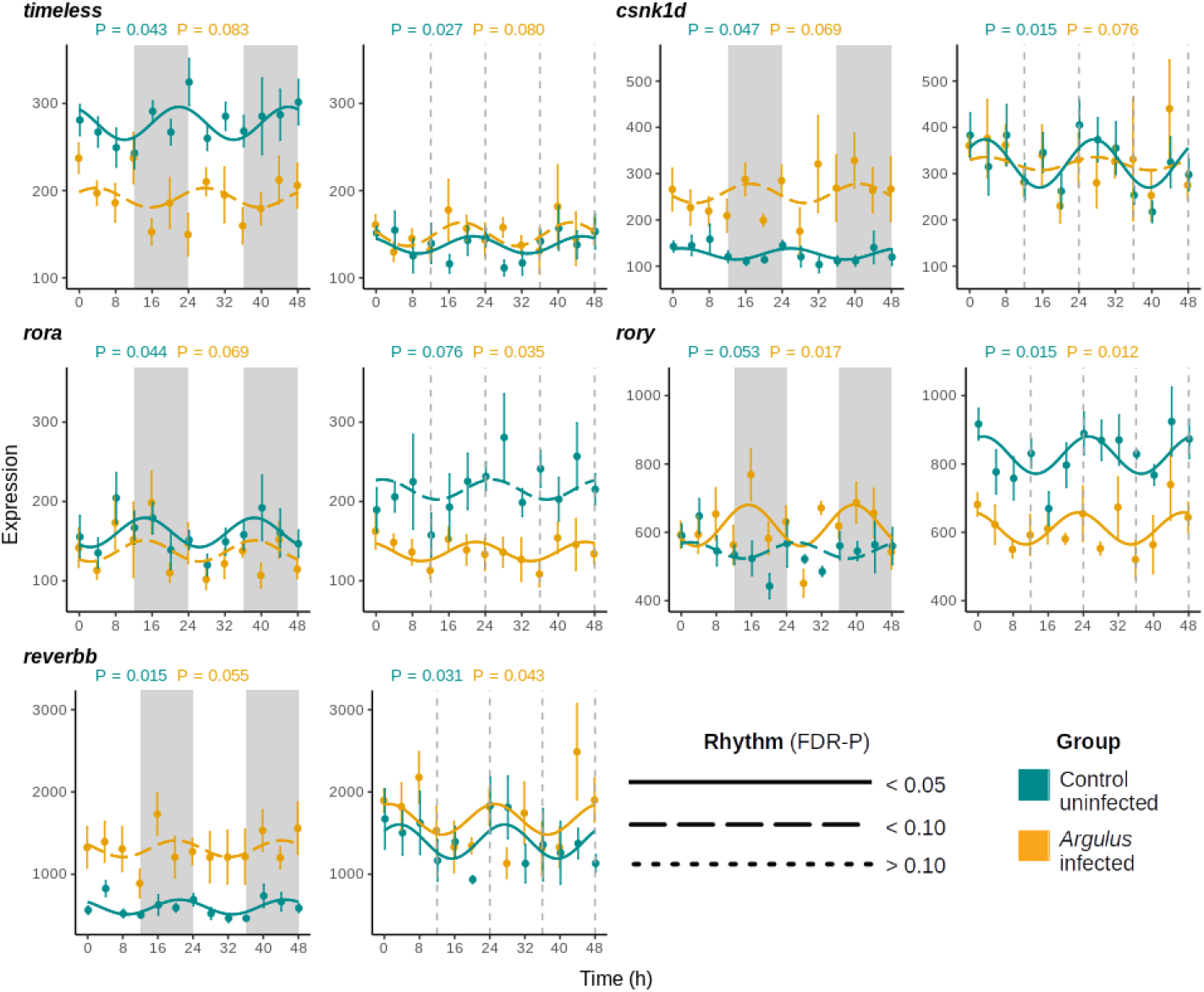
Mean expression (± 1 S.E.) of accessory clock genes of uninfected (cyan) and *Argulus*-infected (orange) rainbow trout maintained at 12:12 LD (left) and 24:0 LD (LL, right). Expression is normalised counts of mRNA copies detected via Nanostring nCounter. Curves denote cosinor waveform fitted using CircaCompare. Grey shading indicates time periods in darkness (grey dashing indicates equivalent 12:12 LD light transitions on LL plots).

**Supplementary Figure 3:**
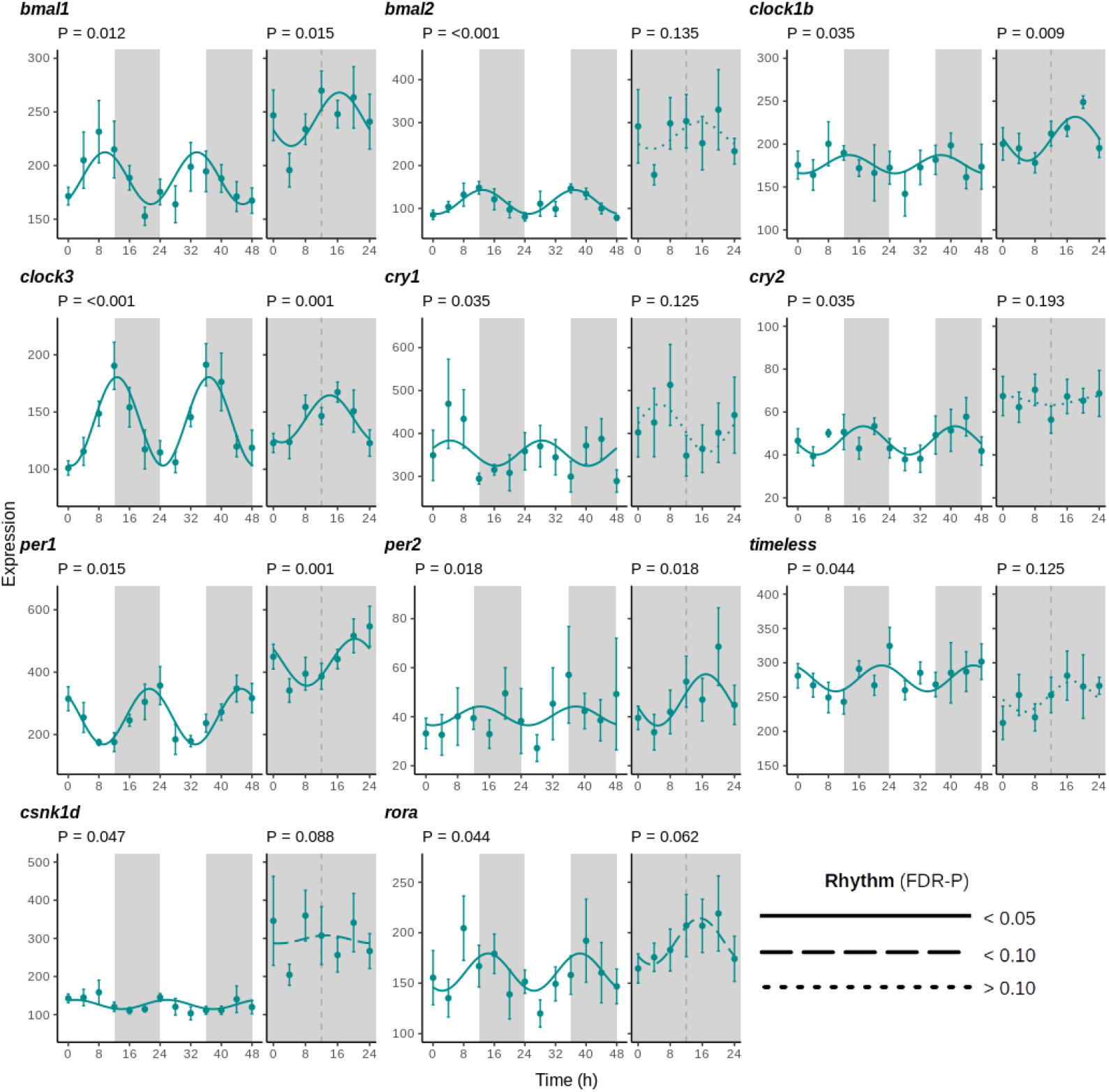
Mean expression (± 1 S.E.) of clock genes of rainbow trout under 12:12 LD and DD. Expression is normalised counts of mRNA copies detected via Nanostring nCounter. Curves denote cosinor waveform fitted using CircaCompare. Grey shading indicates time periods in darkness (grey dashing indicates subjective day-night transition in DD).

**Supplementary Figure 4:**
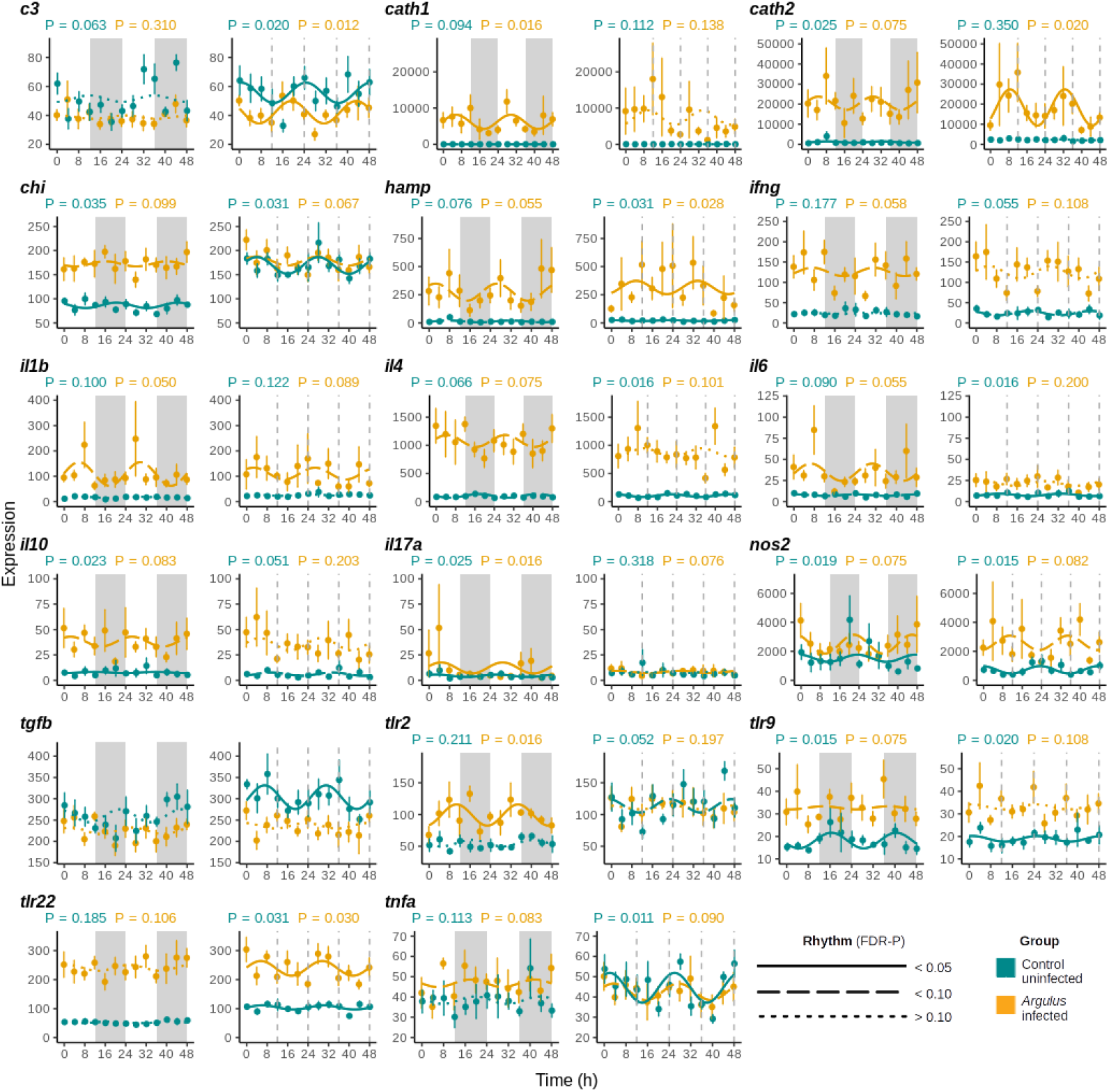
Mean expression (± 1 S.E.) of innate immune genes of uninfected (cyan) and *Argulus*-infected (orange) rainbow trout maintained at 12:12 LD (left) and 24:0 LD (LL, right). Expression is normalised counts of mRNA copies detected via Nanostring nCounter. Curves denote cosinor waveform fitted using CircaCompare. Grey shading indicates time periods in darkness (grey dashing indicates equivalent 12:12 LD light transitions on LL plots). Only genes with significant rhythm in one or more groups shown.

**Supplementary Figure 5:**
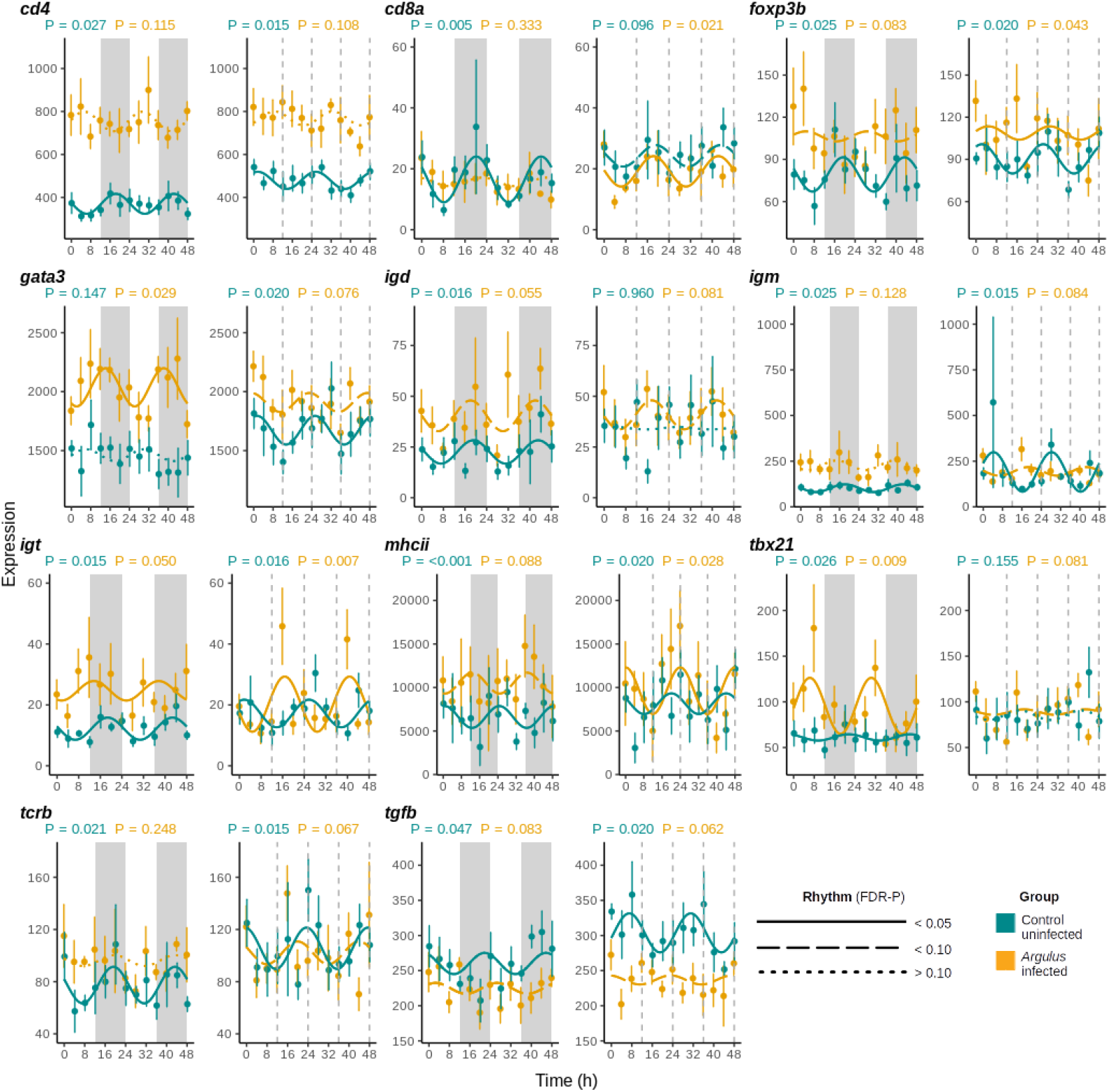
Mean expression (± 1 S.E.) of adaptive immune genes of uninfected (cyan) and Argulus-infected (orange) rainbow trout maintained at 12:12 LD (left) and 24:0 LD (LL, right). Expression is normalised counts of mRNA copies detected via Nanostring nCounter. Curves denote cosinor waveform fitted using CircaCompare. Grey shading indicates time periods in darkness (grey dashing indicates equivalent 12:12 LD light transitions on LL plots). Only genes with significant rhythm in one or more groups shown.

**Supplementary Figure 6:**
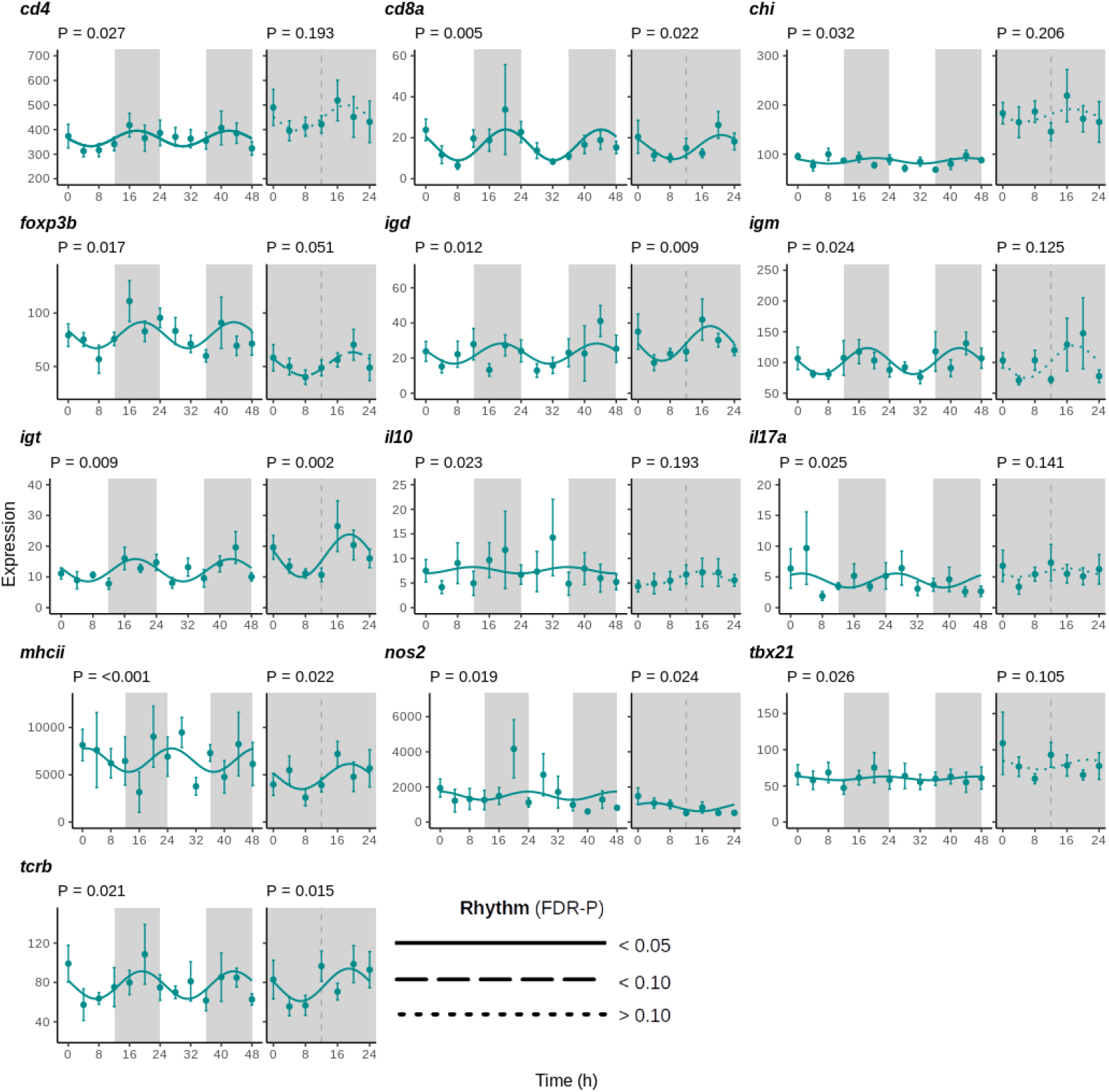
Mean expression (± 1 S.E.) of immune genes of rainbow trout under 12:12 LD and DD. Expression is normalised counts of mRNA copies detected via Nanostring nCounter. Curves denote cosinor waveform fitted using CircaCompare. Grey shading indicates time periods in darkness (grey dashing indicates subjective day-night transition in DD).

**Supplementary Figure 7:**
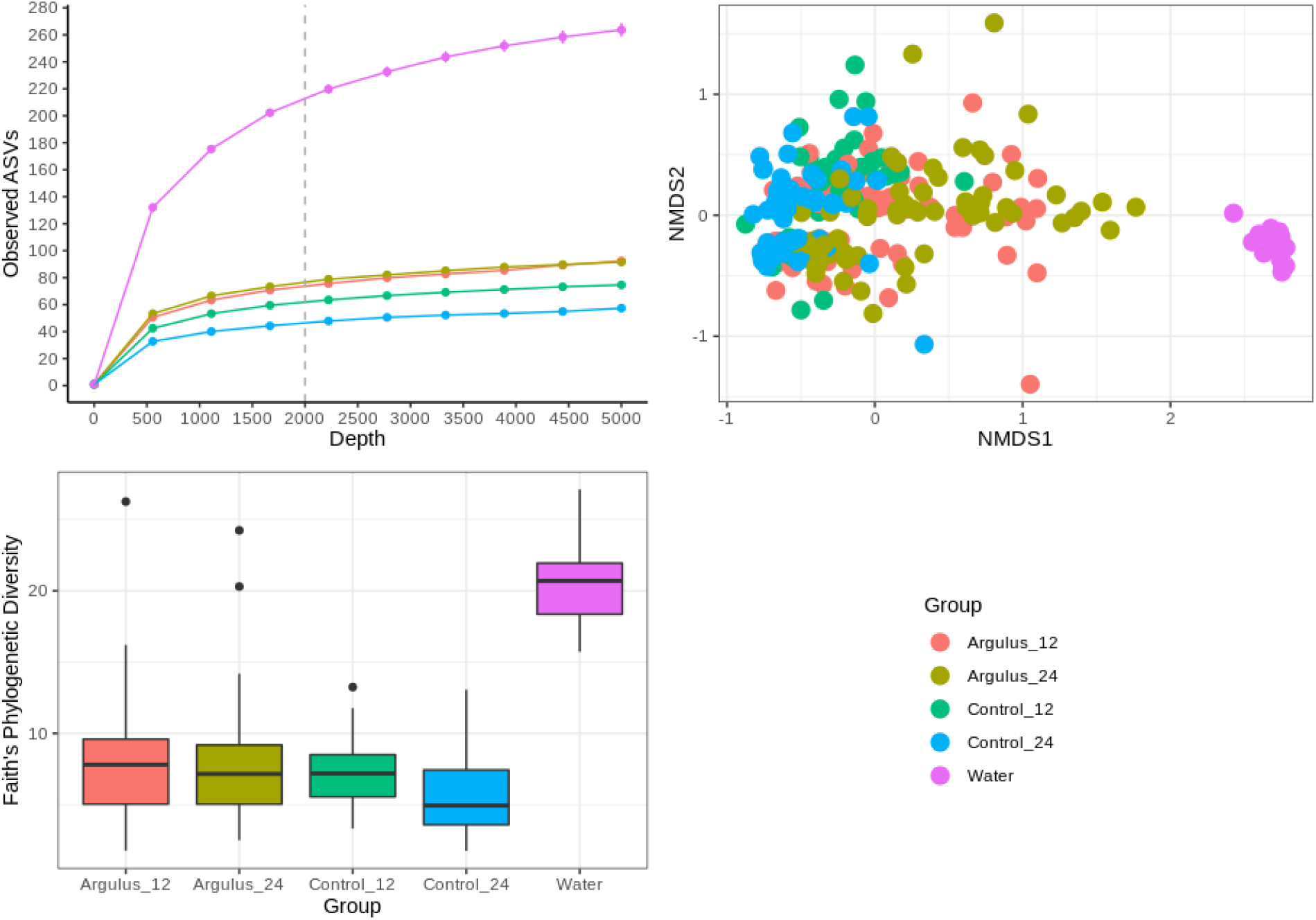
A) Rarefaction plots of detected amplified sequence variants (ASVs) by sampling depth. B) NMDS ordination of microbiome profiles. C) Alpha diversity plots by treatment group.

**Supplementary Figure 8:**
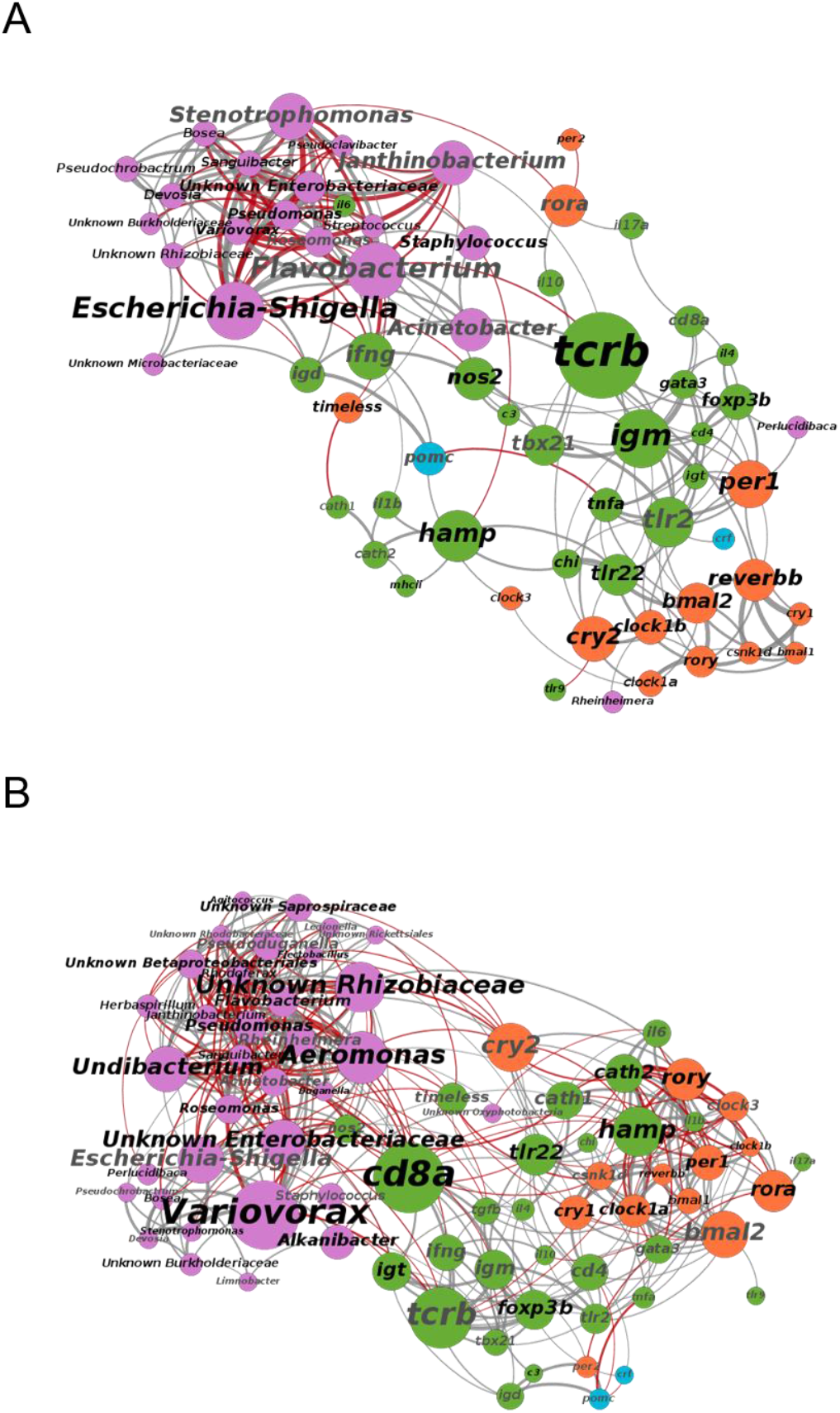
Co-occurrence networks of microbial genera (pink) and host gene expression (orange = clock, green = immune, blue = corticotropin) in healthy (top) and *Argulus*-infected (bottom) trout under 24:0 LD. Node and label size scaled to degree centrality score. Label colour denotes rhythmicity (black = rhythm FDR p-value <0.05, grey = rhythm FDR p-value >0.05). Connection colour indicates association (grey = positive, red = negative, determined by Spearman correlation tests) and connection width scaled to correlation strength (thicker lines denote a higher correlation coefficient).

## References

1. Dunlap, J. C. Molecular bases for circadian clocks. Cell 96, 271–290 (1999).

2. Lochmiller, R. L. & Deerenberg, C. Trade-offs in evolutionary immunology: Just what is the cost of immunity? Oikos 88, 87–98 (2000).

3. Wang, W. et al. Timing of plant immune responses by a central circadian regulator. Nature 470, 110–114 (2011).

4. Scheiermann, C., Kunisaki, Y. & Frenette, P. S. Circadian control of the immune system. Nat. Rev. Immunol. 13, 190–198 (2013).

5. Curtis, A. M., Bellet, M. M., Sassone-Corsi, P. & O’Neill, L. A. J. Circadian clock proteins and immunity. Immunity 40, 178–186 (2014).

6. Okada, K. et al. Injection of LPS causes transient suppression of biological clock genes in rats. J. Surg. Res. 145, 5–12 (2008).

7. Cavadini, G. et al. TNF-α suppresses the expression of clock genes by interfering with E-box-mediated transcription. Proc. Natl. Acad. Sci. 104, 12843–12848 (2007).

8. de Leone, M. J. et al. Bacterial Infection Disrupts Clock Gene Expression to Attenuate Immune Responses. Curr. Biol. (2020).

9. Shirasu-Hiza, M. M., Dionne, M. S., Pham, L. N., Ayres, J. S. & Schneider, D. S. Interactions between circadian rhythm and immunity in *Drosophila melanogaster*. Curr. Biol. 17, R353–R355 (2007).

10. Marpegán, L., Bekinschtein, T. A., Costas, M. A. & Golombek, D. A. Circadian responses to endotoxin treatment in mice. J. Neuroimmunol. 160, 102–109 (2005).

11. Castanon-Cervantes, O. et al. Dysregulation of inflammatory responses by chronic circadian disruption. J. Immunol. 185, 5796–5805 (2010).

12. Adams, K. L., Castanon-Cervantes, O., Evans, J. A. & Davidson, A. J. Environmental circadian disruption elevates the IL-6 response to lipopolysaccharide in blood. J. Biol. Rhythms 28, 272–277 (2013).

13. Touitou, Y., Reinberg, A. & Touitou, D. Association between light at night, melatonin secretion, sleep deprivation, and the internal clock: Health impacts and mechanisms of circadian disruption. Life Sci. 173, 94–106 (2017).

14. Thaiss, C. A., Zmora, N., Levy, M. & Elinav, E. The microbiome and innate immunity. Nature 535, 65 (2016).

15. McDermott, A. J. & Huffnagle, G. B. The microbiome and regulation of mucosal immunity. Immunology 142, 24–31 (2014).

16. Zarrinpar, A., Chaix, A., Yooseph, S. & Panda, S. Diet and feeding pattern affect the diurnal dynamics of the gut microbiome. Cell Metab. 20, 1006–1017 (2014).

17. Liang, X., Bushman, F. D. & FitzGerald, G. A. Rhythmicity of the intestinal microbiota is regulated by gender and the host circadian clock. Proc. Natl. Acad. Sci. 112, 10479–10484 (2015).

18. Leone, V. et al. Effects of diurnal variation of gut microbes and high-fat feeding on host circadian clock function and metabolism. Cell Host Microbe 17, 681–689 (2015).

19. Stentiford, G. D. et al. New paradigms to help solve the global aquaculture disease crisis. PLoS Pathog. 13, e1006160 (2017).

20. Lafferty, K. D. et al. Infectious diseases affect marine fisheries and aquaculture economics. (2015).

21. Tarnecki, A. M., Burgos, F. A., Ray, C. L. & Arias, C. R. Fish intestinal microbiome: diversity and symbiosis unravelled by metagenomics. J. Appl. Microbiol. 123, 2–17 (2017).

22. Ghanbari, M., Kneifel, W. & Domig, K. J. A new view of the fish gut microbiome: advances from next-generation sequencing. Aquaculture 448, 464–475 (2015).

23. Perry, W. B., Lindsay, E., Payne, C. J., Brodie, C. & Kazlauskaite, R. The role of the gut microbiome in sustainable teleost aquaculture. Proc. R. Soc. B 287, 20200184 (2020).

24. Biswas, A. K., Seoka, M., Tanaka, Y., Takii, K. & Kumai, H. Effect of photoperiod manipulation on the growth performance and stress response of juvenile red sea bream (*Pagrus major*). Aquaculture 258, 350–356 (2006).

25. Rad, F., Bozaoğlu, S., Gözükara, S. E., Karahan, A. & Kurt, G. Effects of different long-day photoperiods on somatic growth and gonadal development in Nile tilapia (*Oreochromis niloticus* L.). Aquaculture 255, 292–300 (2006).

26. Berrill, I. K., Porter, M. J. R., Smart, A., Mitchell, D. & Bromage, N. R. Photoperiodic effects on precocious maturation, growth and smoltification in Atlantic salmon, *Salmo salar*. Aquaculture 222, 239–252 (2003).

27. Onoue, T., Nishi, G., Hikima, J., Sakai, M. & Kono, T. Circadian oscillation of TNF-α gene expression regulated by clock gene, BMAL1 and CLOCK1, in the Japanese medaka (*Oryzias latipes*). Int. Immunopharmacol. 70, 362–371 (2019).

28. Zhang, P., Yu, C. & Sun, L. Japanese flounder (*Paralichthys olivaceus*) Bmal1 is involved in the regulation of inflammatory response and bacterial infection. Aquaculture 735330 (2020).

29. Frøland Steindal, I. A. & Whitmore, D. Circadian clocks in fish—what have we learned so far? Biology (Basel). 8, 17 (2019).

30. Binuramesh, C. & Michael, R. D. Diel variations in the selected serum immune parameters in *Oreochromis mossambicus*. Fish Shellfish Immunol. 30, 824–829 (2011).

31. Ángeles Esteban, M., Cuesta, A., Rodríguez, A. & Meseguer, J. Effect of photoperiod on the fish innate immune system: a link between fish pineal gland and the immune system. J. Pineal Res. 41, 261–266 (2006).

32. Lazado, C. C., Skov, P. V. & Pedersen, P. B. Innate immune defenses exhibit circadian rhythmicity and differential temporal sensitivity to a bacterial endotoxin in Nile tilapia (*Oreochromis niloticus*). Fish Shellfish Immunol. 55, 613–622 (2016).

33. Taira, G., Onoue, T., Hikima, J., Sakai, M. & Kono, T. Circadian clock components Bmal1 and Clock1 regulate tlr9 gene expression in the Japanese medaka (*Oryzias latipes*). Fish Shellfish Immunol. 105, 438–445 (2020).

34. Ellison, A. R. et al. Transcriptomic response to parasite infection in Nile tilapia (*Oreochromis niloticus*) depends on rearing density. BMC Genomics 19, 723 (2018).

35. Ellison, A. R. et al. Comparative transcriptomics reveal conserved impacts of rearing density on immune response of two important aquaculture species. Fish Shellfish Immunol. (2020).

36. Greer, R., Dong, X., Morgun, A. & Shulzhenko, N. Investigating a holobiont: Microbiota perturbations and transkingdom networks. Gut Microbes 7, 126–135 (2016).

37. Greenblum, S., Turnbaugh, P. J. & Borenstein, E. Metagenomic systems biology of the human gut microbiome reveals topological shifts associated with obesity and inflammatory bowel disease. Proc. Natl. Acad. Sci. 109, 594–599 (2012).

38. Whiting, J. R., Mahmud, M. A., Bradley, J. E. & MacColl, A. D. C. Prior exposure to long-day photoperiods alters immune responses and increases susceptibility to parasitic infection in stickleback. Proc. R. Soc. B 287, 20201017 (2020).

39. Braden, L. M., Koop, B. F. & Jones, S. R. M. Signatures of resistance to *Lepeophtheirus salmonis* include a TH2-type response at the louse-salmon interface. Dev. Comp. Immunol. 48, 178–191 (2015).

40. Saeidi, A. et al. T-cell exhaustion in chronic infections: reversing the state of exhaustion and reinvigorating optimal protective immune responses. Front. Immunol. 9, 2569 (2018).

41. Graham, A. L., Allen, J. E. & Read, A. F. Evolutionary causes and consequences of immunopathology. Annu. Rev. Ecol. Evol. Syst. 373–397 (2005).

42. Westwood, M. L. et al. The evolutionary ecology of circadian rhythms in infection. Nat. Ecol. Evol. 3, 552–560 (2019).

43. Reece, S. E., Prior, K. F. & Mideo, N. The life and times of parasites: rhythms in strategies for within-host survival and between-host transmission. J. Biol. Rhythms 32, 516–533 (2017).

44. Longcore, T. & Rich, C. Ecological light pollution. Front. Ecol. Environ. 2, 191–198 (2004).

45. Ross, A. A., Hoffmann, A. R. & Neufeld, J. D. The skin microbiome of vertebrates. Microbiome 7, 1–14 (2019).

46. Pérez-Sánchez, T. et al. Identification and characterization of lactic acid bacteria isolated from rainbow trout, *Oncorhynchus mykiss* (Walbaum), with inhibitory activity against *Lactococcus garvieae*. J. Fish Dis. 34, 499–507 (2011).

47. Balcázar, J. L. et al. In vitro competitive adhesion and production of antagonistic compounds by lactic acid bacteria against fish pathogens. Vet. Microbiol. 122, 373–380 (2007).

48. Llewellyn, M. S., Boutin, S., Hoseinifar, S. H. & Derome, N. Teleost microbiomes: the state of the art in their characterization, manipulation and importance in aquaculture and fisheries. Roles Mech. Parasit. Aquat. Microb. Communities 109 (2015).

49. Walker, P. D., Flik, G. & Bonga, S. E. W. The biology of parasites from the genus *Argulus* and a review of the interactions with its host. Host-Parasite Interact. 55, 107–129 (2004).

50. Adikesavalu, H., Patra, A., Banerjee, S., Sarkar, A. & Abraham, T. J. Phenotypic and molecular characterization and pathology of *Flectobacillus roseus* causing flectobacillosis in captive held carp Labeo rohita (Ham.) fingerlings. Aquaculture 439, 60–65 (2015).

51. Loch, T. P. & Faisal, M. Emerging flavobacterial infections in fish: a review. J. Adv. Res. 6, 283–300 (2015).

52. Jakob, E., Barker, D. E. & Garver, K. A. Vector potential of the salmon louse *Lepeophtheirus salmonis* in the transmission of infectious haematopoietic necrosis virus (IHNV). Dis. Aquat. Organ. 97, 155–165 (2011).

53. Ahne, W. *Argulus foliaceus* L. and *Piscicola geometra* L. as mechanical vectors of spring viraemia of carp virus (SVCV). J. Fish Dis. 8, 241–242 (1985).

54. Lloyd-Price, J., Abu-Ali, G. & Huttenhower, C. The healthy human microbiome. Genome Med. 8, 1–11 (2016).

55. Harris, E. V, de Roode, J. C. & Gerardo, N. M. Diet–microbiome–disease: Investigating diet’s influence on infectious disease resistance through alteration of the gut microbiome. PLoS Pathog. 15, e1007891 (2019).

56. Cani, P. D. et al. Microbial regulation of organismal energy homeostasis. Nat. Metab. 1, 34–46 (2019).

57. Fiore, C. L., Jarett, J. K., Steinert, G. & Lesser, M. P. Trait-Based Comparison of Coral and Sponge Microbiomes. Sci. Rep. 10, 1–16 (2020).

58. Pérez-Sánchez, T., Ruiz-Zarzuela, I., de Blas, I. & Balcázar, J. L. Probiotics in aquaculture: a current assessment. Rev. Aquac. 6, 133–146 (2014).

59. Ballesta, A., Innominato, P. F., Dallmann, R., Rand, D. A. & Lévi, F. A. Systems chronotherapeutics. Pharmacol. Rev. 69, 161–199 (2017).

60. Pearson, J. A., Wong, F. S. & Wen, L. Cross talk between circadian rhythms and the microbiota. Immunology (2020).

61. Gibbs, J. E. & Butler, T. D. Circadian host-microbiome interactions in immunity. Front. Immunol. 11, 1783 (2020).

62. Earley, A. M., Graves, C. L. & Shiau, C. E. Critical role for a subset of intestinal macrophages in shaping gut microbiota in adult zebrafish. Cell Rep. 25, 424–436 (2018).

63. Brugman, S. et al. T lymphocytes control microbial composition by regulating the abundance of *Vibrio* in the zebrafish gut. Gut Microbes 5, 737–747 (2014).

64. Xu, Z. et al. Specialization of mucosal immunoglobulins in pathogen control and microbiota homeostasis occurred early in vertebrate evolution. Sci. Immunol. 5, (2020).

65. Takizawa, F. et al. The expression of CD8α discriminates distinct T cell subsets in teleost fish. Dev. Comp. Immunol. 35, 752–763 (2011).

66. Kelly, C. & Salinas, I. Under pressure: interactions between commensal microbiota and the teleost immune system. Front. Immunol. 8, 559 (2017).

67. Sukeda, M. et al. Innate cell-mediated cytotoxicity of CD8+ T cells against the protozoan parasite Ichthyophthirius multifiliis in the ginbuna crucian carp, *Carassius auratus langsdorfii*. Dev. Comp. Immunol. 115, 103886 (2020).

68. Hölker, F. et al. The dark side of light: a transdisciplinary research agenda for light pollution policy. Ecol. Soc. 15, (2010).

69. Hutchison, A. L. et al. Improved statistical methods enable greater sensitivity in rhythm detection for genome-wide data. PLoS Comput Biol 11, e1004094 (2015).

70. Wang, Y. et al. A proteomics landscape of circadian clock in mouse liver. Nat. Commun. 9, 1–16 (2018).

71. Lafaye, G., Desterke, C., Marulaz, L. & Benyamina, A. Cannabidiol affects circadian clock core complex and its regulation in microglia cells. Addict. Biol. 24, 921–934 (2019).

72. Cui, P. et al. Identification of human circadian genes based on time course gene expression profiles by using a deep learning method. Biochim. Biophys. Acta (BBA)-Molecular Basis Dis. 1864, 2274–2283 (2018).

73. Gill, C., van de Wijgert, J. H. H. M., Blow, F. & Darby, A. C. Evaluation of lysis methods for the extraction of bacterial DNA for analysis of the vaginal microbiota. PLoS One 11, e0163148 (2016).

74. Caporaso, J. G. et al. Ultra-high-throughput microbial community analysis on the Illumina HiSeq and MiSeq platforms. ISME J. 6, 1621–1624 (2012).

75. Bolyen, E. et al. Reproducible, interactive, scalable and extensible microbiome data science using QIIME 2. Nat. Biotechnol. 37, 852–857 (2019).

76. McMurdie, P. J. & Holmes, S. phyloseq: an R package for reproducible interactive analysis and graphics of microbiome census data. PLoS One 8, e61217 (2013).

77. Love, M. I., Huber, W. & Anders, S. Moderated estimation of fold change and dispersion for RNA-Seq data with DESeq2. Genome Biol. 15, 550 (2014).

78. Dhariwal, A. et al. MicrobiomeAnalyst: a web-based tool for comprehensive statistical, visual and meta-analysis of microbiome data. Nucleic Acids Res. 45, W180–W188 (2017).

79. Douglas, G. M. et al. PICRUSt2 for prediction of metagenome functions. Nat. Biotechnol. 1–5 (2020).

80. Bastian, M., Heymann, S. & Jacomy, M. Gephi: an open source software for exploring and manipulating networks. in Proceedings of the International AAAI Conference on Web and Social Media vol. 3 (2009).

