## Supplementary Table 4 for "Circadian dynamics of the teleost skin immune-microbiome interface"

| ASV_ID | baseMean | log2FoldChange | lfcSE |
|---|---|---|---|
| 98f49c75988e4d0fd97a180db1678fc8 | 48.8330818 | 27.01540348 | 1.013805693 |
| 9ac9a274b64edbcb2b5e1e1c81743b5 | 10.35692861 | 26.20394757 | 1.138996505 |
| 113e5cb2dbb0b2086e230fc9d845310f | 54.74450093 | 25.48020055 | 1.274527312 |
| b9fcd7d71b74853248517b892d03a745 | 55.31630097 | 23.11054915 | 1.418520085 |
| 88524108706087f7c75e79d6c10c9c61 | 10.96072901 | 25.92929447 | 1.596932892 |
| eb2625c1474f98f554b002fcd5d1088e | 62.89240942 | 8.674284705 | 0.739574633 |
| f9c1c7de158ab4d555ec46bec26478f1 | 47.18641453 | 7.644580693 | 0.840986671 |
| 931bf16c32086439f09655b848d87174 | 369.3952111 | 4.031701206 | 0.504054039 |
| 04175f386c14319829ffb23d5c188b2a | 260.8060318 | 4.697233759 | 0.629259993 |
| 8b1dba1a82d5149143975ddc2cc1089f | 91.55719942 | 6.303940766 | 0.878099271 |
| f34cdcd73e535507d29c29b19d01f658 | 71.11788206 | 6.54142063 | 0.941942278 |
| bbf772e87bb1bd04ae18adfa03247a9f | 89.44846459 | 6.74907595 | 0.984665176 |
| f89e77104762c7e39a9e0fb88c3ba6b3 | 20.45091495 | 8.112140104 | 1.213930156 |
| 783869922c5a707e99ebebcabff8c5c6 | 84.96323871 | 4.144967739 | 0.759529828 |
| d0fbc9aff818121519652ff493a83314 | 71.1331293 | 6.027680264 | 1.125915874 |
| adf8882d5fd60155f03f9a52e900d32d | 118.8179065 | 2.910662125 | 0.566679495 |
| 3dbf66ee7707a9c91038139380697b07 | 46.42682375 | 4.819237145 | 0.942892662 |
| 4cfe5b5468150f8a54cf003ea4c15603 | 112.7343322 | 4.057568617 | 0.798828244 |
| 709d785d9cfcbd7a92cfcc5fae6828f0 | 11.01312335 | 6.537485046 | 1.374795193 |
| 38c27ceaed634984c1225a82648cf571 | 280.8876036 | 4.926287362 | 1.035107818 |
| c0f1efd28e2ac632b9821285c77fe072 | 16.91911017 | 4.336944324 | 0.921582164 |
| 2cd00ea627d63b6c55af5d1d65a6cd44 | 11.31504735 | 5.63910242 | 1.225751035 |
| c0c6dea060404ea17972053bc9e98ae | 15.15068662 | 4.115126058 | 0.902330107 |
| 3683c1682210f16c9067d22b4f404929 | 5.392930686 | 6.688284999 | 1.542126568 |
| 4e6abee1bbb9afea6d2c733a33f76d8e | 8.00197725 | 5.256195586 | 1.238972344 |
| 6e6f306a337ba255a7f4a7f8469ff691 | 12.79955601 | 7.079133322 | 1.69717923 |
| 3badab07e5965e92e5954b746c192d5 | 18.06512302 | 6.268354428 | 1.519269553 |
| e416d0916760d2fc17b616e2ac3ad855 | 5.0021677 | 5.701240413 | 1.400204261 |
| 69179c4a6011d0c9d30036cf18626520 | 66.39688045 | 2.751739881 | 0.687568473 |
| 107273e73071282e4d0f4a0b46548da5 | 90.1483836 | 3.048492046 | 0.806838096 |
| 095dcca0a1cacb655451100eb59c8ef1 | 12.63366984 | 3.519908903 | 1.041088014 |
| 892a0fc53b26d6a2c193d145b2464da7 | 3.215386764 | 5.422233907 | 1.618297778 |
| a8d4ddd7dbd400a101b2d3f1b3babaf2 | 17.37632741 | 4.013534759 | 1.259014328 |
| 52abfae86a69569ae491695349cca6f8 | 6.926038692 | 6.872937326 | 2.19013485 |
| c757ad670c5d796cb866c941c62c4e33 | 38.90084495 | 3.425291977 | 1.15754519 |
| e371861e8798027b6e72500f333c78cc | 3.441387683 | -5.287932895 | 1.852435713 |
| fca1ed7792b0c80cd00c60203ba400e8 | 2.980622588 | 5.298055959 | 1.940460976 |
| dada98173a9d0b48e30800dbf3e16628 | 2.188217402 | 5.031460254 | 1.910200357 |
| 889b3824f9f3709fc805ea887f859e90 | 5.554251352 | 4.778779069 | 1.817323923 |
| ee705bbea43c68a47a9c2ba41efa2067 | 3.754255398 | -4.575396188 | 1.762208743 |
| ff289c73ac6541ac56707936f501a2cd | 5.147029566 | 3.549239697 | 1.395809726 |
| 026b68ad384fbb3f5bb0f6769b1b4fce | 54.30548866 | 3.209250229 | 1.277654889 |
| 637b9b3f4d1cbb1a10c07817619cdf69 | 545.8591622 | 0.304022069 | 0.124171812 |
| 2eeedead0f29786a7ec9cf13de5f5144 | 12.57189045 | 1.466025572 | 0.606778075 |
| ed04a525d9751330e39b5378f11571fc | 3.42971544 | 6.600908666 | 2.748840457 |

| stat | pvalue | padj | Kingdom | Phylum |
|---|---|---|---|---|
| 26.64751604 | 1.9123E-156 | 2.6199E-154 | Bacteria | Proteobacteria |
| 23.00617031 | 4.0434E-117 | 2.7697E-115 | Bacteria | Bacteroidetes |
| 19.99188273 | 6.4804E-89 | 2.95938E-87 | Bacteria | Proteobacteria |
| 16.29201404 | 1.12464E-59 | 3.85188E-58 | Bacteria | Bacteroidetes |
| 16.23693432 | 2.76396E-59 | 7.57325E-58 | Bacteria | Bacteroidetes |
| 11.72874827 | 9.07913E-32 | 2.07307E-30 | Bacteria | Bacteroidetes |
| 9.090014094 | 9.90262E-20 | 1.93808E-18 | Bacteria | Bacteroidetes |
| 7.998549545 | 1.25893E-15 | 2.15592E-14 | Bacteria | Proteobacteria |
| 7.464694738 | 8.34929E-14 | 1.27095E-12 | Bacteria | Proteobacteria |
| 7.179075272 | 7.01845E-13 | 9.61528E-12 | Bacteria | Proteobacteria |
| 6.944608795 | 3.79511E-12 | 4.72663E-11 | Bacteria | Proteobacteria |
| 6.854183651 | 7.17209E-12 | 8.18814E-11 | Bacteria | Proteobacteria |
| 6.682542703 | 2.34832E-11 | 2.47476E-10 | Bacteria | Bacteroidetes |
| 5.45728105 | 4.83481E-08 | 4.7312E-07 | Bacteria | Bacteroidetes |
| 5.35357961 | 8.62311E-08 | 7.87577E-07 | Bacteria | Proteobacteria |
| 5.136346294 | 2.80131E-07 | 2.39862E-06 | Bacteria | Proteobacteria |
| 5.11111958 | 3.20255E-07 | 2.58088E-06 | Bacteria | Proteobacteria |
| 5.079400543 | 3.78628E-07 | 2.88178E-06 | Bacteria | Proteobacteria |
| 4.755242876 | 1.98208E-06 | 1.35772E-05 | Bacteria | Proteobacteria |
| 4.759202158 | 1.9436E-06 | 1.35772E-05 | Bacteria | Proteobacteria |
| 4.705976844 | 2.52653E-06 | 1.64826E-05 | Bacteria | Proteobacteria |
| 4.600528376 | 4.21421E-06 | 2.6243E-05 | Bacteria | Bacteroidetes |
| 4.560554976 | 5.10186E-06 | 3.03893E-05 | Bacteria | Proteobacteria |
| 4.337053221 | 1.44406E-05 | 8.24316E-05 | Bacteria | Verrucomicrobia |
| 4.242383303 | 2.21159E-05 | 0.000121195 | Bacteria | Proteobacteria |
| 4.171117109 | 3.0311E-05 | 0.000159716 | Bacteria | Proteobacteria |
| 4.125900119 | 3.69288E-05 | 0.000187379 | Bacteria | Bacteroidetes |
| 4.071720514 | 4.66672E-05 | 0.000228336 | Bacteria | Proteobacteria |
| 4.002132135 | 6.27742E-05 | 0.000296554 | Bacteria | Proteobacteria |
| 3.778319418 | 0.00015789 | 0.000721032 | Bacteria | Proteobacteria |
| 3.380990709 | 0.00072225 | 0.003191878 | Bacteria | Proteobacteria |
| 3.350578602 | 0.000806429 | 0.003452525 | Bacteria | Bacteroidetes |
| 3.187838828 | 0.001433404 | 0.005950799 | Bacteria | Proteobacteria |
| 3.138134315 | 0.001700269 | 0.006851086 | Bacteria | Proteobacteria |
| 2.959100005 | 0.00308539 | 0.012077096 | Bacteria | Bacteroidetes |
| -2.854583755 | 0.004309327 | 0.016399382 | Bacteria | Bacteroidetes |
| 2.730307914 | 0.00632752 | 0.023428924 | Bacteria | Bacteroidetes |
| 2.633996081 | 0.008438646 | 0.030032199 | Bacteria | Bacteroidetes |
| 2.62956923 | 0.008549312 | 0.030032199 | Bacteria | Proteobacteria |
| -2.596398529 | 0.009420673 | 0.032265804 | Bacteria | Armatimonadetes |
| 2.542781893 | 0.010997385 | 0.03674736 | Bacteria | Proteobacteria |
| 2.511828708 | 0.012010737 | 0.039177881 | Bacteria | Proteobacteria |
| 2.448398421 | 0.014349288 | 0.0457175 | Bacteria | Proteobacteria |
| 2.416081977 | 0.015688529 | 0.048848374 | Bacteria | Proteobacteria |
| 2.401342955 | 0.016335019 | 0.049731058 | Bacteria | Bacteroidetes |

| Class | Order | Family | Genus |
|---|---|---|---|
| Gammaproteobacteria | Betaproteobacteriales | Burkholderiaceae | Undibacterium |
| Bacteroidia | Flavobacteriales | Flavobacteriaceae | Flavobacterium |
| Gammaproteobacteria | Pseudomonadales | Pseudomonadaceae | Pseudomonas |
| Bacteroidia | Flavobacteriales | Flavobacteriaceae | Flavobacterium |
| Bacteroidia | Flavobacteriales | Flavobacteriaceae | Flavobacterium |
| Bacteroidia | Cytophagales | Spirosomaceae | Flectobacillus |
| Bacteroidia | Flavobacteriales | Flavobacteriaceae | Flavobacterium |
| Gammaproteobacteria | Pseudomonadales | Moraxellaceae | Perlucidibaca |
| Gammaproteobacteria | Alteromonadales | Alteromonadaceae | Rheinheimera |
| Gammaproteobacteria | Pseudomonadales | Moraxellaceae | Perlucidibaca |
| Gammaproteobacteria | Betaproteobacteriales | Burkholderiaceae | Rhodoferax |
| Gammaproteobacteria | Betaproteobacteriales | Burkholderiaceae | Undibacterium |
| Bacteroidia | Flavobacteriales | Flavobacteriaceae | Flavobacterium |
| Bacteroidia | Flavobacteriales | Flavobacteriaceae | Flavobacterium |
| Gammaproteobacteria | Betaproteobacteriales | Burkholderiaceae | Pseudoduganella |
| Gammaproteobacteria | Alteromonadales | Alteromonadaceae | Rheinheimera |
| Gammaproteobacteria | Pseudomonadales | Pseudomonadaceae | Pseudomonas |
| Gammaproteobacteria | Pseudomonadales | Moraxellaceae | Perlucidibaca |
| Deltaproteobacteria | Bdellovibrionales | Bacteriovoracaceae | Bacteriovorax |
| Gammaproteobacteria | Pseudomonadales | Pseudomonadaceae | Pseudomonas |
| Gammaproteobacteria | Salinisphaerales | Solimonadaceae | Alkanibacter |
| Bacteroidia | Cytophagales | Spirosomaceae | Arcicella |
| Gammaproteobacteria | Betaproteobacteriales | Burkholderiaceae | Limnobacter |
| Verrucomicrobiae | Verrucomicrobiales | Rubritaleaceae | Luteolibacter |
| Gammaproteobacteria | Legionellales | Legionellaceae | Legionella |
| Gammaproteobacteria | Cellvibrionales | Cellvibrionaceae | Cellvibrio |
| Bacteroidia | Flavobacteriales | Flavobacteriaceae | Flavobacterium |
| Gammaproteobacteria | Betaproteobacteriales | Methylophilaceae | Methylotenera |
| Gammaproteobacteria | Pseudomonadales | Moraxellaceae | Acinetobacter |
| Gammaproteobacteria | Aeromonadales | Aeromonadaceae | Aeromonas |
| Gammaproteobacteria | Pseudomonadales | Moraxellaceae | Agitococcus |
| Bacteroidia | Flavobacteriales | Weeksellaceae | Chryseobacterium |
| Gammaproteobacteria | Betaproteobacteriales | Burkholderiaceae | Massilia |
| Gammaproteobacteria | Betaproteobacteriales | Rhodocyclaceae | Zoogloea |
| Bacteroidia | Flavobacteriales | Flavobacteriaceae | Flavobacterium |
| Bacteroidia | Flavobacteriales | Flavobacteriaceae | Flavobacterium |
| Bacteroidia | Flavobacteriales | Flavobacteriaceae | Flavobacterium |
| Bacteroidia | Cytophagales | Spirosomaceae | Flectobacillus |
| Gammaproteobacteria | Pseudomonadales | Moraxellaceae | Acinetobacter |
| Armatimonadia | Armatimonadales | Armatimonadaceae | Armatimonas |
| Gammaproteobacteria | Pseudomonadales | Moraxellaceae | Agitococcus |
| Gammaproteobacteria | Betaproteobacteriales | Burkholderiaceae | Undibacterium |
| Gammaproteobacteria | Pseudomonadales | Pseudomonadaceae | Pseudomonas |
| Gammaproteobacteria | Pseudomonadales | Pseudomonadaceae | Azotobacter |
| Bacteroidia | Flavobacteriales | Flavobacteriaceae | Flavobacterium |

| Species |
|---|
| NA |
| NA |
| NA |
| NA |
| NA |
| Flectobacillus fontis |
| NA |
| NA |
| NA |
| NA |
| NA |
| NA |
| NA |
| NA |
| NA |
| NA |
| NA |
| NA |
| NA |
| NA |
| NA |
| Arcicella rigui |
| NA |
| NA |
| NA |
| NA |
| NA |
| NA |
| NA |
| NA |
| NA |
| NA |
| NA |
| NA |
| NA |
| Flavobacterium columnare |
| Flavobacterium aquatile |
| NA |
| NA |
| NA |
| NA |
| NA |
| NA |
| Azotobacter chroococcum |
| NA |

### Ctl24_vs_Arg24

| ASV_ID | baseMean | log2FoldChange | lfcSE |
|---|---|---|---|
| 9ac9a274b64edbcb2b5e1e1c81743b51 | 10.35692861 | 24.39100137 | 1.168647024 |
| 88524108706087f7c75e79d6c10c9c61 | 10.96072901 | 25.56533863 | 1.635042651 |
| 2500422919f98bed627f3fd491e508a8 | 148.8388826 | 14.01293962 | 1.227282173 |
| f9c1c7de158ab4d555ec46bec26478f1 | 47.18641453 | 9.769647825 | 0.859757141 |
| 931bf16c32086439f09655b848d87174 | 369.3952111 | 5.922470942 | 0.52810199 |
| adf8882d5fd60155f03f9a52e900d32d | 118.8179065 | 6.487490634 | 0.588757823 |
| 04175f386c14319829ffb23d5c188b2a | 260.8060318 | 6.508335013 | 0.646671092 |
| 38c27ceaed634984c1225a82648cf571 | 280.8876036 | 9.447636333 | 1.067141648 |
| eb2625c1474f98f554b002fcd5d1088e | 62.89240942 | 6.417097592 | 0.729421888 |
| f34cdcd73e535507d29c29b19d01f658 | 71.11788206 | 7.922561051 | 0.967763432 |
| 98f49c75988e4d0fd97a180db1678fc8 | 48.8330818 | 8.314892192 | 1.029942437 |
| 4cfe5b5468150f8a54cf003ea4c15603 | 112.7343322 | 6.282001371 | 0.848628479 |
| 69179c4a6011d0c9d30036cf18626520 | 66.39688045 | 5.085097848 | 0.705654394 |
| d0fbc9aff818121519652ff493a83314 | 71.1331293 | 8.001709825 | 1.15432822 |
| a53888a61cc6766a2f54d75f5ac0cdef | 287.0563382 | 6.111312604 | 0.925817159 |
| c757ad670c5d796cb866c941c62c4e33 | 38.90084495 | 7.750211981 | 1.196103628 |
| f37fc1dccfeba0004cbeb4f1ad6083c6 | 24.922065 | 7.337687687 | 1.159924196 |
| bbf772e87bb1bd04ae18adfa03247a9f | 89.44846459 | 6.297691356 | 0.994221397 |
| 107273e73071282e4d0f4a0b46548da5 | 90.1483836 | 5.181703249 | 0.821415042 |
| b9fcd7d71b74853248517b892d03a745 | 55.31630097 | 9.025117555 | 1.446664199 |
| 2cd00ea627d63b6c55af5d1d65a6cd44 | 11.31504735 | 7.692920102 | 1.252085647 |
| 783869922c5a707e99ebebcabff8c5c6 | 84.96323871 | 4.783296336 | 0.778449294 |
| f89e77104762c7e39a9e0fb88c3ba6b3 | 20.45091495 | 7.423361172 | 1.241272364 |
| 113e5cb2dbb0b2086e230fc9d845310f | 54.74450093 | 7.570156422 | 1.290958255 |
| 3dbf66ee7707a9c91038139380697b07 | 46.42682375 | 5.663367606 | 0.971371843 |
| 4e6abee1bbb9afea6d2c733a33f76d8e | 8.00197725 | 7.169961058 | 1.265032087 |
| c0c6dea060404ea17972053bc9e98ae7 | 15.15068662 | 5.147854941 | 0.931163328 |
| 5648dccee530d68ceb3e4d7d22cf8756 | 58.40539467 | 2.95083383 | 0.559835751 |
| 709d785d9cfcbd7a92cfcc5fae6828f0 | 11.01312335 | 7.391906162 | 1.406250778 |
| 3badab07e5965e92e5954b746c192d50 | 18.06512302 | 8.102417316 | 1.553153114 |
| 8b1dba1a82d5149143975ddc2cc1089f | 91.55719942 | 4.567481076 | 0.875867799 |
| 6e6f306a337ba255a7f4a7f8469ff691 | 12.79955601 | 8.806286382 | 1.735248948 |
| 026b68ad384fbb3f5bb0f6769b1b4fce | 54.30548866 | 6.267320662 | 1.313616229 |
| 3ebe761bfb1238c87195d431f41bf976 | 52.38743114 | 2.956564036 | 0.653300165 |
| a8d4ddd7dbd400a101b2d3f1b3babaf2 | 17.37632741 | 5.773303558 | 1.300982353 |
| 095dcca0a1cacb655451100eb59c8ef1 | 12.63366984 | 4.317722981 | 1.080931516 |
| ff289c73ac6541ac56707936f501a2cd | 5.147029566 | 5.392951434 | 1.432665526 |
| c58754f3613677615cb80ce5013b088c | 17.02649803 | 4.782063027 | 1.282333597 |
| e416d0916760d2fc17b616e2ac3ad855 | 5.0021677 | 5.242178733 | 1.433275775 |
| a9d81e900e15e4fa6aad6ae109463454 | 5.864280573 | 6.443431153 | 1.776206864 |
| 861c357388b498ed9ef7ef6440b4dffa | 6.262005826 | 6.811509951 | 1.93026225 |
| 62b0bf21cce919c492e653ba630b9ea5 | 7.491337823 | 7.278585947 | 2.105280527 |
| 2c87d01f359e7d77cbdf931e0868986a | 15.62637297 | 4.987893532 | 1.495598233 |
| 36ecd054f5309a9658926a0926d7ee82 | 6.277950288 | 7.01943749 | 2.23460923 |
| 9ae0c31083aa43a5350972de7d54032b | 8.514490414 | 7.52157003 | 2.396311905 |
| 289811f7d4d770d3aafc67c5f8983af6 | 6.014606825 | 6.987982337 | 2.251459304 |
| c0f1efd28e2ac632b9821285c77fe072 | 16.91911017 | 2.905388859 | 0.951393978 |
| 41e6c0b55654af1d5cd503359fb77d18 | 4.701312972 | 6.582165915 | 2.226772349 |
| 892a0fc53b26d6a2c193d145b2464da7 | 3.215386764 | 4.883922467 | 1.65570513 |
| ee705bbea43c68a47a9c2ba41efa2067 | 3.754255398 | 4.98254297 | 1.804754729 |
| e9430eb4536e2d2615fd906f12c86712 | 3.851476078 | 6.375466095 | 2.355418016 |
| 7813794721c1199f44cbf1802167f1ec | 1.868182556 | 5.307668116 | 2.043508964 |
| 02a021a4a33ccd51f4b020422088397b | 7.740000228 | 6.837126863 | 2.66132767 |
| b2ffc94eb4b629641b6d0e124791e79e | 2.389958714 | 5.222645559 | 2.048089459 |

| | | | |
|---|---|---|---|
| e8386d3a307c208c4b9f0a756259cd6b | 17.3502534 | 2.927671137 | 1.201363882 |
| 8445499c647e84fa415763b46500feee | 3.435729604 | 5.808444639 | 2.465838183 |

| stat | pvalue | padj | Kingdom | Phylum |
|---|---|---|---|---|
| 20.8711449 | 9.79594E-97 | 1.43021E-94 | Bacteria | Bacteroidetes |
| 15.63588486 | 4.14674E-55 | 3.02712E-53 | Bacteria | Bacteroidetes |
| 11.41786293 | 3.40504E-30 | 1.65712E-28 | Bacteria | Proteobacteria |
| 11.36326453 | 6.37182E-30 | 2.32571E-28 | Bacteria | Bacteroidetes |
| 11.21463478 | 3.4561E-29 | 1.00918E-27 | Bacteria | Proteobacteria |
| 11.01894595 | 3.09664E-28 | 7.53515E-27 | Bacteria | Proteobacteria |
| 10.06436672 | 7.93967E-24 | 1.65599E-22 | Bacteria | Proteobacteria |
| 8.853216767 | 8.50323E-19 | 1.55184E-17 | Bacteria | Proteobacteria |
| 8.797511702 | 1.39883E-18 | 2.26922E-17 | Bacteria | Bacteroidetes |
| 8.18646457 | 2.69012E-16 | 3.92757E-15 | Bacteria | Proteobacteria |
| 8.073162043 | 6.85006E-16 | 9.0919E-15 | Bacteria | Proteobacteria |
| 7.402534235 | 1.3361E-13 | 1.62559E-12 | Bacteria | Proteobacteria |
| 7.206215804 | 5.75282E-13 | 6.46086E-12 | Bacteria | Proteobacteria |
| 6.931919089 | 4.15169E-12 | 4.32962E-11 | Bacteria | Proteobacteria |
| 6.600993021 | 4.08413E-11 | 3.97522E-10 | Bacteria | Proteobacteria |
| 6.47954893 | 9.19972E-11 | 8.39475E-10 | Bacteria | Bacteroidetes |
| 6.326006228 | 2.51588E-10 | 2.04066E-09 | Bacteria | Proteobacteria |
| 6.334294732 | 2.38429E-10 | 2.04066E-09 | Bacteria | Proteobacteria |
| 6.308264378 | 2.82182E-10 | 2.16835E-09 | Bacteria | Proteobacteria |
| 6.238571165 | 4.41586E-10 | 3.22358E-09 | Bacteria | Bacteroidetes |
| 6.144084569 | 8.0426E-10 | 5.33736E-09 | Bacteria | Bacteroidetes |
| 6.144647279 | 8.01414E-10 | 5.33736E-09 | Bacteria | Bacteroidetes |
| 5.980445056 | 2.22529E-09 | 1.41257E-08 | Bacteria | Bacteroidetes |
| 5.863982351 | 4.51896E-09 | 2.74903E-08 | Bacteria | Proteobacteria |
| 5.830277712 | 5.53352E-09 | 3.23158E-08 | Bacteria | Proteobacteria |
| 5.667809639 | 1.44635E-08 | 8.12178E-08 | Bacteria | Proteobacteria |
| 5.528412459 | 3.23142E-08 | 1.74736E-07 | Bacteria | Proteobacteria |
| 5.270892084 | 1.35762E-07 | 7.07903E-07 | Bacteria | Proteobacteria |
| 5.256463697 | 1.46852E-07 | 7.39323E-07 | Bacteria | Proteobacteria |
| 5.216753739 | 1.82086E-07 | 8.66624E-07 | Bacteria | Bacteroidetes |
| 5.214806481 | 1.84009E-07 | 8.66624E-07 | Bacteria | Proteobacteria |
| 5.074941201 | 3.87616E-07 | 1.7685E-06 | Bacteria | Proteobacteria |
| 4.771043873 | 1.83274E-06 | 8.10847E-06 | Bacteria | Proteobacteria |
| 4.525582871 | 6.02293E-06 | 2.58632E-05 | Bacteria | Proteobacteria |
| 4.437649399 | 9.09466E-06 | 3.79377E-05 | Bacteria | Proteobacteria |
| 3.994446381 | 6.48456E-05 | 0.000262985 | Bacteria | Proteobacteria |
| 3.764278078 | 0.000167031 | 0.000659095 | Bacteria | Proteobacteria |
| 3.729187973 | 0.000192098 | 0.00073806 | Bacteria | Bacteroidetes |
| 3.657480873 | 0.000254706 | 0.000953516 | Bacteria | Proteobacteria |
| 3.627635543 | 0.000286029 | 0.001044004 | Bacteria | Proteobacteria |
| 3.528800271 | 0.000417448 | 0.001486522 | Bacteria | Bacteroidetes |
| 3.4572998 | 0.000545617 | 0.00189667 | Bacteria | Proteobacteria |
| 3.335049094 | 0.000852843 | 0.002895701 | Bacteria | Bacteroidetes |
| 3.141237132 | 0.001682358 | 0.005503709 | Bacteria | Proteobacteria |
| 3.138810943 | 0.001696349 | 0.005503709 | Bacteria | Bacteroidetes |
| 3.103756895 | 0.001910803 | 0.006064722 | Bacteria | Bacteroidetes |
| 3.053823049 | 0.002259453 | 0.007018727 | Bacteria | Proteobacteria |
| 2.955922242 | 0.003117357 | 0.009475914 | Bacteria | Proteobacteria |
| 2.949753781 | 0.003180273 | 0.009475914 | Bacteria | Bacteroidetes |
| 2.760786765 | 0.005766231 | 0.016837395 | Bacteria | Armatimonadetes |
| 2.706723839 | 0.006795076 | 0.01945257 | Bacteria | Proteobacteria |
| 2.597330479 | 0.009395148 | 0.026378686 | Bacteria | Proteobacteria |
| 2.569066162 | 0.010197299 | 0.028090672 | Bacteria | Bacteroidetes |
| 2.550008514 | 0.010772029 | 0.029124374 | Bacteria | Proteobacteria |

| | | | | |
|---|---|---|---|---|
| 2.43695618 | 0.014811474 | 0.039317731 | Bacteria | Proteobacteria |
| 2.355566022 | 0.01849451 | 0.04821783 | Bacteria | Proteobacteria |

| Class | Order | Family | Genus |
|---|---|---|---|
| Bacteroidia | Flavobacteriales | Flavobacteriaceae | Flavobacterium |
| Bacteroidia | Flavobacteriales | Flavobacteriaceae | Flavobacterium |
| Gammaproteobacteria | Aeromonadales | Aeromonadaceae | Aeromonas |
| Bacteroidia | Flavobacteriales | Flavobacteriaceae | Flavobacterium |
| Gammaproteobacteria | Pseudomonadales | Moraxellaceae | Perlucidibaca |
| Gammaproteobacteria | Alteromonadales | Alteromonadaceae | Rheinheimera |
| Gammaproteobacteria | Alteromonadales | Alteromonadaceae | Rheinheimera |
| Gammaproteobacteria | Pseudomonadales | Pseudomonadaceae | Pseudomonas |
| Bacteroidia | Cytophagales | Spirosomaceae | Flectobacillus |
| Gammaproteobacteria | Betaproteobacteriales | Burkholderiaceae | Rhodoferax |
| Gammaproteobacteria | Betaproteobacteriales | Burkholderiaceae | Undibacterium |
| Gammaproteobacteria | Pseudomonadales | Moraxellaceae | Perlucidibaca |
| Gammaproteobacteria | Pseudomonadales | Moraxellaceae | Acinetobacter |
| Gammaproteobacteria | Betaproteobacteriales | Burkholderiaceae | Pseudoduganella |
| Gammaproteobacteria | Betaproteobacteriales | Chitinibacteraceae | Deefgea |
| Bacteroidia | Flavobacteriales | Flavobacteriaceae | Flavobacterium |
| Gammaproteobacteria | Betaproteobacteriales | Burkholderiaceae | Duganella |
| Gammaproteobacteria | Betaproteobacteriales | Burkholderiaceae | Undibacterium |
| Gammaproteobacteria | Aeromonadales | Aeromonadaceae | Aeromonas |
| Bacteroidia | Flavobacteriales | Flavobacteriaceae | Flavobacterium |
| Bacteroidia | Cytophagales | Spirosomaceae | Arcicella |
| Bacteroidia | Flavobacteriales | Flavobacteriaceae | Flavobacterium |
| Bacteroidia | Flavobacteriales | Flavobacteriaceae | Flavobacterium |
| Gammaproteobacteria | Pseudomonadales | Pseudomonadaceae | Pseudomonas |
| Gammaproteobacteria | Pseudomonadales | Pseudomonadaceae | Pseudomonas |
| Gammaproteobacteria | Legionellales | Legionellaceae | Legionella |
| Gammaproteobacteria | Betaproteobacteriales | Burkholderiaceae | Limnobacter |
| Gammaproteobacteria | Pseudomonadales | Pseudomonadaceae | Pseudomonas |
| Deltaproteobacteria | Bdellovibrionales | Bacteriovoracaceae | Bacteriovorax |
| Bacteroidia | Flavobacteriales | Flavobacteriaceae | Flavobacterium |
| Gammaproteobacteria | Pseudomonadales | Moraxellaceae | Perlucidibaca |
| Gammaproteobacteria | Cellvibrionales | Cellvibrionaceae | Cellvibrio |
| Gammaproteobacteria | Betaproteobacteriales | Burkholderiaceae | Undibacterium |
| Gammaproteobacteria | Pseudomonadales | Moraxellaceae | Acinetobacter |
| Gammaproteobacteria | Betaproteobacteriales | Burkholderiaceae | Massilia |
| Gammaproteobacteria | Pseudomonadales | Moraxellaceae | Agitococcus |
| Gammaproteobacteria | Pseudomonadales | Moraxellaceae | Agitococcus |
| Bacteroidia | Cytophagales | Spirosomaceae | Emticicia |
| Gammaproteobacteria | Betaproteobacteriales | Methylophilaceae | Methylotenera |
| Gammaproteobacteria | Betaproteobacteriales | Burkholderiaceae | Undibacterium |
| Bacteroidia | Sphingobacteriales | Sphingobacteriaceae | Pedobacter |
| Gammaproteobacteria | Betaproteobacteriales | Burkholderiaceae | Undibacterium |
| Bacteroidia | Flavobacteriales | Flavobacteriaceae | Flavobacterium |
| Alphaproteobacteria | Rhizobiales | Rhizobiaceae | Shinella |
| Bacteroidia | Flavobacteriales | Flavobacteriaceae | Flavobacterium |
| Bacteroidia | Flavobacteriales | Weeksellaceae | Chryseobacterium |
| Gammaproteobacteria | Salinisphaerales | Solimonadaceae | Alkanibacter |
| Alphaproteobacteria | Caulobacterales | Caulobacteraceae | Brevundimonas |
| Bacteroidia | Flavobacteriales | Weeksellaceae | Chryseobacterium |
| Armatimonadia | Armatimonadales | Armatimonadaceae | Armatimonas |
| Deltaproteobacteria | Myxococcales | Polyangiaceae | Pajaroellobacter |
| Gammaproteobacteria | Legionellales | Legionellaceae | Legionella |
| Bacteroidia | Flavobacteriales | Flavobacteriaceae | Flavobacterium |
| Gammaproteobacteria | Betaproteobacteriales | Burkholderiaceae | Duganella |

| | | | |
|---|---|---|---|
| Gammaproteobacteria | Pseudomonadales | Moraxellaceae | Acinetobacter |
| Alphaproteobacteria | Rhodobacterales | Rhodobacteraceae | Pseudorhodobacter |

| Species |
| --- |
| NA |
| NA |
| NA |
| NA |
| NA |
| NA |
| NA |
| NA |
| Flectobacillus fontis |
| NA |
| NA |
| NA |
| NA |
| NA |
| NA |
| NA |
| NA |
| NA |
| NA |
| NA |
| Arcicella rigui |
| NA |
| NA |
| NA |
| NA |
| NA |
| NA |
| NA |
| NA |
| NA |
| NA |
| NA |
| NA |
| NA |
| NA |
| NA |
| NA |
| NA |
| NA |
| NA |
| NA |
| NA |
| NA |
| NA |
| NA |
| NA |
| NA |
| NA |
| NA |
| NA |
| NA |
| NA |
| NA |
| NA |

NA
NA

| ASV_ID | baseMean | log2FoldChange | lfcSE |
|---|---|---|---|
| 98f49c75988e4d0fd97a180db1678fc8 | 48.8330818 | 20.28014948 | 1.066785487 |
| 113e5cb2dbb0b2086e230fc9d845310f | 54.74450093 | 21.61422336 | 1.323985911 |
| b9fcd7d71b74853248517b892d03a745 | 55.31630097 | 17.87072721 | 1.478200318 |
| 931bf16c32086439f09655b848d87174 | 369.3952111 | -4.333524917 | 0.531857704 |
| 31c862c1b2e0ffdd72b11a0b50c208f6 | 13.13589934 | -7.140462999 | 0.890352155 |
| 2500422919f98bed627f3fd491e508a8 | 148.8388826 | -8.335277737 | 1.236762012 |
| 4cfe5b5468150f8a54cf003ea4c15603 | 112.7343322 | -5.711742641 | 0.85403106 |
| 8b1dba1a82d5149143975ddc2cc1089f | 91.55719942 | 4.679172648 | 0.90514734 |
| eb2625c1474f98f554b002fcd5d1088e | 62.89240942 | 3.258172072 | 0.773802388 |
| adf8882d5fd60155f03f9a52e900d32d | 118.8179065 | -2.068284845 | 0.594285468 |
| 65d43491988bfe557da4d86a5ba25dae | 40.85217081 | 2.054421436 | 0.59617631 |
| a53888a61cc6766a2f54d75f5ac0cdef | 287.0563382 | -3.003599561 | 0.9325886 |
| e371861e8798027b6e72500f333c78cc | 3.441387683 | -5.990071062 | 1.911553321 |

| stat | pvalue | padj | | Kingdom | Phylum |
|---|---|---|---|---|---|
| 19.01052248 | 1.39556E-80 | 5.23333E-78 | Bacteria | Proteobacteria |
| 16.32511583 | 6.54164E-60 | 1.22656E-57 | Bacteria | Proteobacteria |
| 12.08951655 | 1.19991E-33 | 1.49988E-31 | Bacteria | Bacteroidetes |
| -8.147902877 | 3.7029E-16 | 3.47146E-14 | Bacteria | Proteobacteria |
| -8.01981885 | 1.05901E-15 | 7.94259E-14 | Bacteria | Firmicutes |
| -6.739597155 | 1.58826E-11 | 9.92665E-10 | Bacteria | Proteobacteria |
| -6.687979992 | 2.26272E-11 | 1.21217E-09 | Bacteria | Proteobacteria |
| 5.169514885 | 2.34702E-07 | 1.10017E-05 | Bacteria | Proteobacteria |
| 4.210599663 | 2.54694E-05 | 0.001061224 | Bacteria | Bacteroidetes |
| -3.480288442 | 0.000500874 | 0.018782783 | Bacteria | Proteobacteria |
| 3.44599643 | 0.000568958 | 0.019396291 | Bacteria | Firmicutes |
| -3.220712287 | 0.001278724 | 0.039960141 | Bacteria | Proteobacteria |
| -3.133614426 | 0.001726676 | 0.049807956 | Bacteria | Bacteroidetes |

| Class | Order | Family | Genus |
|---|---|---|---|
| Gammaproteobacteria | Betaproteobacteriales | Burkholderiaceae | Undibacterium |
| Gammaproteobacteria | Pseudomonadales | Pseudomonadaceae | Pseudomonas |
| Bacteroidia | Flavobacteriales | Flavobacteriaceae | Flavobacterium |
| Gammaproteobacteria | Pseudomonadales | Moraxellaceae | Perlucidibaca |
| Bacilli | Bacillales | Bacillaceae | Bacillus |
| Gammaproteobacteria | Aeromonadales | Aeromonadaceae | Aeromonas |
| Gammaproteobacteria | Pseudomonadales | Moraxellaceae | Perlucidibaca |
| Gammaproteobacteria | Pseudomonadales | Moraxellaceae | Perlucidibaca |
| Bacteroidia | Cytophagales | Spirosomaceae | Flectobacillus |
| Gammaproteobacteria | Alteromonadales | Alteromonadaceae | Rheinheimera |
| Bacilli | Bacillales | Staphylococcaceae | Staphylococcus |
| Gammaproteobacteria | Betaproteobacteriales | Chitinibacteraceae | Deefgea |
| Bacteroidia | Flavobacteriales | Flavobacteriaceae | Flavobacterium |

| Species |
|---|
| NA |
| NA |
| NA |
| NA |
| NA |
| NA |
| NA |
| NA |
| Flectobacillus fontis |
| NA |
| NA |
| NA |
| Flavobacterium columnare |

### Arg12_vs_Arg24

| ASV_ID | baseMean | log2FoldChange | lfcSE |
|---|---|---|---|
| 31c862c1b2e0ffdd72b11a0b50c208f6 | 13.13589934 | -5.76163132 | 0.854160072 |
| 931bf16c32086439f09655b848d87174 | 369.3952111 | -2.442755181 | 0.500089563 |
| 2500422919f98bed627f3fd491e508a8 | 148.8388826 | 5.528884575 | 1.164884415 |
| 107273e73071282e4d0f4a0b46548da5 | 90.1483836 | 3.792185109 | 0.794505409 |
| 4cfe5b5468150f8a54cf003ea4c15603 | 112.7343322 | -3.487309887 | 0.793049695 |
| 8b1dba1a82d5149143975ddc2cc1089 | 91.55719942 | 2.942712958 | 0.847886253 |
| 38c27ceaed634984c1225a82648cf571 | 280.8876036 | 3.525361989 | 1.022459854 |
| f9c1c7de158ab4d555ec46bec26478f1 | 47.18641453 | 2.583494777 | 0.797144297 |
| 9ae0c31083aa43a5350972de7d54032 | 8.514490414 | 7.431077099 | 2.32285914 |
| 69179c4a6011d0c9d30036cf18626520 | 66.39688045 | 2.157085019 | 0.679456748 |
| 52abfae86a69569ae491695349cca6f8 | 6.926038692 | -6.779943534 | 2.173960693 |
| f37fc1dccfeba0004cbeb4f1ad6083c6 | 24.922065 | 3.436758108 | 1.107712269 |
| 113e5cb2dbb0b2086e230fc9d845310f | 54.74450093 | 3.704179235 | 1.240183314 |
| ee705bbea43c68a47a9c2ba41efa2067 | 3.754255398 | 5.079575151 | 1.749641998 |
| e9430eb4536e2d2615fd906f12c86712 | 3.851476078 | 6.284974449 | 2.283222677 |
| b9fcd7d71b74853248517b892d03a745 | 55.31630097 | 3.785295617 | 1.385626466 |

| stat | pvalue | padj | Kingdom | Phylum |
|---|---|---|---|---|
| -6.745376553 | 1.52631E-11 | 1.8163E-09 | Bacteria | Firmicutes |
| -4.884635398 | 1.0362E-06 | 6.16351E-05 | Bacteria | Proteobacteria |
| 4.746294572 | 2.07177E-06 | 6.16351E-05 | Bacteria | Proteobacteria |
| 4.773013581 | 1.8149E-06 | 6.16351E-05 | Bacteria | Proteobacteria |
| -4.397340932 | 1.09585E-05 | 0.000260813 | Bacteria | Proteobacteria |
| 3.470645911 | 0.000519208 | 0.009603586 | Bacteria | Proteobacteria |
| 3.447922162 | 0.000564917 | 0.009603586 | Bacteria | Proteobacteria |
| 3.240937413 | 0.001191373 | 0.017721679 | Bacteria | Bacteroidetes |
| 3.199107931 | 0.001378535 | 0.017847748 | Bacteria | Bacteroidetes |
| 3.174720136 | 0.001499811 | 0.017847748 | Bacteria | Proteobacteria |
| -3.118705668 | 0.001816473 | 0.019024755 | Bacteria | Proteobacteria |
| 3.102572937 | 0.001918463 | 0.019024755 | Bacteria | Proteobacteria |
| 2.986799767 | 0.002819143 | 0.025806001 | Bacteria | Proteobacteria |
| 2.903208289 | 0.003693609 | 0.031395676 | Bacteria | Armatimonadetes |
| 2.752676956 | 0.005911019 | 0.046844109 | Bacteria | Proteobacteria |
| 2.73182976 | 0.006298368 | 0.046844109 | Bacteria | Bacteroidetes |

| Class | Order | Family | Genus | Species |
|---|---|---|---|---|
| Bacilli | Bacillales | Bacillaceae | Bacillus | NA |
| Gammaproteobacteria | Pseudomonadales | Moraxellaceae | Perlucidibaca | NA |
| Gammaproteobacteria | Aeromonadales | Aeromonadaceae | Aeromonas | NA |
| Gammaproteobacteria | Aeromonadales | Aeromonadaceae | Aeromonas | NA |
| Gammaproteobacteria | Pseudomonadales | Moraxellaceae | Perlucidibaca | NA |
| Gammaproteobacteria | Pseudomonadales | Moraxellaceae | Perlucidibaca | NA |
| Gammaproteobacteria | Pseudomonadales | Pseudomonadaceae | Pseudomonas | NA |
| Bacteroidia | Flavobacteriales | Flavobacteriaceae | Flavobacterium | NA |
| Bacteroidia | Flavobacteriales | Flavobacteriaceae | Flavobacterium | NA |
| Gammaproteobacteria | Pseudomonadales | Moraxellaceae | Acinetobacter | NA |
| Gammaproteobacteria | Betaproteobacteriales | Rhodocyclaceae | Zoogloea | NA |
| Gammaproteobacteria | Betaproteobacteriales | Burkholderiaceae | Duganella | NA |
| Gammaproteobacteria | Pseudomonadales | Pseudomonadaceae | Pseudomonas | NA |
| Armatimonadia | Armatimonadales | Armatimonadaceae | Armatimonas | NA |
| Deltaproteobacteria | Myxococcales | Polyangiaceae | Pajaroellobacter | NA |
| Bacteroidia | Flavobacteriales | Flavobacteriaceae | Flavobacterium | NA |
