## Supplementary Table 3 for "Circadian dynamics of the teleost skin immune-microbiome interface"

| Group 1 | Group 2 | pseudo-F | p-value | q-value |
|---|---|---|---|---|
| Argulus_12 | Argulus_24 | 1.927441837 | 0.004 | 0.004 |
| Argulus_12 | Control_12 | 3.167869012 | 0.001 | 0.001111111 |
| Argulus_12 | Control_24 | 4.887142676 | 0.001 | 0.001111111 |
| Argulus_12 | Water | 14.804493 | 0.001 | 0.001111111 |
| Argulus_24 | Control_12 | 6.276399245 | 0.001 | 0.001111111 |
| Argulus_24 | Control_24 | 7.37383445 | 0.001 | 0.001111111 |
| Argulus_24 | Water | 15.17727828 | 0.001 | 0.001111111 |
| Control_12 | Control_24 | 3.307888287 | 0.001 | 0.001111111 |
| Control_12 | Water | 17.5535308 | 0.001 | 0.001111111 |
| Control_24 | Water | 19.88133954 | 0.001 | 0.001111111 |
