## Supplementary Table 2 for "Circadian dynamics of the teleost skin immune-microbiome interface"

| Group 1 | Group 2 | H | p-value |
|---|---|---|---|
| Argulus_12 | Argulus_24 | 0.052886766 | 0.818114126 |
| Argulus_12 | Control_12 | 0.509925111 | 0.475171277 |
| Argulus_12 | Control_24 | 13.50547798 | 0.000237868 |
| Argulus_12 | Water | 32.64 | 1.10909E-08 |
| Argulus_24 | Control_12 | 0.10845712 | 0.741907566 |
| Argulus_24 | Control_24 | 14.01007431 | 0.000181834 |
| Argulus_24 | Water | 31.85643414 | 1.65999E-08 |
| Control_12 | Control_24 | 12.92404919 | 0.000324387 |
| Control_12 | Water | 34.22222222 | 4.91643E-09 |
| Control_24 | Water | 33.81818182 | 6.05107E-09 |

| q-value |
| --- |
| 0.818114126 |
| 0.593964096 |
| 0.000396447 |
| 3.69695E-08 |
| 0.818114126 |
| 0.000363668 |
| 4.14997E-08 |
| 0.000463411 |
| 3.02553E-08 |
| 3.02553E-08 |
