## Supplementary Table 1 for "Circadian dynamics of the teleost skin immune-microbiome interface"

| Gene | Genbank Accession | RefSeq Gene ID | |
|---|---|---|---|
| clock1a | GU228520 | 100135915 | Clock genes |
| clock1b | GU228521 | 110521676 | |
| clock3 | GU228522 | 110533857 | |
| bmal1 | GQ489026 | 100499618 | |
| bmal2 | CX717649 | 110499965 | |
| per1 | AF228695 | 100135905 | |
| per2 | | 110508801 | |
| cry1 | CA383214 | 110522881 | |
| cry2 | | 110499866 | |
| reverbb | AF342943 | 100135954 | |
| aanat2 | AF106006 | 100135881 | |
| rora | | 110520931 | |
| csnk1d | | 110499233 | |
| timeless | | 110494196 | |
| il1b | AJ223954 | 100136024 | Immune genes |
| il4 | FN820501 | 100653462 | |
| il6 | DQ866150 | 100136689 | |
| il10 | AB118099 | 100136835 | |
| il17a | AJ580842 | 100136642 | |
| tnfa | AJ277604 | 100136034 | |
| nos2 | AJ295230 | 100136036 | |
| ifng | AJ616215 | 100136643 | |
| tgfb | AJ007836 | 100136774 | |
| tcrb | AJ517930 | 110504270 | |
| igt | AY870263 | | |
| igm | X65261 | | |
| igd | AY870262 | | |
| tbx21 | FM863825 | 100500940 | |
| gata3 | FM863826 | 100500939 | |
| foxp3b | FM883711 | 100653438 | |
| rory | FM883712 | 100528059 | |
| cd4 | AY973030 | 100136285 | |
| cd8a | AF178053 | 100135889 | |
| cath1 | AY594646 | 100136204 | |
| cath2 | AY542963 | 100136187 | |
| hamp | AF281354 | 100135935 | |
| tlr2 | HE979560 | 100750259 | |
| tlr9 | EU627195 | 100170212 | |
| tlr22 | AJ628348 | 100136113 | |
| mhcii | AY273808 | 100500791 | |
| chi | AJ535688 | 100136076 | |
| c3 | L24433 | 110489027 | |
| crf | AY049980 | 100135941 | Corti |
| pomc | X69808 | 100136771 | |
| rplp0 | | 110494133 | Controls |
| polr2i | | 100305160 | |
| hprt1 | | 110504699 | |
| polr1b | | 110526594 | |
