## Supplementary Datafile 1 for "Circadian dynamics of the teleost skin immune-microbiome interface"

| **ID** | **Desc** |
|---|---|
| *POLYAMINSYN3-PWY* | polyamine biosynthesis II |
| *PWY-2942* | L-lysine biosynthesis III |
| PWY-4361 | S-methyl-5-thio-α-D-ribose 1-phosphate degradation I |
| PWY-7527 | L-methionine salvage cycle III |
| *PWY-5097* | L-lysine biosynthesis VI |
| PWY-6165 | chorismate biosynthesis II (archaea) |
| PWY0-1241 | ADP-L-glycero-β-D-manno-heptose biosynthesis |
| *P241-PWY* | coenzyme B biosynthesis |
| PWY-6654 | phosphopantothenate biosynthesis III (archaebacteria) |
| PWY-5920 | heme b biosynthesis from glycine |
| *NAD-BIOSYNTHESIS-II* | NAD salvage pathway III (to nicotinamide riboside) |
| PWY-7221 | guanosine ribonucleotides de novo biosynthesis |
| PWY-7228 | guanosine nucleotides de novo biosynthesis I |
| PWY-6609 | adenine and adenosine salvage III |
| PWY-6125 | guanosine nucleotides de novo biosynthesis II |
| PWY-6174 | mevalonate pathway II (haloarchaea) |
| *PWY-7391* | isoprene biosynthesis II |
| *PWY-7431* | aromatic biogenic amine degradation (bacteria) |
| PWY-5022 | 4-aminobutanoate degradation V |
| PWY-5181 | toluene degradation III (aerobic) (via p-cresol) |
| *CATECHOL-ORTHO-CLEAVAGE-PWY* | catechol degradation to β-ketoadipate |
| *PWY-5417* | catechol degradation III (ortho-cleavage pathway) |
| *PWY-5431* | aromatic compounds degradation via β-ketoadipate |
| *PWY-5183* | aerobic toluene degradation |
| CODH-PWY | reductive acetyl coenzyme A pathway I (homoacetogenic |
| PWY-6572 | chondroitin sulfate degradation I (bacterial) |
| PWY-3801 | sucrose degradation II (sucrose synthase) |
| *PWY-6737* | starch degradation V |
| *PWY-6944* | androstenedione degradation |
| LIPASYN-PWY | phospholipases |
| *PWY0-1296* | purine ribonucleosides degradation |
| PWY0-1297 | purine deoxyribonucleosides degradation |
| PWY-6353 | purine nucleotides degradation II (aerobic) |
| *P221-PWY* | octane oxidation |
| *PWY-7237* | myo-, chiro- and scyllo-inositol degradation |
| *PWY0-1338* | *polymyxin resistance* |
| *GLYCOLYSIS* | glycolysis I (from glucose 6-phosphate) |
| CALVIN-PWY | Calvin-Benson-Bassham cycle |
| *P105-PWY* | TCA cycle IV (2-oxoglutarate decarboxylase) |
| *PWY-6969* | TCA cycle V (2-oxoglutarate:ferredoxin oxidoreductase) |

| eJTK_BF | Peak | AvProp | Superclass |
|---|---|---|---|
| 0.01458275 | 12.88595 | 0.003084 | Biosynthesis |
| 0.00510748 | 2.627986 | 0.00547015 | Biosynthesis |
| 0.01252049 | 6.427066 | 1.66E-05 | Biosynthesis |
| 0.0127468 | 6.325732 | 2.57E-05 | Biosynthesis |
| 0.02253547 | 2.782936 | 0.00530019 | Biosynthesis |
| 0.003653 | 2.972316 | 2.38E-05 | Biosynthesis |
| 0.04602597 | 14.17549 | 0.00266054 | Biosynthesis |
| 0.00022438 | 23.21666 | 1.87E-06 | Biosynthesis |
| 0.00321305 | 2.965556 | 1.78E-05 | Biosynthesis |
| 0.02084569 | 0.2532651 | 0.00113167 | Biosynthesis |
| 0.02931922 | 21.53345 | 0.00166294 | Biosynthesis |
| 0.02150775 | 0.2965713 | 0.00480853 | Biosynthesis |
| 0.03818027 | 0.9234364 | 0.0052048 | Biosynthesis |
| 0.04602597 | 1.897756 | 0.00132033 | Biosynthesis |
| 0.04939549 | 0.837666 | 0.00508615 | Biosynthesis |
| 0.0016419 | 1.666899 | 1.13E-05 | Biosynthesis |
| 0.0337347 | 2.790154 | 2.04E-05 | Biosynthesis |
| 0.01711526 | 15.31357 | 0.0024523 | Degradation/Utilization/Assimilation |
| 0.04124837 | 13.75506 | 0.00400379 | Degradation/Utilization/Assimilation |
| 0.02698494 | 15.82952 | 0.00345912 | Degradation/Utilization/Assimilation |
| 0.0164469 | 14.8543475 | 0.00274011 | Degradation/Utilization/Assimilation |
| 0.01711526 | 15.0864067 | 0.00272625 | Degradation/Utilization/Assimilation |
| 0.01711526 | 15.0863985 | 0.00272625 | Degradation/Utilization/Assimilation |
| 0.02684517 | 21.53345 | 0.00012944 | Degradation/Utilization/Assimilation |
| 0.03977813 | 21.01987 | 9.81E-06 | Degradation/Utilization/Assimilation |
| 0.0013237 | 5.314672 | 2.31E-06 | Degradation/Utilization/Assimilation |
| 0.00358437 | 2.359213 | 1.16E-05 | Degradation/Utilization/Assimilation |
| 0.03427187 | 13.9716708 | 0.00441379 | Degradation/Utilization/Assimilation |
| 0.0259928 | 21.85183 | 8.86E-06 | Degradation/Utilization/Assimilation |
| 0.04823662 | 5.314672 | 1.17E-05 | Degradation/Utilization/Assimilation |
| 0.03427187 | 1.388168 | 0.0008885 | Degradation/Utilization/Assimilation |
| 0.03975825 | 1.931602 | 0.0010104 | Degradation/Utilization/Assimilation |
| 0.04124837 | 1.619075 | 0.00216428 | Degradation/Utilization/Assimilation |
| 0.03301205 | 15.7577179 | 0.00372876 | Degradation/Utilization/Assimilation |
| 0.02948037 | 16.4732668 | 0.0050147 | Degradation/Utilization/Assimilation |
| *0.02167555* | *14.88154* | *0.00235998* | *Detoxification* |
| 0.02530526 | 1.35234 | 0.00421532 | Generation of Precursor Metabolites |
| 0.01868302 | 0.5373235 | 0.00504978 | Generation of Precursor Metabolites |
| 0.00488907 | 1.918829 | 0.00485251 | Generation of Precursor Metabolites |
| 0.00634214 | 1.991988 | 0.00486768 | Generation of Precursor Metabolites |

| **Subclass** |
| --- |
| Amine and Polyamine Biosynthesis |
| Amino Acid Biosynthesis |
| Amino Acid Biosynthesis |
| Amino Acid Biosynthesis |
| Amino Acid Biosynthesis |
| Aromatic Compound Biosynthesis |
| Carbohydrate Biosynthesis |
| Cofactor, Carrier, and Vitamin Biosynthesis |
| Cofactor, Carrier, and Vitamin Biosynthesis |
| Cofactor, Carrier, and Vitamin Biosynthesis |
| Cofactor, Carrier, and Vitamin Biosynthesis |
| Nucleoside and Nucleotide Biosynthesis |
| Nucleoside and Nucleotide Biosynthesis |
| Nucleoside and Nucleotide Biosynthesis |
| Nucleoside and Nucleotide Biosynthesis |
| Secondary Metabolite Biosynthesis |
| Secondary Metabolite Biosynthesis |
| Amine and Polyamine Degradation |
| Amine and Polyamine Degradation |
| Amino Acid Degradation |
| Aromatic Compound Degradation |
| Aromatic Compound Degradation |
| Aromatic Compound Degradation |
| Aromatic Compound Degradation |
| C1 Compound Utilization and Assimilation |
| Carbohydrate Degradation |
| Carbohydrate Degradation |
| Carbohydrate Degradation |
| Fatty Acid and Lipid Degradation |
| Fatty Acid and Lipid Degradation |
| Nucleoside and Nucleotide Degradation |
| Nucleoside and Nucleotide Degradation |
| Nucleoside and Nucleotide Degradation |
| Other |
| Secondary Metabolite Degradation |
| *Antibiotic Resistance* |
| Glycolysis |
| Photosynthesis |
| TCA Cycle |
| TCA Cycle |

| ID | Desc |
|---|---|
| PWY-5189 | tetrapyrrole biosynthesis II (from glycine) |
| PWY-6897 | thiamine diphosphate salvage II |
| COA-PWY | coenzyme A biosynthesis I (prokaryotic) |
| PWY-5188 | tetrapyrrole biosynthesis I (from glutamate) |
| PWY-841 | purine nucleotides de novo biosynthesis I |
| PWY-7234 | inosine-5'-phosphate biosynthesis III |
| PWY-6123 | inosine-5'-phosphate biosynthesis I |
| PWY-5103 | L-isoleucine biosynthesis III |
| POLYISOPRENSYN-PWY | polyisoprenoid biosynthesis (E. coli) |
| PWY-4361 | S-methyl-5-thio-α-D-ribose 1-phosphate de |
| PWY-7527 | L-methionine salvage cycle III |
| PWY-6126 | adenosine nucleotides de novo biosynthesi |
| PWY-5686 | UMP biosynthesis I |
| PWY-7229 | adenosine nucleotides de novo biosynthesi |
| HISTSYN-PWY | L-histidine biosynthesis |
| PWY-7219 | adenosine ribonucleotides de novo biosynt |
| P261-PWY | coenzyme M biosynthesis I |
| PWY0-162 | pyrimidine ribonucleotides de novo biosynt |
| PWY0-1479 | tRNA processing |
| BRANCHED-CHAIN-AA-SYN-PWY | branched chain amino acid biosynthesis |
| PWY-5918 | heme b biosynthesis from glutamate |
| ILEUSYN-PWY | L-isoleucine biosynthesis I (from threonine) |
| VALSYN-PWY | L-valine biosynthesis |
| PWY-5101 | L-isoleucine biosynthesis II |
| PWY-6174 | mevalonate pathway II (haloarchaea) |
| PWY-3781 | aerobic respiration I (cytochrome c) |
| HEMESYN2-PWY | heme b biosynthesis II (oxygen-independer |
| PWY-5183 | aerobic toluene degradation |
| PWY0-845 | pyridoxal 5'-phosphate biosynthesis and sa |
| *PWY-7391* | *isoprene biosynthesis II* |
| GLUCONEO-PWY | gluconeogenesis I |
| PWY-6700 | queuosine biosynthesis I (de novo) |
| PWY-5913 | partial TCA cycle (obligate autotrophs) |
| TRNA-CHARGING-PWY | tRNA charging |
| PEPTIDOGLYCANSYN-PWY | peptidoglycan biosynthesis I (meso-diamin |
| TRPSYN-PWY | L-tryptophan biosynthesis |
| PWY-3001 | L-isoleucine biosynthesis I |
| HEME-BIOSYNTHESIS-II | heme b biosynthesis I (aerobic) |
| PWY-6125 | guanosine nucleotides de novo biosynthesi |
| PWY-1882 | C1 compounds oxidation to CO2 |
| PWY-2942 | L-lysine biosynthesis III |
| PWY-7220 | adenosine deoxyribonucleotides de novo b |
| PWY-7222 | guanosine deoxyribonucleotides de novo b |
| PWY-7221 | guanosine ribonucleotides de novo biosynt |
| HSERMETANA-PWY | L-methionine biosynthesis III |
| PWY-5097 | L-lysine biosynthesis VI |
| PWY-7228 | guanosine nucleotides de novo biosynthesi |
| PWY-6387 | UDP-N-acetylmuramoyl-pentapeptide biosy |
| *PWY-5531* | *3,8-divinyl-chlorophyllide a biosynthesis II (* |
| *PWY-7159* | |
| PWY-5198 | factor 420 biosynthesis II (mycobacteria) |
| PWY-7208 | pyrimidine nucleobases salvage |

| | |
|---|---|
| PWY-6969 | TCA cycle V (2-oxoglutarate:ferredoxin oxi |
| CALVIN-PWY | Calvin-Benson-Bassham cycle |
| PWY-5022 | 4-aminobutanoate degradation V |
| PWY0-1241 | ADP-L-glycero-β-D-manno-heptose biosyn |
| PWY0-1061 | L-alanine biosynthesis |
| P105-PWY | TCA cycle IV (2-oxoglutarate decarboxylas |
| PWY-6386 | UDP-N-acetylmuramoyl-pentapeptide bios |
| GLYCOLYSIS | glycolysis I (from glucose 6-phosphate) |
| PWY0-1338 | polymyxin resistance |
| PWY-6385 | peptidoglycan biosynthesis III (mycobacteri |
| *UDPNAGSYN-PWY* | *UDP-N-acetyl-D-glucosamine biosynthesis* |
| PWY-5505 | L-glutamate and L-glutamine biosynthesis |
| PWY-7237 | myo-, chiro- and scyllo-inositol degradation |
| *PWY-6944* | |
| *PWY-6165* | |
| *NAGLIPASYN-PWY* | |
| *PWY-6654* | |
| CATECHOL-ORTHO-CLEAVAGE-PWY | catechol degradation to β-ketoadipate |
| PWY-7094 | fatty acid salvage |
| POLYAMINSYN3-PWY | polyamine biosynthesis II |
| *CHLOROPHYLL-SYN* | |
| P184-PWY | protocatechuate degradation I (meta-cleav |
| PANTOSYN-PWY | coenzyme A biosynthesis I (bacteria) |
| P562-PWY | myo-inositol degradation I |
| PWY-5417 | catechol degradation III (ortho-cleavage pa |
| PWY-5431 | aromatic compounds degradation via β-ket |
| PWY-7211 | pyrimidine deoxyribonucleotides de novo b |
| PENTOSE-P-PWY | pentose phosphate pathway |
| PWY0-1533 | methylphosphonate degradation I |
| PWY-7431 | aromatic biogenic amine degradation (bact |
| PWY-7197 | pyrimidine deoxyribonucleotide phosphoryl |
| PWY-7376 | cob(II)yrinate a,c-diamide biosynthesis II (l |
| PROTOCATECHUATE-ORTHO-CLEAVAGE-PWY | protocatechuate degradation II (ortho-clea |
| HISDEG-PWY | L-histidine degradation I |
| PWY-5005 | biotin biosynthesis II |
| PWY-5747 | 2-methylcitrate cycle II |
| PWY-6151 | S-adenosyl-L-methionine cycle I |
| *PWY-6339* | |
| PWY-6121 | 5-aminoimidazole ribonucleotide biosynthe |
| FAO-PWY | fatty acid β-oxidation I (generic) |
| COLANSYN-PWY | colanic acid building blocks biosynthesis |
| PWY-6122 | 5-aminoimidazole ribonucleotide biosynthe |
| PWY-6277 | 5-aminoimidazole ribonucleotide biosynthe |
| AST-PWY | L-arginine degradation II (AST pathway) |
| PWY-5028 | L-histidine degradation II |
| *PWY-5941* | |
| *PWY-6572* | |
| P124-PWY | |
| PWY-5484 | glycolysis II (from fructose 6-phosphate) |
| PPGPPMET-PWY | ppGpp metabolism |
| PWY-7295 | L-arabinose degradation IV |
| FASYN-INITIAL-PWY | fatty acid biosynthesis initiation |
| GLUCOSE1PMETAB-PWY | glucose and glucose-1-phosphate degrada |
| *NONMEVIPP-PWY* | |

| | |
|---|---|
| *PWY-7560* | |
| P281-PWY | 3-phenylpropanoate degradation |
| ASPASN-PWY | L-aspartate and L-asparagine biosynthesis |
| PWY-5430 | meta cleavage pathway of aromatic compo |
| *PWY-3801* | |
| PWY-6737 | starch degradation V |
| *DENITRIFICATION-PWY* | |
| PWY-5384 | sucrose degradation IV (sucrose phosphor |
| GALLATE-DEGRADATION-I-PWY | gallate degradation II |
| PWY-5154 | L-arginine biosynthesis III (via N-acetyl-L-c |
| PWY-7323 | GDP-mannose-derived O-antigen building |
| METHYLGALLATE-DEGRADATION-PWY | methylgallate degradation |
| GLYCOLYSIS-TCA-GLYOX-BYPASS | glycolysis, pyruvate dehydrogenase, TCA, |
| *PWY-5177* | |
| *P162-PWY* | |
| THRESYN-PWY | L-threonine biosynthesis |
| GLYCOCAT-PWY | glycogen degradation I |
| CRNFORCAT-PWY | creatinine degradation I |
| PWY-5104 | L-isoleucine biosynthesis IV |
| PWY-6628 | L-phenylalanine biosynthesis |
| *PWY-5304* | |
| KETOGLUCONMET-PWY | ketogluconate metabolism |
| ANAEROFRUCAT-PWY | homolactic fermentation |
| PWY-6630 | L-tyrosine biosynthesis |
| *PWY-7374* | |
| PWY-621 | sucrose degradation III (sucrose invertase) |
| PWY-6467 | Kdo transfer to lipid IVA III (Chlamydia) |
| PWY-5181 | toluene degradation III (aerobic) (via p-cres |
| *P621-PWY* | |
| 1CMET2-PWY | folate transformations III (E. coli) |
| TYRFUMCAT-PWY | L-tyrosine degradation I |
| PWYG-321 | mycolate biosynthesis |
| *PWY-6470* | |

| eJTK_BF | Peak | AvProp | Superclass | Subclass |
|---|---|---|---|---|
| 1.9769E-07 | 21.54656 | 0.0048215 | Biosynthesis | Tetrapyrrole Biosynthesis |
| 6.08E-07 | 21.39857 | 0.00475505 | Biosynthesis | Cofactor, Carrier, and Vitamin Biosynthesis |
| 1.6956E-06 | 20.78448 | 0.00474703 | Biosynthesis | Cofactor, Carrier, and Vitamin Biosynthesis |
| 3.233E-06 | 21.16595 | 0.00491236 | Biosynthesis | Tetrapyrrole Biosynthesis |
| 3.6299E-06 | 20.70408 | 0.00538561 | Biosynthesis | Nucleoside and Nucleotide Biosynthesis |
| 3.8455E-06 | 20.48015 | 0.00467968 | Biosynthesis | Nucleoside and Nucleotide Biosynthesis |
| 5.1229E-06 | 20.68864 | 0.00469437 | Biosynthesis | Nucleoside and Nucleotide Biosynthesis |
| 6.0763E-06 | 21.6987 | 0.00605258 | Biosynthesis | Amino Acid Biosynthesis |
| 6.4304E-06 | 20.47147 | 0.00465169 | Biosynthesis | Polyprenyl Biosynthesis |
| 7.6484E-06 | 4.48257 | 1.40E-05 | Biosynthesis | Amino Acid Biosynthesis |
| 7.6484E-06 | 4.233041 | 1.60E-05 | Biosynthesis | Amino Acid Biosynthesis |
| 1.065E-05 | 20.75974 | 0.00575682 | Biosynthesis | Nucleoside and Nucleotide Biosynthesis |
| 1.2574E-05 | 20.70333 | 0.00557962 | Biosynthesis | Nucleoside and Nucleotide Biosynthesis |
| 1.4037E-05 | 20.7344 | 0.00599543 | Biosynthesis | Nucleoside and Nucleotide Biosynthesis |
| 1.6544E-05 | 20.41131 | 0.0051802 | Biosynthesis | Amino Acid Biosynthesis |
| 2.1702E-05 | 20.3798 | 0.00581327 | Biosynthesis | Nucleoside and Nucleotide Biosynthesis |
| 3.6354E-05 | 0.03907041 | 6.67E-06 | Biosynthesis | Cofactor, Carrier, and Vitamin Biosynthesis |
| 3.9023E-05 | 20.48819 | 0.00555099 | Biosynthesis | Nucleoside and Nucleotide Biosynthesis |
| 5.6235E-05 | 23.84892 | 0.00359766 | Macromolecul | Nucleic Acid Processing |
| 0.0001264 | 22.28346 | 0.00632471 | Biosynthesis | Amino Acid Biosynthesis |
| 0.00018886 | 20.73949 | 0.0047148 | Biosynthesis | Cofactor, Carrier, and Vitamin Biosynthesis |
| 0.00021681 | 20.73949 | 0.00683059 | Biosynthesis | Amino Acid Biosynthesis |
| 0.00021681 | 22.63277 | 0.00683059 | Biosynthesis | Amino Acid Biosynthesis |
| 0.00022755 | 23.14395 | 0.00742455 | Biosynthesis | Amino Acid Biosynthesis |
| 0.00024244 | 0.5374792 | 6.62E-06 | Biosynthesis | Secondary Metabolite Biosynthesis |
| 0.00025057 | 22.9034161 | 0.01302978 | Generation of | Respiration |
| 0.00032162 | 20.18175 | 0.00525593 | Biosynthesis | Cofactor, Carrier, and Vitamin Biosynthesis |
| 0.00037373 | 19.66225 | 0.00015046 | Degradation/U | Aromatic Compound Degradation |
| 0.00053273 | 11.2850558 | 0.00330237 | Biosynthesis | Cofactor, Carrier, and Vitamin Biosynthesis |
| *0.00058149* | | *2.29E-05* | | |
| 0.00068025 | 21.45754 | 0.0053221 | Biosynthesis | Carbohydrate Biosynthesis |
| 0.00074546 | 20.61366 | 0.00457884 | Macromolecul | Nucleic Acid Processing |
| 0.00078023 | 22.65912 | 0.00498716 | Generation of | TCA cycle |
| 0.00081654 | 20.17477 | 0.00506404 | Biosynthesis | Aminoacyl-tRNA Charging |
| 0.00085443 | 20.14463 | 0.00480638 | Biosynthesis | Cell Structure Biosynthesis |
| 0.00085443 | 19.76157 | 0.00532468 | Biosynthesis | Amino Acid Biosynthesis |
| 0.00089397 | 21.73842 | 0.0056171 | Biosynthesis | Amino Acid Biosynthesis |
| 0.00093524 | 20.16514 | 0.00454139 | Biosynthesis | Cofactor, Carrier, and Vitamin Biosynthesis |
| 0.00093524 | 21.49091 | 0.00520434 | Biosynthesis | Nucleic Acid Processing |
| 0.00110765 | 19.61249 | 1.26E-05 | Degradation/U | C1 Compound Utilization and Assimilation |
| 0.00114947 | 22.69257 | 0.00557642 | Biosynthesis | Amino Acid Biosynthesis |
| 0.00116991 | 21.22301 | 0.00569836 | Biosynthesis | Nucleoside and Nucleotide Biosynthesis |
| 0.00116991 | 21.22301 | 0.00569836 | Biosynthesis | Nucleoside and Nucleotide Biosynthesis |
| 0.00133624 | 21.64033 | 0.00489245 | Biosynthesis | Nucleoside and Nucleotide Biosynthesis |
| 0.00152462 | 21.46389 | 0.00500977 | Biosynthesis | Amino Acid Biosynthesis |
| 0.00170401 | 22.45109 | 0.00541741 | Biosynthesis | Amino Acid Biosynthesis |
| 0.00189503 | 21.62173 | 0.00531178 | Biosynthesis | Nucleoside and Nucleotide Biosynthesis |
| 0.00192183 | 19.93268 | 0.00484681 | Biosynthesis | Cell Structure Biosynthesis |
| *0.00225795* | | *5.74E-05* | *Biosynthesis* | *Tetrapyrrole Biosynthesis* |
| *0.00225795* | | | | |
| 0.0024496 | 22.63423 | 1.18E-05 | Biosynthesis | Cofactor, Carrier, and Vitamin Biosynthesis |
| 0.00267953 | 20.07163 | 0.00598838 | Biosynthesis | Nucleoside and Nucleotide Biosynthesis |

| | | | | |
|---|---|---|---|---|
| 0.00321711 | 23.43107 | 0.00498248 | Generation of | TCA cycle |
| 0.00329081 | 21.63897 | 0.00516146 | Generation of | Photosynthesis |
| 0.00349478 | 12.0174 | 0.00392304 | Degradation/L | Amine and Polyamine Degradation |
| 0.00349478 | 11.20282 | 0.00260981 | Biosynthesis | Carbohydrate Biosynthesis |
| 0.00370984 | 19.0735 | 0.00524028 | Biosynthesis | Amino Acid Biosynthesis |
| 0.00379469 | 23.45178 | 0.00494792 | Generation of | TCA cycle |
| 0.00418368 | 20.29138 | 0.00467053 | Biosynthesis | Cell Structure Biosynthesis |
| 0.00428981 | 0.65574879 | 0.00422768 | Generation of | Glycolysis |
| 0.00446779 | 11.4186 | 0.00223592 | Detoxification | Antibiotic Resistance |
| 0.0047134 | 20.21351 | 0.00469892 | Biosynthesis | Cell Structure Biosynthesis |
| *0.00476722* | | *0.00473299* | *Biosynthesis* | *Carbohydrate Biosynthesis* |
| 0.00525071 | 0.34125991 | 0.00202591 | Biosynthesis | Amino Acid Biosynthesis |
| 0.00546545 | 11.2080329 | 0.0045965 | Degradation/L | Secondary Metabolite Degradation |
| *0.00548263* | | | | |
| 0.00624416 | | | | |
| 0.00657558 | | | | |
| 0.00659878 | | | | |
| 0.00666687 | 11.3398854 | 0.00255882 | Degradation/L | Aromatic Compound Degradation |
| 0.00666687 | 1.677602 | 0.00838107 | Biosynthesis | Fatty Acid and Lipid Biosynthesis |
| 0.00693479 | 10.90911 | 0.00296217 | Biosynthesis | Amine and Polyamine Biosynthesis |
| *0.00771064* | | | | |
| 0.00880855 | 23.89241 | 0.00021728 | Degradation/L | Aromatic Compound Degradation |
| 0.0093592 | 19.78353 | 0.00488439 | Biosynthesis | Cofactor, Carrier, and Vitamin Biosynthesis |
| 0.00946556 | 11.2704824 | 0.00273328 | Degradation/L | Secondary Metabolite Degradation |
| 0.00946556 | 11.1341382 | 0.00252207 | Degradation/L | Aromatic Compound Degradation |
| 0.00946556 | 11.1341272 | 0.00252207 | Degradation/L | Aromatic Compound Degradation |
| 0.01017709 | 19.28465 | 0.00552088 | Biosynthesis | Nucleoside and Nucleotide Biosynthesis |
| 0.01021955 | 10.90585 | 0.00491186 | Generation of | Pentose Phosphate Pathways |
| 0.01021955 | 11.2089401 | 0.00288115 | Degradation/L | Inorganic Nutrient Metabolism |
| 0.01061698 | 11.4923 | 0.00245675 | Degradation/L | Amine and Polyamine Degradation |
| 0.01096514 | 19.41834 | 0.00460367 | Biosynthesis | Nucleoside and Nucleotide Biosynthesis |
| 0.0114549 | 11.3869723 | 0.00279599 | Biosynthesis | Cofactor, Carrier, and Vitamin Biosynthesis |
| 0.01189632 | 10.9019595 | 0.00577375 | Degradation/L | Aromatic Compound Degradation |
| 0.01235336 | 10.3625814 | 0.00425075 | Degradation/L | Amino Acid Degradation |
| 0.01280344 | 2.08838 | 1.36E-05 | Biosynthesis | Cofactor, Carrier, and Vitamin Biosynthesis |
| 0.01282651 | 13.56752 | 0.00373373 | Degradation/L | Carboxylate Degradation |
| 0.01282651 | 0.42860774 | 0.00247923 | Biosynthesis | Amino Acid Biosynthesis |
| *0.01285804* | | | | |
| 0.01367906 | 18.82623 | 0.00562554 | Biosynthesis | Nucleoside and Nucleotide Biosynthesis |
| 0.01393654 | 1.313601 | 0.00807021 | Degradation/L | Fatty Acid and Lipid Degradation |
| 0.01422752 | 10.8309071 | 0.00469205 | Biosynthesis | Carbohydrate Biosynthesis |
| 0.01525607 | 19.0118 | 0.00517144 | Biosynthesis | Nucleoside and Nucleotide Biosynthesis |
| 0.01525607 | 19.01174 | 0.00517144 | Biosynthesis | Nucleoside and Nucleotide Biosynthesis |
| 0.01603309 | 11.77824 | 0.00301227 | Degradation/L | Amino Acid Degradation |
| 0.01663401 | 10.4314454 | 0.00370562 | Degradation/L | Amino Acid Degradation |
| *0.01684358* | | | | |
| *0.01746672* | | | | |
| 0.01777494 | | | | |
| 0.01789822 | 0.4452275 | 0.0036826 | Generation of | Glycolysis |
| 0.0182592 | 14.18736 | 0.00412128 | Biosynthesis | Metabolic Regulator Biosynthesis |
| 0.01848593 | 23.07375 | 0.0001518 | Degradation/L | Carbohydrate Degradation |
| 0.01924983 | 11.86353 | 0.00805644 | Biosynthesis | Fatty Acid and Lipid Biosynthesis |
| 0.02069418 | 10.8865373 | 0.00465811 | Degradation/L | Carbohydrate Degradation |
| *0.02104114* | | | | |

| | | | | |
|---|---|---|---|---|
| 0.02104114 | | | | |
| 0.02130361 | 2.843223 | 0.0008236 | Degradation/L | Aromatic Compound Degradation |
| 0.02145288 | 10.55693 | 0.00313983 | Biosynthesis | Amino Acid Biosynthesis |
| 0.02190184 | 20.76845 | 0.00013149 | Degradation/L | Aromatic Compound Degradation |
| *0.02267801* | | | | |
| 0.02304697 | 10.6860237 | 0.00415224 | Degradation/L | Carbohydrate Degradation |
| *0.02376505* | | | | |
| 0.02388387 | 10.5101434 | 0.00398253 | Degradation/L | Carbohydrate Degradation |
| 0.02469984 | 0.1431483 | 0.00026736 | Degradation/L | Aromatic Compound Degradation |
| 0.02474838 | 12.67755 | 0.00530625 | Biosynthesis | Amino Acid Biosynthesis |
| 0.02552565 | 10.4768417 | 0.00449515 | Biosynthesis | Carbohydrate Biosynthesis |
| 0.02559135 | 0.184094 | 0.00031995 | Degradation/L | Aromatic Compound Degradation |
| 0.02751568 | 0.2374286 | 0.00457682 | Generation of | Glycolysis |
| *0.02866037* | | | | |
| 0.02929334 | | | | |
| 0.0294077 | 21.52065 | 0.00513648 | Biosynthesis | Amino Acid Biosynthesis |
| 0.03275885 | 10.75643 | 0.00383843 | Degradation/L | Carbohydrate Degradation |
| 0.03391035 | 14.33289 | 0.00031199 | Degradation/L | Amine and Polyamine Degradation |
| 0.03391035 | 1.33327013 | 0.00242679 | Biosynthesis | Amino Acid Biosynthesis |
| 0.0351669 | 12.35229 | 0.00612333 | Biosynthesis | Amino Acid Biosynthesis |
| 0.03628591 | | | | |
| 0.03694973 | 4.356073 | 0.00103866 | Degradation/L | Carboxylate Degradation |
| 0.04023668 | 1.42100487 | 0.00338385 | Generation of | Fermentation |
| 0.04023668 | 0.6938823 | 0.0048852 | Biosynthesis | Amino Acid Biosynthesis |
| 0.04105068 | | | | |
| 0.04162311 | 10.2521586 | 0.00383702 | Degradation/L | Carbohydrate Degradation |
| 0.04300116 | 19.25475 | 0.00439794 | Biosynthesis | Cell Structure Biosynthesis |
| 0.04448228 | 8.66485 | 0.00300959 | Degradation/L | Aromatic Compound Degradation |
| 0.0456403 | | | | |
| 0.04604482 | 12.2390022 | 0.00614593 | Biosynthesis | Cofactor, Carrier, and Vitamin Biosynthesis |
| 0.04758322 | 10.41203 | 0.00457516 | Degradation/L | Amino Acid Degradation |
| 0.04915475 | 13.06886 | 0.00834762 | Biosynthesis | Fatty Acid and Lipid Biosynthesis |
| 0.04921667 | | | | |

| ID | Desc | eJTK_BF | Peak |
|---|---|---|---|
| POLYAMINSYN3-PWY | polyamine biosynthesis II | 0.01181769 | 14.24225 |
| PWY-2942 | L-lysine biosynthesis III | 0.00107423 | 0.4640838 |
| PWY-6151 | S-adenosyl-L-methionine cycle I | 0.00291762 | 2.50907 |
| PWY-4361 | S-methyl-5-thio-α-D-ribose 1-pho | 0.00521667 | 1.359399 |
| PWY-7527 | L-methionine salvage cycle III | 0.00521667 | 1.368213 |
| PWY-5505 | L-glutamate and L-glutamine bio | 0.00967176 | 2.25966707 |
| BRANCHED-CHAIN-AA-SYN-PWY | branched chain amino acid biosy | 0.01139516 | 23.77327 |
| ASPASN-PWY | L-aspartate and L-asparagine bio | 0.01319197 | 14.90918 |
| ILEUSYN-PWY | L-isoleucine biosynthesis I (from | 0.01701832 | 0.2098248 |
| VALSYN-PWY | L-valine biosynthesis | 0.01701832 | 0.2098239 |
| PWY-3001 | L-isoleucine biosynthesis I | 0.0174237 | 22.11811 |
| PWY-5101 | L-isoleucine biosynthesis II | 0.01857189 | 0.6445518 |
| PWY-6630 | L-tyrosine biosynthesis | 0.02842071 | 2.593421 |
| PWY-5097 | L-lysine biosynthesis VI | 0.02930627 | 23.14649 |
| GLUTORN-PWY | L-ornithine biosynthesis I | 0.03610503 | 14.6357 |
| PWY-5154 | L-arginine biosynthesis III (via N- | 0.04233608 | 14.9656 |
| PWY-6165 | chorismate biosynthesis II (archa | 0.00010365 | 1.616014 |
| PWY0-1241 | ADP-L-glycero-β-D-manno-hepto | 0.02964945 | 14.7731551 |
| PWY-6749 | CMP-legionaminate biosynthesis | 0.03013338 | 0.2248954 |
| PWY-5659 | GDP-mannose biosynthesis | 0.03610503 | 15.2239192 |
| PWY-6167 | flavin biosynthesis II (archaea) | 6.5205E-05 | 1.643688 |
| PWY-6654 | phosphopantothenate biosynthes | 0.00010365 | 1.637631 |
| PWY0-845 | pyridoxal 5'-phosphate biosynthe | 0.00141469 | 14.1404798 |
| P261-PWY | coenzyme M biosynthesis I | 0.007646 | 23.25483 |
| BIOTIN-BIOSYNTHESIS-PWY | biotin biosynthesis I | 0.01011498 | 14.26564 |
| 1CMET2-PWY | folate transformations III (E. coli) | 0.02301099 | 15.01738 |
| PWY-5198 | factor 420 biosynthesis II (mycob | 0.02949096 | 2.445492 |
| PWY-5920 | heme b biosynthesis from glycine | 0.02964945 | 1.46966324 |
| PWY-7376 | cob(II)yrinate a,c-diamide biosyn | 0.03071576 | 14.9517 |
| PYRIDOXSYN-PWY | pyridoxal 5'-phosphate biosynthe | 0.03071576 | 14.20977 |
| FOLSYN-PWY | tetrahydrofolate biosynthesis and | 0.04069345 | 15.34843 |
| PWY-7663 | gondoate biosynthesis (anaerobi | 0.00439425 | 14.68228 |
| PWYG-321 | mycolate biosynthesis | 0.00587285 | 14.54942 |
| PWY-7664 | oleate biosynthesis IV (anaerobic | 0.01129236 | 14.75275 |
| FASYN-ELONG-PWY | fatty acid elongation -- saturated | 0.01181769 | 14.63671 |
| PWY0-862 | (5Z)-dodecenoate biosynthesis I | 0.01353166 | 15.04154 |
| FASYN-INITIAL-PWY | fatty acid biosynthesis initiation (I | 0.01927897 | 14.4426076 |
| PWY-6282 | palmitoleate biosynthesis I (from | 0.0249699 | 14.64343 |
| PPGPPMET-PWY | ppGpp metabolism | 0.00703722 | 14.32313 |
| PWY-7221 | guanosine ribonucleotides de no | 0.00322773 | 23.1981 |
| PWY-6125 | guanosine nucleotides de novo b | 0.00333618 | 22.47322 |
| PWY-7228 | guanosine nucleotides de novo b | 0.00393722 | 22.85595 |
| PWY-7200 | pyrimidine deoxyribonucleoside s | 0.01439528 | 2.388056 |
| PWY-7220 | adenosine deoxyribonucleotides | 0.01903241 | 20.60992 |
| PWY-7222 | guanosine deoxyribonucleotides | 0.01903241 | 20.60982 |
| PWY-7199 | pyrimidine deoxyribonucleosides | 0.02292211 | 2.079752 |
| PWY-841 | purine nucleotides de novo biosy | 0.02470772 | 21.56337 |
| PWY-6609 | adenine and adenosine salvage | 0.03610503 | 2.16887792 |
| PWY-6122 | 5-aminoimidazole ribonucleotide | 0.04623035 | 20.8905 |
| PWY-6277 | 5-aminoimidazole ribonucleotide | 0.04623035 | 20.89054 |
| PWY-6519 | 8-amino-7-oxononanoate biosyn | 0.04762899 | 14.0817419 |
| POLYISOPRENSYN-PWY | polyisoprenoid biosynthesis (E. c | 0.01524305 | 19.21801 |

| | | | |
|---|---|---|---|
| PWY-6581 | spirilloxanthin and 2,2'-diketo-spi | 0.00198822 | 23.27743 |
| PWY-6174 | mevalonate pathway II (haloarch | 0.00479574 | 23.62934 |
| ENTBACSYN-PWY | enterobactin biosynthesis | 0.04486645 | 2.531403 |
| CRNFORCAT-PWY | creatinine degradation I | 0.00299958 | 14.82222 |
| PWY-5022 | 4-aminobutanoate degradation V | 0.00703722 | 14.20711 |
| PWY-7431 | aromatic biogenic amine degrada | 0.03198989 | 14.72142 |
| PWY-6505 | L-tryptophan degradation XII (Ge | 0.0101952 | 2.545333 |
| AST-PWY | L-arginine degradation II (AST pa | 0.02103614 | 14.3173977 |
| PWY-5655 | L-tryptophan degradation IX | 0.03604063 | 2.545476 |
| PWY-6210 | 2-aminophenol degradation | 0.01037128 | 2.455527 |
| CATECHOL-ORTHO-CLEAVAGE-PWY | catechol degradation to β-ketoac | 0.01364088 | 14.15661 |
| PWY-5431 | aromatic compounds degradatio | 0.02835517 | 14.27361 |
| PWY-5430 | meta cleavage pathway of aroma | 0.02909483 | 3.131856 |
| PWY-5417 | catechol degradation III (ortho-cl | 0.02835517 | 14.273605 |
| PWY-5647 | 2-nitrobenzoate degradation I | 0.03604063 | 2.28502 |
| PWY-1882 | C1 compounds oxidation to CO2 | 0.03461024 | 21.64286 |
| DHGLUCONATE-PYR-CAT-PWY | glucose degradation (oxidative) | 0.00246111 | 12.14957 |
| PWY-5747 | 2-methylcitrate cycle II | 0.02716486 | 13.92641 |
| PWY-5654 | 2-amino-3-carboxymuconate ser | 0.03604063 | 2.199813 |
| PWY-6641 | sulfolactate degradation | 0.00251725 | 1.530559 |
| SO4ASSIM-PWY | assimilatory sulfate reduction I | 0.02301099 | 14.2719838 |
| PWY0-1533 | methylphosphonate degradation | 0.02964945 | 14.54508 |
| PWY0-1297 | purine deoxyribonucleosides deg | 0.01785758 | 2.266681 |
| P164-PWY | purine nucleobases degradation | 0.03403874 | 0.00405378 |
| PWY-6353 | purine nucleotides degradation II | 0.03757968 | 2.27563982 |
| PWY0-1296 | purine ribonucleosides degradati | 0.02206736 | 2.125236 |
| P221-PWY | octane oxidation | 0.00807222 | 14.3452877 |
| PWY0-1261 | anhydromuropeptides recycling I | 0.02013786 | 2.90950758 |
| PWY-7237 | myo-, chiro- and scyllo-inositol de | 0.04069345 | 14.8459775 |
| PWY0-1338 | polymyxin resistance | 0.00844739 | 14.28478 |
| ANAEROFRUCAT-PWY | homolactic fermentation | 0.0210286 | 2.898976 |
| GLYCOLYSIS | glycolysis I (from glucose 6-phos | 0.02395495 | 2.693878 |
| PWY-5484 | glycolysis II (from fructose 6-pho | 0.0380901 | 2.759216 |
| PENTOSE-P-PWY | pentose phosphate pathway | 0.00462354 | 15.03732 |
| PWY-181 | photorespiration | 0.02964945 | 3.082635 |
| CALVIN-PWY | Calvin-Benson-Bassham cycle | 0.03225926 | 22.72633 |
| PWY-3781 | aerobic respiration I (cytochrome | 0.02508665 | 2.69166991 |
| P105-PWY | TCA cycle IV (2-oxoglutarate dec | 0.00528626 | 2.622187 |
| PWY-6969 | TCA cycle V (2-oxoglutarate:ferr | 0.00554865 | 2.535885 |
| PWY-7254 | TCA cycle VII (acetate-producers | 0.00906749 | 13.1901661 |
| REDCITCYC | TCA cycle VI (Helicobacter) | 0.00967176 | 13.43173 |
| TCA-GLYOX-BYPASS | glyoxylate bypass and TCA | 0.03198989 | 3.504293 |

| AvProp | Superclass | Subclass |
|---|---|---|
| 0.00298841 | Biosynthesis | Amine and Polyamine Biosynthesis |
| 0.00550471 | Biosynthesis | Amino Acid Biosynthesis |
| 0.00281659 | Biosynthesis | Amino Acid Biosynthesis |
| 1.0959E-06 | Biosynthesis | Amino Acid Biosynthesis |
| 1.8009E-06 | Biosynthesis | Amino Acid Biosynthesis |
| 0.00195807 | Biosynthesis | Amino Acid Biosynthesis |
| 0.00626044 | Biosynthesis | Amino Acid Biosynthesis |
| 0.00319298 | Biosynthesis | Amino Acid Biosynthesis |
| 0.00683603 | Biosynthesis | Amino Acid Biosynthesis |
| 0.00683603 | Biosynthesis | Amino Acid Biosynthesis |
| 0.00549441 | Biosynthesis | Amino Acid Biosynthesis |
| 0.0074879 | Biosynthesis | Amino Acid Biosynthesis |
| 0.0050413 | Biosynthesis | Amino Acid Biosynthesis |
| 0.00531615 | Biosynthesis | Amino Acid Biosynthesis |
| 0.00601301 | Biosynthesis | Amino Acid Biosynthesis |
| 0.00520577 | Biosynthesis | Amino Acid Biosynthesis |
| 5.089E-06 | Biosynthesis | Aromatic Compound Biosynthesis |
| 0.00260498 | Biosynthesis | Carbohydrate Biosynthesis |
| 2.3537E-05 | Biosynthesis | Carbohydrate Biosynthesis |
| 0.00587121 | Biosynthesis | Carbohydrate Biosynthesis |
| 6.3302E-06 | Biosynthesis | Cofactor, Carrier, and Vitamin Biosynthesis |
| 3.8116E-06 | Biosynthesis | Cofactor, Carrier, and Vitamin Biosynthesis |
| 0.00343884 | Biosynthesis | Cofactor, Carrier, and Vitamin Biosynthesis |
| 2.6094E-06 | Biosynthesis | Cofactor, Carrier, and Vitamin Biosynthesis |
| 0.00508217 | Biosynthesis | Cofactor, Carrier, and Vitamin Biosynthesis |
| 0.00597742 | Biosynthesis | Cofactor, Carrier, and Vitamin Biosynthesis |
| 9.1684E-06 | Biosynthesis | Cofactor, Carrier, and Vitamin Biosynthesis |
| 0.001248 | Biosynthesis | Cofactor, Carrier, and Vitamin Biosynthesis |
| 0.00289434 | Biosynthesis | Cofactor, Carrier, and Vitamin Biosynthesis |
| 0.00387373 | Biosynthesis | Cofactor, Carrier, and Vitamin Biosynthesis |
| 0.00555335 | Biosynthesis | Cofactor, Carrier, and Vitamin Biosynthesis |
| 0.00849158 | Biosynthesis | Fatty Acid and Lipid Biosynthesis |
| 0.00810194 | Biosynthesis | Fatty Acid and Lipid Biosynthesis |
| 0.00730762 | Biosynthesis | Fatty Acid and Lipid Biosynthesis |
| 0.00760872 | Biosynthesis | Fatty Acid and Lipid Biosynthesis |
| 0.00694986 | Biosynthesis | Fatty Acid and Lipid Biosynthesis |
| 0.00799134 | Biosynthesis | Fatty Acid and Lipid Biosynthesis |
| 0.00716247 | Biosynthesis | Fatty Acid and Lipid Biosynthesis |
| 0.00386578 | Biosynthesis | Metabolic Regulator Biosynthesis |
| 0.00483871 | Biosynthesis | Nucleoside and Nucleotide Biosynthesis |
| 0.00509612 | Biosynthesis | Nucleoside and Nucleotide Biosynthesis |
| 0.00521677 | Biosynthesis | Nucleoside and Nucleotide Biosynthesis |
| 0.00235527 | Biosynthesis | Nucleoside and Nucleotide Biosynthesis |
| 0.00551023 | Biosynthesis | Nucleoside and Nucleotide Biosynthesis |
| 0.00551023 | Biosynthesis | Nucleoside and Nucleotide Biosynthesis |
| 0.00147325 | Biosynthesis | Nucleoside and Nucleotide Biosynthesis |
| 0.0052573 | Biosynthesis | Nucleoside and Nucleotide Biosynthesis |
| 0.00136183 | Biosynthesis | Nucleoside and Nucleotide Biosynthesis |
| 0.00510372 | Biosynthesis | Nucleoside and Nucleotide Biosynthesis |
| 0.00510372 | Biosynthesis | Nucleoside and Nucleotide Biosynthesis |
| 0.00590119 | Biosynthesis | Other |
| 0.00448635 | Biosynthesis | Polyprenyl Biosynthesis |

| | | |
|---|---|---|
| 2.7928E-06 | Biosynthesis | Secondary Metabolite Biosynthesis |
| 2.1503E-06 | Biosynthesis | Secondary Metabolite Biosynthesis |
| 0.00117934 | Biosynthesis | Secondary Metabolite Biosynthesis |
| 0.0003129 | Degradation/U | Amine and Polyamine Degradation |
| 0.00380292 | Degradation/U | Amine and Polyamine Degradation |
| 0.00229814 | Degradation/U | Amine and Polyamine Degradation |
| 2.7211E-05 | Degradation/U | Amino Acid Degradation |
| 0.0028984 | Degradation/U | Amino Acid Degradation |
| 6.5895E-05 | Degradation/U | Amino Acid Degradation |
| 1.1082E-05 | Degradation/U | Aromatic Compound Degradation |
| 0.00269879 | Degradation/U | Aromatic Compound Degradation |
| 0.00269496 | Degradation/U | Aromatic Compound Degradation |
| 9.4917E-05 | Degradation/U | Aromatic Compound Degradation |
| 0.00269496 | Degradation/U | Aromatic Compound Degradation |
| 3.4473E-05 | Degradation/U | Aromatic Compound Degradation |
| 2.2415E-06 | Degradation/U | C1 Compound Utilization and Assimilation |
| 0.00072437 | Degradation/U | Carbohydrate Degradation |
| 0.00351219 | Degradation/U | Carboxylate Degradation |
| 2.9498E-05 | Degradation/U | Carboxylate Degradation |
| 3.7385E-06 | Degradation/U | Inorganic Nutrient Metabolism |
| 0.00698497 | Degradation/U | Inorganic Nutrient Metabolism |
| 0.00294204 | Degradation/U | Inorganic Nutrient Metabolism |
| 0.00106358 | Degradation/U | Nucleoside and Nucleotide Degradation |
| 0.0002128 | Degradation/U | Nucleoside and Nucleotide Degradation |
| 0.00222637 | Degradation/U | Nucleoside and Nucleotide Degradation |
| 0.00095736 | Degradation/U | Nucleoside and Nucleotide Degradation |
| 0.00351393 | Degradation/U | Other |
| 0.00437805 | Degradation/U | Secondary Metabolite Degradation |
| 0.00493102 | Degradation/U | Secondary Metabolite Degradation |
| 0.002264 | Detoxification | Antibiotic Resistance |
| 0.00371325 | Generation of | Fermentation |
| 0.00430438 | Generation of | Glycolysis |
| 0.00386352 | Generation of | Glycolysis |
| 0.00509678 | Generation of | Pentose Phosphate Pathways |
| 0.00260144 | Generation of | Photosynthesis |
| 0.0050757 | Generation of | Photosynthesis |
| 0.01279567 | Generation of | Respiration |
| 0.004922 | Generation of | TCA cycle |
| 0.00495477 | Generation of | TCA cycle |
| 0.00610661 | Generation of | TCA cycle |
| 0.00637222 | Generation of | TCA cycle |
| 0.00592074 | Generation of | TCA cycle |

| ID | Desc | eJTK_BF | Peak |
|---|---|---|---|
| PWY-5505 | L-glutamate and L-g | 0.001421708 | 3.970232131 |
| ILEUSYN-PWY | L-isoleucine biosynth | 0.004543789 | 4.345035 |
| VALSYN-PWY | L-valine biosynthesis | 0.004543789 | 4.345035 |
| HSERMETANA-PWY | L-methionine biosyn | 0.005581124 | 3.037407 |
| BRANCHED-CHAIN-AA-SYN-PWY | branched chain amir | 0.006054548 | 2.962367 |
| PWY-3001 | L-isoleucine biosynth | 0.006143178 | 1.443449 |
| PWY-5101 | L-isoleucine biosynth | 0.009394609 | 4.222059 |
| THRESYN-PWY | L-threonine biosynth | 0.009644006 | 0.5940647 |
| PWY-5097 | L-lysine biosynthesis | 0.026371586 | 0.9127464 |
| PWY-5104 | L-isoleucine biosynth | 0.034570952 | 3.902741 |
| HISTSYN-PWY | L-histidine biosynthe | 0.043492245 | 22.16943 |
| ARGSYNBSUB-PWY | L-arginine biosynthe | 0.045053243 | 18.09423 |
| DAPLYSINESYN-PWY | L-lysine biosynthesis | 0.048327538 | 19.53305 |
| PWY-6165 | chorismate biosynth | 0.002854316 | 1.615874 |
| PWY0-1241 | ADP-L-glycero-β-D-r | 0.004359157 | 15.23928214 |
| UDPNAGSYN-PWY | UDP-N-acetyl-D-gluc | 0.022574372 | 20.28038 |
| OANTIGEN-PWY | O-antigen building bl | 0.031490649 | 19.55511 |
| GLYCOGENSYNTH-PWY | glycogen biosynthes | 0.001774644 | 14.70316207 |
| PWY0-845 | pyridoxal 5'-phospha | 0.00274041 | 15.0958907 |
| PWY-6167 | flavin biosynthesis II | 0.003190803 | 0.8689589 |
| NADSYN-PWY | NAD de novo biosyn | 0.005509694 | 5.05939 |
| PYRIDNUCSAL-PWY | NAD salvage pathwa | 0.008020816 | 17.18851 |
| PWY-6654 | phosphopantothenat | 0.009416284 | 0.9057542 |
| PYRIDOXSYN-PWY | pyridoxal 5'-phospha | 0.012815443 | 16.00489 |
| PYRIDNUCSYN-PWY | NAD de novo biosyn | 0.02345085 | 18.96047 |
| P381-PWY | adenosylcobalamin b | 0.027474896 | 4.741932 |
| NAD-BIOSYNTHESIS-II | NAD salvage pathwa | 0.030275161 | 3.154132 |
| PWY-7539 | 6-hydroxymethyl-dihy | 0.037578797 | 17.36494 |
| UBISYN-PWY | ubiquinol-8 biosynthe | 0.046664541 | 19.28877 |
| PWY-5973 | cis-vaccenate biosyn | 0.012255232 | 3.30482 |
| PWY-5989 | stearate biosynthesis | 0.027365974 | 9.254426 |
| PPGPPMET-PWY | ppGpp metabolism | 0.001954237 | 17.83819 |
| PWY-7197 | pyrimidine deoxyribo | 0.00351924 | 20.47036 |
| PWY-7211 | pyrimidine deoxyribo | 0.046664541 | 21.14813 |
| PWY-6519 | 8-amino-7-oxononar | 0.0194696 | 17.38199359 |
| PWY-7392 | taxadiene biosynthes | 0.016115597 | 5.105335729 |
| NONMEVIPP-PWY | methylerythritol phos | 0.020936278 | 20.3298 |
| PWY-7560 | methylerythritol phos | 0.020936278 | 20.3298 |
| PWY-5188 | tetrapyrrole biosynth | 0.01231821 | 0.5289638 |
| PWY-5189 | tetrapyrrole biosynth | 0.01616326 | 0.7363197 |
| PWY-5022 | 4-aminobutanoate de | 0.026949007 | 16.45844026 |
| PWY-5651 | L-tryptophan degrad | 0.00598376 | 5.136845 |
| PWY-5088 | L-glutamate degrada | 0.015852445 | 15.48333 |
| PWY-5655 | L-tryptophan degrad | 0.018159856 | 4.998904 |
| THREOCAT-PWY | L-threonine metaboli | 0.023239897 | 4.501625 |
| AST-PWY | L-arginine degradati | 0.020887282 | 16.51537 |
| GALLATE-DEGRADATION-I-PWY | gallate degradation I | 0.011866639 | 4.077029189 |
| METHYLGALLATE-DEGRADATION-PWY | methylgallate degrac | 0.012815443 | 4.012823 |
| PWY-5647 | 2-nitrobenzoate degr | 0.021102367 | 5.078541 |
| P184-PWY | protocatechuate deg | 0.02331383 | 3.670533 |
| P281-PWY | 3-phenylpropanoate | 0.025241389 | 6.233675 |
| PWY-5178 | toluene degradation | 0.037508947 | 4.922143 |

| | | | |
|---|---|---|---|
| PWY-5420 | catechol degradation | 0.04627134 | 5.325998 |
| GLYCOCAT-PWY | glycogen degradatio | 0.001554128 | 15.30074885 |
| PWY-6737 | starch degradation V | 0.003249712 | 15.27246744 |
| PWY-6901 | glucose and xylose c | 0.00528597 | 6.00861119 |
| PWY-621 | sucrose degradation | 0.014925558 | 14.84242644 |
| PWY-5384 | sucrose degradation | 0.015500719 | 16.05329 |
| PWY-6713 | L-rhamnose degrada | 0.018593581 | 14.69518 |
| GLUCOSE1PMETAB-PWY | glucose and glucose | 0.033374211 | 15.98753504 |
| PWY-6572 | chondroitin sulfate d | 0.049079073 | 4.743496 |
| PWY-5654 | 2-amino-3-carboxym | 0.022603345 | 5.083428 |
| PWY-4984 | urea cycle | 0.018058015 | 4.842164 |
| DENITRIFICATION-PWY | nitrate reduction I (d | 0.018234462 | 3.604276 |
| PWY490-3 | nitrate reduction VI ( | 0.039756278 | 3.985901 |
| PWY-6641 | sulfolactate degrada | 0.043606071 | 12.80359 |
| GALACT-GLUCUROCAT-PWY | hexuronide and hexu | 0.047899028 | 4.30078 |
| PWY-7237 | myo-, chiro- and scy | 0.039756278 | 15.01279896 |
| PWY0-1338 | polymyxin resistance | 0.022477812 | 15.03892 |
| ANAEROFRUCAT-PWY | homolactic fermenta | 0.005509694 | 4.164371 |
| P122-PWY | heterolactic ferment | 0.026182569 | 4.777284 |
| P108-PWY | pyruvate fermentatio | 0.045634158 | 4.426327 |
| PWY-5484 | glycolysis II (from fru | 0.019400183 | 3.671253 |
| GLYCOLYSIS-E-D | glycolysis and the En | 0.02115146 | 2.70668 |
| GLYCOLYSIS | glycolysis I (from glu | 0.02166923 | 3.290313 |
| ANAGLYCOLYSIS-PWY | glycolysis III (from gl | 0.034570952 | 4.755769 |
| PENTOSE-P-PWY | pentose phosphate p | 0.048857234 | 15.21065 |
| PWY-3781 | aerobic respiration I | 0.020131187 | 1.778433965 |
| GLYCOLYSIS-TCA-GLYOX-BYPASS | glycolysis, pyruvate d | 0.003620054 | 3.183469 |
| PWY-6969 | TCA cycle V (2-oxog | 0.031092676 | 2.883366 |
| P105-PWY | TCA cycle IV (2-oxog | 0.03708172 | 2.908569 |

| AvProp | Superclass | Subclass |
|---|---|---|
| 0.002246188 | Biosynthesis | Amino Acid Biosynthesis |
| 0.006881252 | Biosynthesis | Amino Acid Biosynthesis |
| 0.006881252 | Biosynthesis | Amino Acid Biosynthesis |
| 0.00510453 | Biosynthesis | Amino Acid Biosynthesis |
| 0.006341953 | Biosynthesis | Amino Acid Biosynthesis |
| 0.005689249 | Biosynthesis | Amino Acid Biosynthesis |
| 0.007470301 | Biosynthesis | Amino Acid Biosynthesis |
| 0.005253538 | Biosynthesis | Amino Acid Biosynthesis |
| 0.005496256 | Biosynthesis | Amino Acid Biosynthesis |
| 0.002913613 | Biosynthesis | Amino Acid Biosynthesis |
| 0.005140456 | Biosynthesis | Amino Acid Biosynthesis |
| 0.00530721 | Biosynthesis | Amino Acid Biosynthesis |
| 0.005604531 | Biosynthesis | Amino Acid Biosynthesis |
| 5.17277E-06 | Biosynthesis | Aromatic Compound Biosynthesis |
| 0.002440588 | Biosynthesis | Carbohydrate Biosynthesis |
| 0.00466699 | Biosynthesis | Carbohydrate Biosynthesis |
| 0.004881189 | Biosynthesis | Carbohydrate Biosynthesis |
| 0.003809261 | Biosynthesis | Carbohydrate Biosynthesis |
| 0.003157525 | Biosynthesis | Cofactor, Carrier, and Vitamin Biosynthesis |
| 7.00034E-06 | Biosynthesis | Cofactor, Carrier, and Vitamin Biosynthesis |
| 0.000921375 | Biosynthesis | Cofactor, Carrier, and Vitamin Biosynthesis |
| 0.004316655 | Biosynthesis | Cofactor, Carrier, and Vitamin Biosynthesis |
| 4.38788E-06 | Biosynthesis | Cofactor, Carrier, and Vitamin Biosynthesis |
| 0.003910828 | Biosynthesis | Cofactor, Carrier, and Vitamin Biosynthesis |
| 0.004549027 | Biosynthesis | Cofactor, Carrier, and Vitamin Biosynthesis |
| 0.000192164 | Biosynthesis | Cofactor, Carrier, and Vitamin Biosynthesis |
| 0.001814153 | Biosynthesis | Cofactor, Carrier, and Vitamin Biosynthesis |
| 0.005768482 | Biosynthesis | Cofactor, Carrier, and Vitamin Biosynthesis |
| 0.005099068 | Biosynthesis | Cofactor, Carrier, and Vitamin Biosynthesis |
| 0.00775419 | Biosynthesis | Fatty Acid and Lipid Biosynthesis |
| 0.007646438 | Biosynthesis | Fatty Acid and Lipid Biosynthesis |
| 0.003971792 | Biosynthesis | Metabolic Regulator Biosynthesis |
| 0.004539271 | Biosynthesis | Nucleoside and Nucleotide Biosynthesis |
| 0.005455668 | Biosynthesis | Nucleoside and Nucleotide Biosynthesis |
| 0.006187973 | Biosynthesis | Other |
| 0.001312291 | Biosynthesis | Secondary Metabolite Biosynthesis |
| 0.004416301 | Biosynthesis | Secondary Metabolite Biosynthesis |
| 0.004416301 | Biosynthesis | Secondary Metabolite Biosynthesis |
| 0.004890666 | Biosynthesis | Tetrapyrrole Biosynthesis |
| 0.004804664 | Biosynthesis | Tetrapyrrole Biosynthesis |
| 0.003690869 | Degradation/Utilization/Assimilation | Amine and Polyamine Degradation |
| 0.000641595 | Degradation/Utilization/Assimilation | Amino Acid Degradation |
| 3.75225E-06 | Degradation/Utilization/Assimilation | Amino Acid Degradation |
| 9.59624E-05 | Degradation/Utilization/Assimilation | Amino Acid Degradation |
| 7.16386E-05 | Degradation/Utilization/Assimilation | Amino Acid Degradation |
| 0.003072328 | Degradation/Utilization/Assimilation | Amino Acid Degradation |
| 0.000397 | Degradation/Utilization/Assimilation | Aromatic Compound Degradation |
| 0.000457403 | Degradation/Utilization/Assimilation | Aromatic Compound Degradation |
| 4.9579E-05 | Degradation/Utilization/Assimilation | Aromatic Compound Degradation |
| 0.00033991 | Degradation/Utilization/Assimilation | Aromatic Compound Degradation |
| 0.000882233 | Degradation/Utilization/Assimilation | Aromatic Compound Degradation |
| 0.000343784 | Degradation/Utilization/Assimilation | Aromatic Compound Degradation |

| | | |
|---|---|---|
| 0.000423899 | Degradation/Utilization/Assimilation | Aromatic Compound Degradation |
| 0.003742478 | Degradation/Utilization/Assimilation | Carbohydrate Degradation |
| 0.00402326 | Degradation/Utilization/Assimilation | Carbohydrate Degradation |
| 0.001292317 | Degradation/Utilization/Assimilation | Carbohydrate Degradation |
| 0.003793939 | Degradation/Utilization/Assimilation | Carbohydrate Degradation |
| 0.004027744 | Degradation/Utilization/Assimilation | Carbohydrate Degradation |
| 4.62587E-06 | Degradation/Utilization/Assimilation | Carbohydrate Degradation |
| 0.004558663 | Degradation/Utilization/Assimilation | Carbohydrate Degradation |
| 8.13244E-06 | Degradation/Utilization/Assimilation | Carbohydrate Degradation |
| 4.2839E-05 | Degradation/Utilization/Assimilation | Carboxylate Degradation |
| 0.001961605 | Degradation/Utilization/Assimilation | Inorganic Nutrient Metabolism |
| 0.000313593 | Degradation/Utilization/Assimilation | Inorganic Nutrient Metabolism |
| 0.000211471 | Degradation/Utilization/Assimilation | Inorganic Nutrient Metabolism |
| 1.27388E-05 | Degradation/Utilization/Assimilation | Inorganic Nutrient Metabolism |
| 0.000578658 | Degradation/Utilization/Assimilation | Other |
| 0.003953157 | Degradation/Utilization/Assimilation | Secondary Metabolite Degradation |
| 0.00193172 | Detoxification | Antibiotic Resistance |
| 0.003468229 | Generation of Precursor Metabolites | Fermentation |
| 0.00031409 | Generation of Precursor Metabolites | Fermentation |
| 0.001328978 | Generation of Precursor Metabolites | Fermentation |
| 0.004031083 | Generation of Precursor Metabolites | Glycolysis |
| 0.004387894 | Generation of Precursor Metabolites | Glycolysis |
| 0.004453307 | Generation of Precursor Metabolites | Glycolysis |
| 0.003902824 | Generation of Precursor Metabolites | Glycolysis |
| 0.004574708 | Generation of Precursor Metabolites | Pentose Phosphate Pathways |
| 0.013302668 | Generation of Precursor Metabolites | Respiration |
| 0.0045788 | Generation of Precursor Metabolites | TCA cycle |
| 0.005272762 | Generation of Precursor Metabolites | TCA cycle |
| 0.005213865 | Generation of Precursor Metabolites | TCA cycle |
